# The spatiotemporal distribution of human pathogens in ancient Eurasia and the emergence of zoonotic diseases

**DOI:** 10.1101/2023.10.06.561165

**Authors:** Martin Sikora, Elisabetta Canteri, Antonio Fernandez-Guerra, Nikolay Oskolkov, Rasmus Ågren, Lena Hansson, Evan K. Irving-Pease, Barbara Mühlemann, Sofie Holtsmark Nielsen, Gabriele Scorrano, Morten E. Allentoft, Frederik Valeur Seersholm, Hannes Schroeder, Charleen Gaunitz, Jesper Stenderup, Lasse Vinner, Terry C. Jones, Björn Nystedt, Karl-Göran Sjögren, Julian Parkhill, Lars Fugger, Fernando Racimo, Kristian Kristiansen, Astrid K. N. Iversen, Eske Willerslev

## Abstract

Infectious diseases have had devastating impacts on human populations throughout history, but important questions about their origins and past dynamics remain^1^. To create the first archaeogenetic-based spatiotemporal map of human pathogens, we screened shotgun sequencing data from 1,313 ancient humans covering 37,000 years of Eurasian history. We demonstrate the widespread presence of ancient bacterial, viral and parasite DNA, identifying 5,486 individual hits against 492 species from 136 genera. Among those hits, 3,384 involve known human pathogens^2^, many of which were detected for the first time in ancient human remains. Grouping the ancient microbial species according to their likely reservoir and type of transmission, we find that most groups are identified throughout the entire sampling period. Intriguingly, zoonotic pathogens are only detected ∼6,500 years ago, peaking ∼5,000 years ago, coinciding with the widespread domestication of livestock^3^. Our findings provide the first direct evidence that this lifestyle change resulted in an increased infectious disease burden. Importantly, they also suggest that the spread of these pathogens increased substantially during subsequent millenia, coinciding with the pastoralist migrations from the Eurasian Steppe^4,5^.

## Introduction

Pathogens have been a constant threat to human health throughout our evolutionary history. Until approximately 1850, at least a quarter of all children died before age one, and around another quarter before turning 15. Infectious diseases are estimated to have been responsible for over half of these deaths^6^. Larger disease outbreaks have profoundly impacted human societies, sometimes devastatingly affecting entire civilizations^7^. Infectious diseases have left lasting impressions on human genomes, as selective pressures from pathogens have continuously shaped human genetic variation^8–10^. Where and when different human pathogens first emerged, how and why they spread, and how they affected human populations are important but largely unresolved questions.

During the Holocene (beginning ∼12,000 years ago), the agricultural transition created larger and more sedentary communities, facilitating pathogen transmission and persistence within populations^11^. Simultaneously, the rise of animal husbandry and pastoralism are thought to have increased the risk of zoonoses^3^. Technological advances, such as horses and carts, increased both mobility and the risk of disease transmission between populations^12^. It has been hypothesised that these changes led to the so-called “first epidemiological transition” characterised by increased infectious disease mortality^3^.

However, direct evidence remains scarce and the idea is debated^13^. Paleopathological examinations of ancient skeletons offer insights into past infectious disease burden^14^, but are limited to few diseases identifiable from the available tissue. Recent advances in ancient DNA (aDNA) techniques allow for the retrieval of direct genomic evidence of past microbial infections, which can enable the reconstruction of complete ancient pathogen genomes. These studies have typically concentrated on specific pathogens and have provided surprising insights into the evolutionary history of the causative agents of some of the most historically important infectious diseases affecting humans, including plague (*Yersinia pestis*)^12,15–23^, tuberculosis (*Mycobacterium tuberculosis*)^24,25^, smallpox (*Variola virus*)^26,27^, Hepatitis B (*Hepatitis B virus*)^28–30^ and others^31–36^. However, there is an unmet need to investigate the combined landscape of ancient bacteria, viruses, and parasites that impacted our ancestors across various regions and time periods. Here we use a new high-throughput computational workflow to screen for ancient microbial DNA and use our data to investigate long-standing questi no ons in paleoepidemiology: When and where did important human pathogens arise? And what factors influenced their spatiotemporal distribution?

### Ancient microbial DNA in remains of 1,313 Eurasians

To understand the distribution of ancient pathogenic challenges, we developed an accurate and scalable workflow to identify ancient microbial DNA in shotgun-sequenced aDNA data (Extended Data Figs. 1-4; Supplementary information 1). The data (∼405 billion sequencing reads) derived from 1,313 ancient individuals from Western Eurasia (n=1,015; 77%), Central and North Asia (n=265; 20%) and Southeast Asia (n=33; 3%), spanning a ∼37,000 year period, from the Upper Paleolithic to historical times (Fig. 1b; Supplementary table S1; Supplementary information 2). As burial practices varied across cultures and time, these samples represent a subset of groups within past societies.

**Fig. 1.**
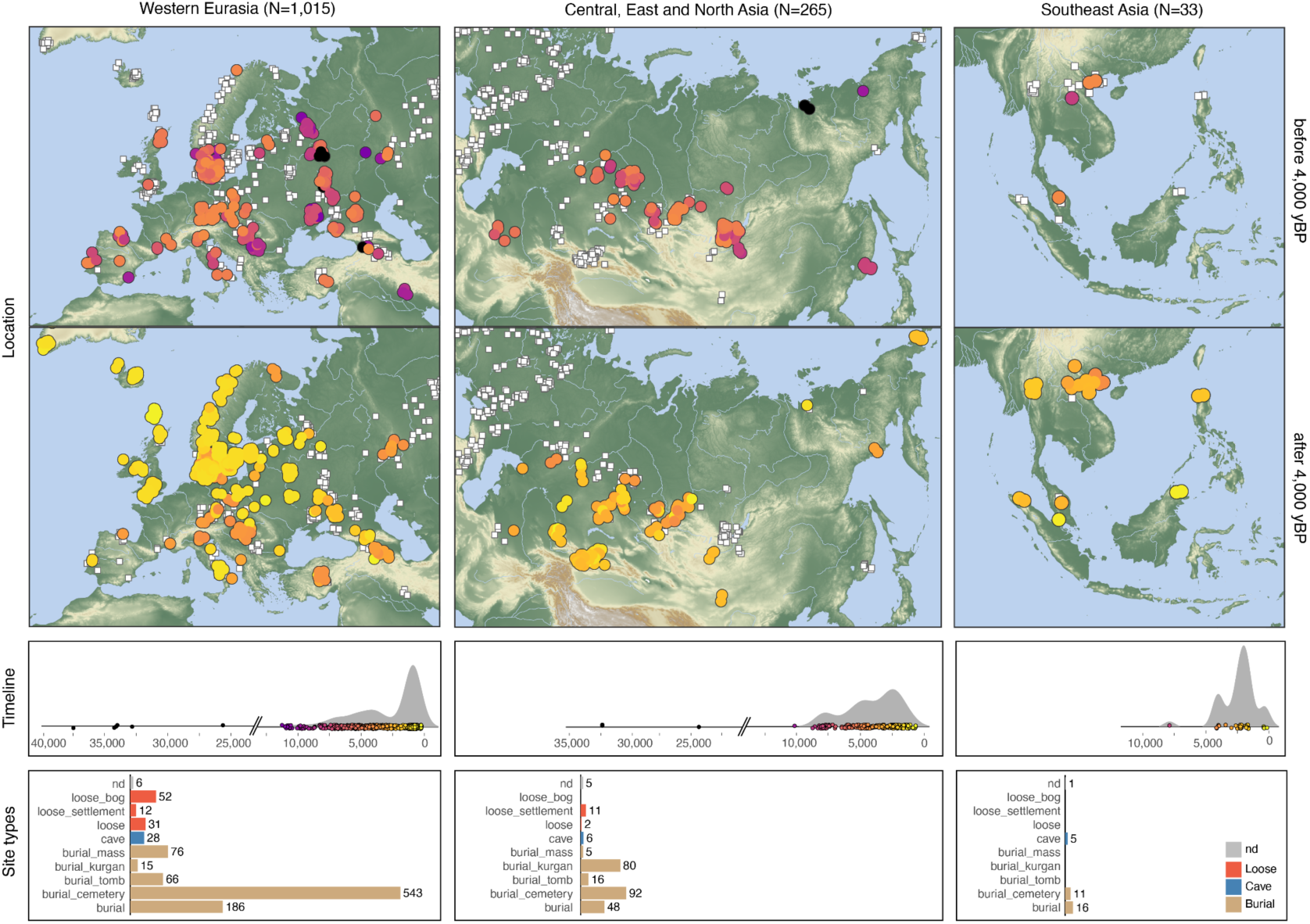
Dataset overview. Spatiotemporal distribution and site contexts of the study samples. White squares in the geographic maps indicate locations of the full set of 1,313 study samples, whereas coloured circles highlight location and age of samples from the time period and region indicated in the respective panel. Bar plots show numbers of samples for different site type contexts in each region (nd - not determined).

Nevertheless, the identified pathogens likely affected the broader population, as diseases spread easily in communities with poor sanitation and hygiene^37^. Initial metagenomic classification showed a large fraction of reads classified as soil-dwelling taxa including genera such as *Streptomyces* or *Pseudomonas,* reflecting a predominantly environmental source of microbial DNA. Further characterization using topic-model however suggested that microbial DNA in ancient tooth samples often derives from genera commonly associated with the human oral microbiome such *Actinomyces* or *Streptococcus* (Extended Data Fig. 1d-g).

We selected a set of 136 bacterial and protozoan genera (11,553 species total) containing human pathogenic species^2^ as well as 1,356 viral genera (259,979 species total) for further authentication and detection of ancient taxa. We found that ancient microbial DNA was widely detected, with 5,486 authenticated individual hits identified across 1,005 samples (Z-score for aDNA damage rate from *metaDMG* ≥ 1.5; Fig. 2a; Supplementary table S2; Extended Data Fig. 4). Of those, 3,384 hits were found among 214 known human pathogen species^2^, with the remaining 2,104 hits involving 278 other species. The highest numbers were observed in bacterial genera associated with the human oral microbiome, such as *Actinomyces* (380; 28.5% of samples) and *Streptococcus* (242; 18.1% of samples) or those commonly found in soil environments, such as *Clostridium* (252; 18.9% of samples) and Pseudomonas (111; 8.3% of samples).

**Fig. 2.**
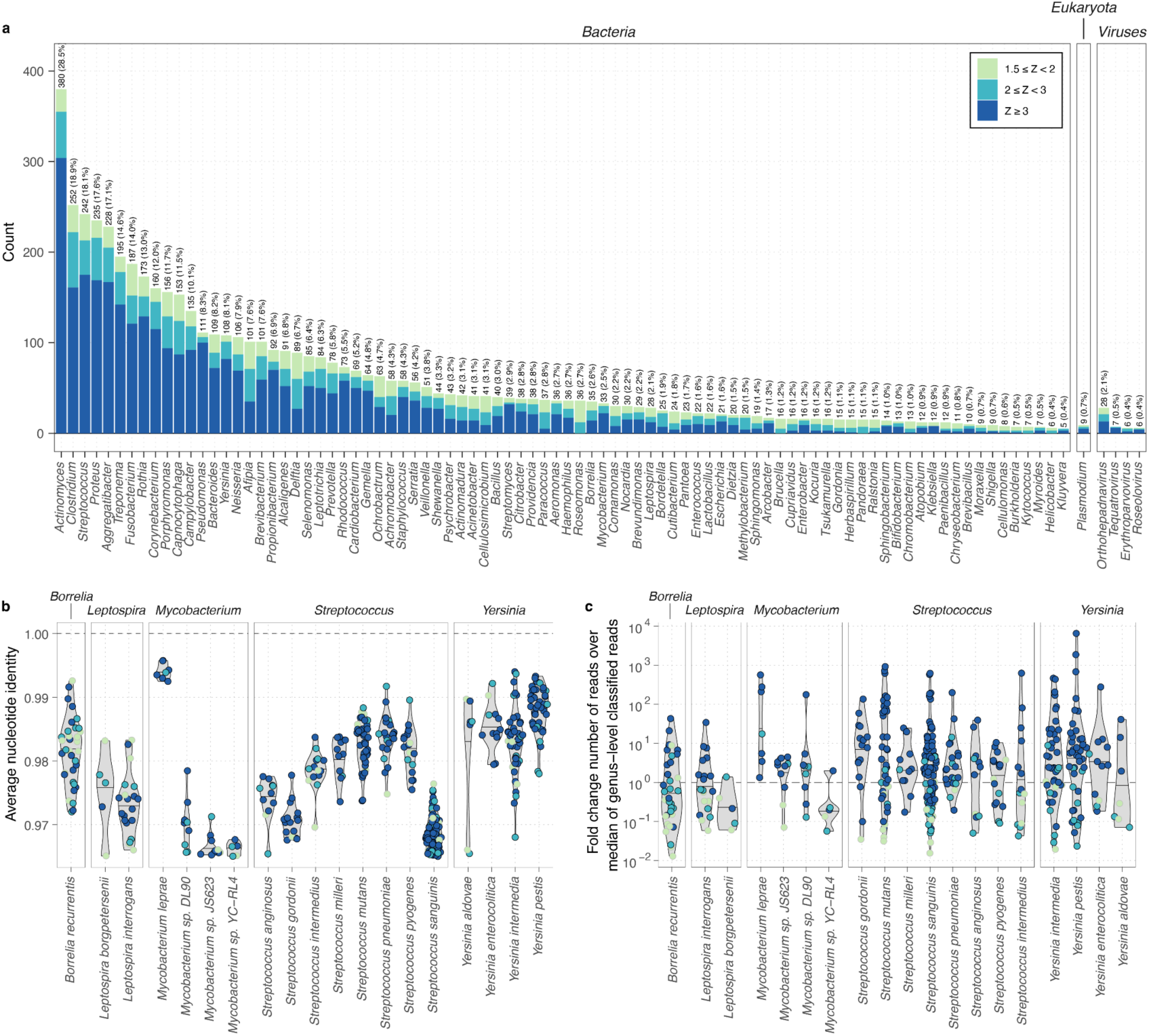
Overview and characteristics of detected ancient microbial DNA. **a,** Barplot showing total number of putative ancient microbial hits (overall detection rate in brackets) for bacterial, eukaryotic and viral (n ≥ 4) genera. Bar colour and shading distinguishes counts in the different aDNA damage categories. **b, c** Distributions of ANI (**b**) and log10-fold change of mapped reads over median of reads classified at taxonomic rank of genus per sample (**c**) for individual species hits in selected example genera. Symbol colour indicates aDNA damage category.

We observed marked differences in the distributions of the genetic similarity of the ancient microbial sequences to their reference assemblies, both among genera and between species within a genus (Fig. 2b; Extended Data Fig. 5).. High average nucleotide identity (ANI) indicates that ancient microbial sequences are closely related to a reference assembly in the modern database, and was observed in hits across all species from some genera (e.g. *Yersinia,* Fig. 2b). In other genera, only a few hits had a closely related database reference assembly match. An example is the genus *Mycobacterium,* where only hits of the leprosy-causing bacterium *Mycobacterium leprae* were highly similar to their reference assembly (ANI > 99%; Fig. 2b). Low ANI indicates that the ancient microbial DNA is only distantly related to the reference assembly, for example, due to aDNA damage, poor representation of the diversity of the genus in the database or false-positive classification of ancient microbial reads deriving from a related genus (Extended Data Fig. 3). Alternatively, ANI can also be reduced when reads mapped to a particular reference assembly originate from multiple closely related strains or species in a sample. To test for such mixtures, we quantified the rate of observing different alleles at two randomly sampled reads at nucleotide positions across the genomes of hits with read depth ≥1X. We found a high rate of multiple alleles in many species associated with the human oral microbiome, such as *Streptococcus sanguinis* or *Treponema denticola*. Hits for these species also showed lower ANI, consistent with the expectation for mixtures of ancient microbial DNA (Extended Data Fig. 6b, c).

The rate of read mapping varied by orders of magnitude between species, from hits in species with high read recruitment, such as *Mycobacterium leprae* (> 100-fold enrichment over the median number of classified reads across target genera) to hits at the lower limits of detection, e.g., for the louse-borne pathogen *Borrelia recurrentis* (lowest read recruitment ∼100-fold less than the median number of classified reads across target genera; Fig. 2c; Extended Data Fig. 5b). Ancient microbial DNA from species commonly found in soil, such as *Clostridium botulinum,* was detected at similar rates in tooth and bone samples. Conversely, species associated with the human oral microbiome (e.g., *Fusobacterium nucleatum, Streptococcus mutans* and *Porphyromonas gingivalis*) or pathogenic infections (e.g., *Yersinia pestis* and *Hepatitis B virus*) were significantly more frequently identified in tooth samples (Extended Data Fig. 6a). To further verify hits with low read numbers, we performed a BLASTn search for all reads of each hit with N ≤ 100 final reads (n=712 hits total; Supplementary table S3). Most hits showed a high proportion (≥ 80%) of reads assigned to the same species using BLASTn, and the species with the most top-ranked BLASTn hits generally matched the inferred hit species (Extended Data Fig. 7a, b).

Our results show that ancient microbial DNA isolated from human remains originates from complex mixtures of distinct endogenous and exogenous sources. The high detection rate, high read recruitment, lower ANI, and evidence of mixtures in genera such as *Clostridium* or *Pseudomonas* (Fig 2.; Extended Data Fig. 5, 6) suggest that a substantial fraction of this ancient microbial metagenome derives from environmental sources, possibly associated with the “necrobiome” involved in post-mortem putrefaction processes (Supplementary information 3)^38,39^. By contrast, species from other frequently observed genera, including *Actinomyces* or *Streptococcus,* were predominantly identified from teeth and likely originated from the endogenous oral microbiome^38^. Species representing likely cases of pathogenic infections (e.g., *Yersinia pestis* and *Mycobacterium leprae*) were often characterised by higher ANI and/or low multi-allele rate, consistent with pathogen load predominantly originating from a single dominant strain.

### The landscape of ancient pathogens across Eurasia

Our dataset provides a unique opportunity to investigate the origins and spatiotemporal distribution of major human pathogens in Eurasia, expanding the known range of some ancient pathogenic species and identifying others for the first time using paleogenomic data (Supplementary tables S3, S5).

Considering bacterial pathogens, we found widespread distribution of the plague-causing bacterium *Yersinia pestis*, consistent with previous studies^12,16,17,19,22,41,42^. We identified 42 putative cases of *Y. pestis* (35 newly reported; Extended Data Fig. 6e), corresponding to a detection rate of ∼3% in our samples. These newly identified cases expand the spatial and temporal extent of ancient plague over previous results (Fig. 3). The earliest three cases were dated between approximately 5,700-5,300 calibrated years before present (cal. BP), across a broad geographic area ranging from Western Russia (NEO168, 5,583-5,322 cal. BP), to Central Asia (BOT2016, 5,582-5,318 cal. BP), and to Lake Baikal in Siberia^43^ (DA342, 5,745-5,474 cal. BP). This broad range of detection among individuals pre- dating 5,000 cal. BP challenges previous interpretations that early plague strains represent only isolated zoonotic spillovers^20^. We replicated previously identified cases of plague in Late Neolithic and Bronze Age (LNBA) contexts across the Eurasian Steppe^16^ and identified many instances where multiple individuals from the same burial context were infected (Afanasievo Gora, Russia; Kytmanovo, Russia; Kapan, Armenia; Arban 1, Russia) (Supplementary table S2). These results indicate that the transmissibility and potential for local epidemic outbreaks for strains at those sites were likely higher than previously assumed^20^. Finally, 11 out of 42 cases were identified in late mediaeval and early modern period individuals (800-200 BP) from two cemeteries in Denmark (Aalborg, Randers), highlighting the high burden of plague during this time in Europe. All but one hit (NEO627, n=84 reads total) showed expected coverage for the virulence plasmids pCD1 and pMT1, with hits before 2,500 years BP characterized by the previously reported absence of a 19 kilobase region on pMT1 containing the *ymt* gene^16^ (Extended Data Fig. 7c; Supplementary information 4).

**Fig. 3.**
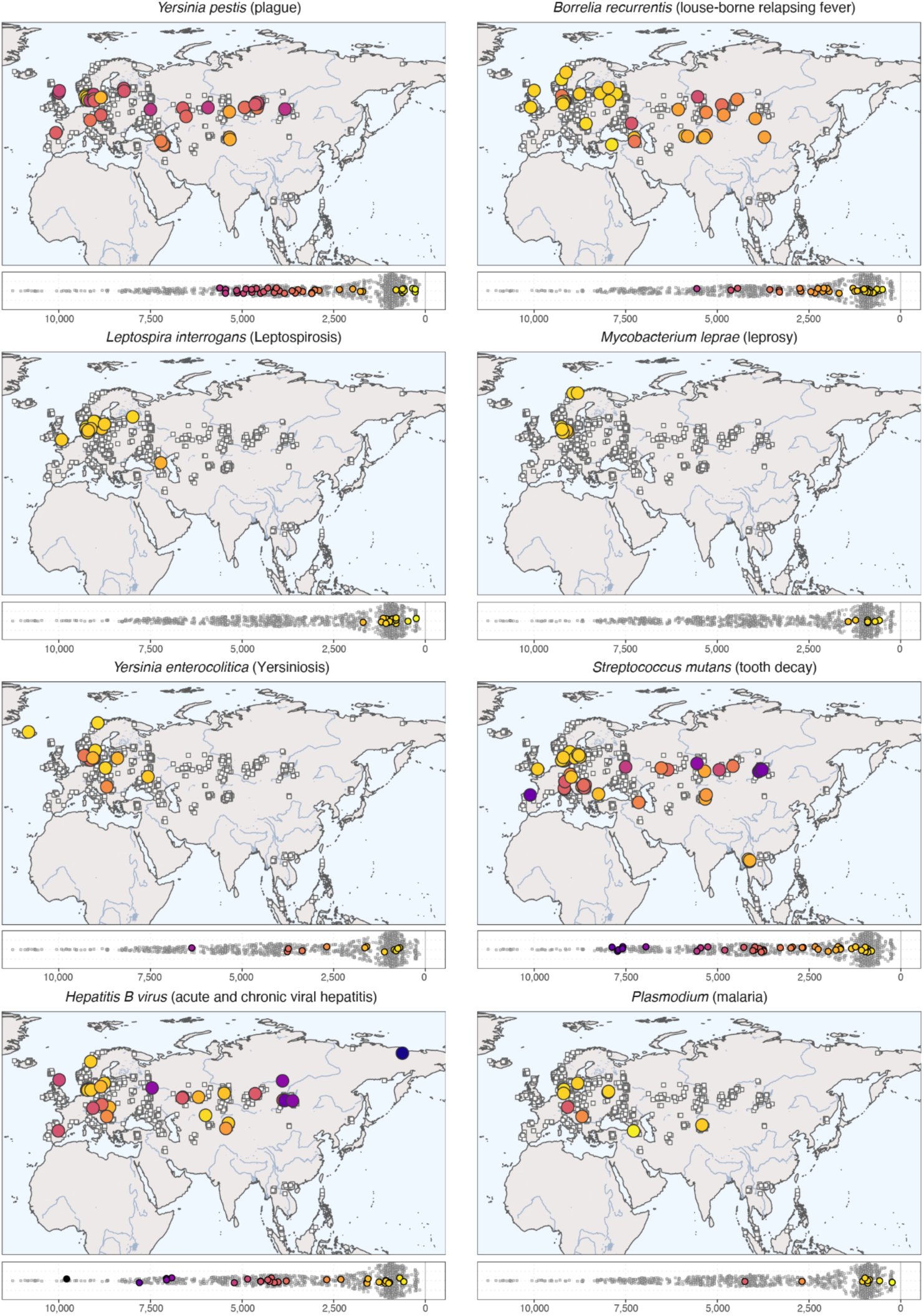
Spatiotemporal distribution of selected ancient pathogens. Each panel shows geographic distribution (top) and timeline (bottom) for identified cases of the respective pathogen (indicated by coloured circle). Geographic locations and age distributions of all 1,313 study samples are shown in each panel using white squares. The panel for *Plasmodium* combines the three species detected (*P. vivax* n=5; *P. malariae* n=3; *P. falciparum* n=1).

Another bacterial pathogen frequently detected was the spirochaete bacterium *Borrelia recurrentis*, causative agent of louse-borne relapsing fever (LBRF), a disease with a mortality of 10-40% (Supplementary information 5)^44^. While previous paleogenomic evidence for LBRF is limited to a few cases from Scandinavia and Britain^31,45^, we report 34 new putative cases (2.5% detection rate; Extended Data Fig. 6e), with wide geographic distribution across Europe, Central Asia, and Siberia (Fig. 3). We detected the earliest case in a Neolithic farmer individual from Scandinavia (NEO29, Lohals, 5,647-5,471 cal. BP), suggesting that human body lice were already vectors for infectious disease during the Neolithic period, supported by phylogenetic analyses of *B. recurrentis* in a recent preprint^45^. The highest detection rates were found during the Iron and Viking Ages. LBRF outbreaks were historically associated with crowded living conditions, poor personal hygiene, and wet and cold seasons, but are rare today in most regions (Supplementary information 5)^46^. Our results suggest that *B. recurrentis* infections exerted a substantial disease burden on past populations.

We also report novel cases of other bacterial pathogens previously detected in paleogenomic data. The leprosy-causing bacterium *Mycobacterium leprae* was identified in seven individuals (0.5% detection rate) from Scandinavia and only appeared from the Late Iron Age onwards (earliest case RISE174, 1,523-1,339 cal. BP). Because *M. leprae* can infect both red squirrels and humans^47^, and archaeological evidence demonstrates that fur trade from Scandinavia, including squirrel fur, increased substantially during the late Iron and Viking Ages^48^, our results support the suggestion that squirrel fur trade could have facilitated transmission^49^. Our findings are also consistent with the widespread distribution of leprosy in mediaeval Europe^50^. We further detected three putative cases of *Treponema pallidum* - subspecies of which are the causative agents of treponematoses such as yaws, and endemic and venereal syphilis - in three individuals from recent time periods (earliest case 101809T, Denmark, 600-500 BP; Fig. 3). Two cases were identified in individuals from Borneo in Southeast Asia (approximately 500-300 years BP); to our knowledge the first paleogenomic evidence for treponemal disease from this region.

Among the species reported for the first time using paleogenomic data, we identified twelve putative cases of *Yersinia enterocolitica,* the causative agent of yersiniosis, commonly contracted through consuming contaminated raw or undercooked meat (Fig. 3). The animal reservoirs for *Y. enterocolitica* include boars, deer, horses, cattle, and sheep. As *Y. enterocolitica* rarely enters the bloodstream, our results likely underestimate the disease burden. Interestingly, this species includes some of the only identified putative zoonotic infections in individuals from Mesolithic hunter-gatherer contexts (NEO941, Denmark, 6,446-6,302 cal. BP). We also detected other members of the order *Enterobacterales*, transmitted via the fecal-oral route, including members of the genera *Shigella*, *Salmonella*, and *Escherichia* (Supplementary table S2). We report the first evidence for ancient leptospirosis (genus *Leptospira*) dating back to the Neolithic, 5,650-5,477 cal. BP (NEO46, Sweden; *Leptospira borgpetersenii*). While earlier cases predominantly involved *Leptospira borgpetersenii* (n=5, 0.4% detection rate), the majority of hits were *Leptospira interrogans* (n=20, 1.5% detection rate), almost exclusively in Scandinavian contexts from the Viking Age onwards (Fig. 3). *Leptospira borgpetersenii* is today primarily found in cattle, while *Leptospira interrogans* is detected more broadly in both domestic and wild animals. Although the clinical manifestations are similar, with an untreated fatality rate of 1% today, transmission routes vary^51^. While host-to-host transmission predominates for *Leptospira borgpetersenii*, transmission via urine-contaminated environments dominates for *Leptospira interrogans* transmission. We also report two putative cases of *Corynebacterium diphtheriae*, the causative agent of diphtheria; the oldest of which dates back to the Mesolithic (Sidelkino, 11,336-11,181 cal. BP) (Supplementary table S2).

Other diseases associated with animals and livestock, such as listeriosis (*Listeria monocytogenes*) and brucellosis (genus *Brucella*), could not be reliably identified. Another major human pathogen not identified in our dataset is *Mycobacterium tuberculosis,* which causes tuberculosis (TB). However, as the *Mycobacterium tuberculosis* load in blood is typically low in immunocompetent patients without advanced disease^52^ and latent TB develops in 60% of cases and can persist for decades, it is, based on current knowledge, unlikely to be readily identified using aDNA data from tooth and bone remains sampled for ancient human DNA.

Identifying eukaryotic pathogens is challenging as sequence contamination from other organisms frequently occurs in their large and often fragmented reference genomes^53^. An illustrative example in our dataset is the protozoan parasite *Toxoplasma gondii*, which we readily identified in hits with high ANI and aDNA damage but low support from coverage evenness statistics, due to reads mapping to short contigs representing human contamination (Extended Data Fig. 4a,b; Supplementary Data 1).

Despite these challenges, we identified nine putative malaria infections across three different human-infecting species (*P. vivax* n=5; *P. malariae* n=3; *P. falciparum* n=1; Fig. 3; Supplementary table S2). The most widely detected parasite species was *P. vivax,* with the earliest evidence in a Bronze Age individual from Central Europe (RISE564, 4,750-3,750 BP based on archaeological context). Other cases include a mediaeval individual from Central Asia (DA204, Kazakhstan; 1,053-1,025 cal. BP) and two Viking Age individuals from Eastern Europe (VK224, 950-750 BP and VK253, 950-850 BP; Russia). The *P. vivax* malaria vector *Anopheles atroparvus* is currently widespread in Europe and nearby regions, including the Pontic Steppe, and our cases suggest this was also true in the past^54,55^. The single case of *P. falciparum* malaria was found in a sample from Armenia (NEO111; 463-0 cal. BP), where malaria was eliminated in the 1960s^56^.

Among DNA viral species, we found widespread infections with *Hepatitis B virus* (HBV; 28 cases, 2.1% detection rate), consistent with previous studies^28–30^ (Extended Data Fig. 6e). Our newly reported HBV cases include individuals from Mesolithic (Kolyma River, n=1) and Neolithic (Lake Baikal, n=3) contexts in Siberia dating back to 9,906-9,665 cal. BP, providing the first evidence for ancient HBV from those regions (Fig. 3). We also report the first putative ancient case (n=1) of *Torque teno virus* (TTV) dating back ∼7,000 years (NEO498, Ukraine; 7,161-6,950 cal. BP). TTV infects approximately 80% of the human population today, and while it is not associated with any particular disease, it replicates rapidly in immunocompromised individuals^57^. Other ancient virus hits included viruses not known to infect humans, such as ancient phage DNA (e.g,. *Escherichia phage T4, Proteus virus Isfahan;* Supplementary table S2) and one putative case of an ancient insect virus (*Invertebrate iridescent virus 31 (IIV-31))* in a tooth sample of a Viking Age individual from Sweden (VK30, Varnhem; 950-650 BP)^58^. The virus source is likely exogenous, potentially originating from aDNA of food sources in the tooth remains.

Co-infections with multiple pathogens can worsen disease progression and outcomes^59^ and they were likely an important morbidity factor in ancient human populations. Searching for individuals showing co-occurrence of distinct ancient microbial species, we identified 15 cases of putative co-infections in our dataset (Supplementary table S2). A striking case was a Viking Age individual from Norway (VK388), where we replicated previous results of infection with a likely smallpox-causing variola virus^27^ and additionally found evidence of infection with the leprosy-causing bacterium *Mycobacterium leprae.* Another case of possible co-infection with *Mycobacterium leprae* was found in VK366, a Viking Age individual from Denmark, who also showed evidence for leptospirosis (*Leptospira interrogan*s). Interestingly, among the 15 cases, six involved co-infections of HBV with non-viral pathogens (*Yersinia pestis* n=3; *Borrelia recurrentis* n=2; *Plasmodium malariae* n=1; Supplementary table S2). This suggests that some of these cases involved chronic hepatitis, possibly reflecting HBV infection during infancy, when hepatitis becomes chronic in 90-95% of modern cases, compared to only 2-6% in adult infections. An intriguing early case of a possible co-infection was found in a Mesolithic hunter-gatherer from Russia (Sidelkino, 11,336-11,181 cal. BP). This individual showed evidence of the respiratory pathogen *Corynebacterium diphtheriae*, and *Helicobacter pylori*, usually restricted to gastric infections; however, rare contemporary examples of bacteremia have been reported for both^60,61^. Overall, our results show that co-infections can be detected using ancient metagenomic screening, but are likely underestimated given methodological limitations such as differences in pathogen load, tissue availability, and other factors impacting detectability of ancient microbial DNA.

### Temporal dynamics and drivers of epidemic pathogens

Understanding the factors affecting the dynamics of past epidemics is a major aim of paleoepidemiology. Our dataset allows us to address this question using direct molecular evidence for ancient pathogens across prehistory. To investigate changes in pathogen incidence over time, we performed Bayesian change-point detection and time series decomposition^62^ on two pathogens with high detection rates, *Yersinia pestis* (plague) and *Borrelia recurrentis* (LBRF), using the detection rate of the respective pathogen as a proxy for its incidence (Methods). For plague, we inferred a gradual rise in detection rate starting from ∼6,000 BP, about 1,000 years after the estimated time to the most recent common ancestor of currently known ancient strains (7,100 cal. BP)^20^. It reached a first peak around ∼5,000 BP across Europe and the Eurasian Steppe, coinciding with the emergence and early spread of the LNBA- strains, believed to have had limited flea-borne transmissibility^16,17,22^ (Fig. 4). Detection remained high with additional peaks for a ∼3,000 year period, until an abrupt change ∼2,800 BP led to a ∼800 year period where plague was only detected in one sample (VK522, Oland, Sweden 2,343-2,154 cal. BP). Starting at ∼2,000 BP, plague reappeared in three samples from Central Asia (DA92, DA101, DA104, Kazakhstan and Kyrgyzstan; Fig. 3; Supplementary table S2), just before the first historically documented plague pandemic (Fig. 4). Another hiatus of ∼600 years led to a rise and peak associated with the second plague pandemic ∼600 BP (European late mediaeval cases, Denmark and previously published cases^63^; Fig. 4). This pattern of change coincides with the extinction of the LNBA- strains ∼2,700 BP^22^ and the second *Yersinia pestis* diversification event starting ∼3,700 BP, which gave rise to an extinct Bronze Age lineage (RT5, LNBA+)^18^ and present- day lineages; these had increased flea-mediated transmission adaptations favouring bubonic plague and led to all known later plague pandemics^64^. The adaptations included acquiring two plasmids: one with the *ymt* gene for survival in the flea midgut and another with the *pla* gene for invasiveness after transmission^65^. The lack of detection during both periods is also seen in publicly available ancient *Yersinia pestis* genomes from other Eurasian sites^63^, suggesting that sampling bias is unlikely to significantly influence the observed dynamics.

**Fig. 4.**
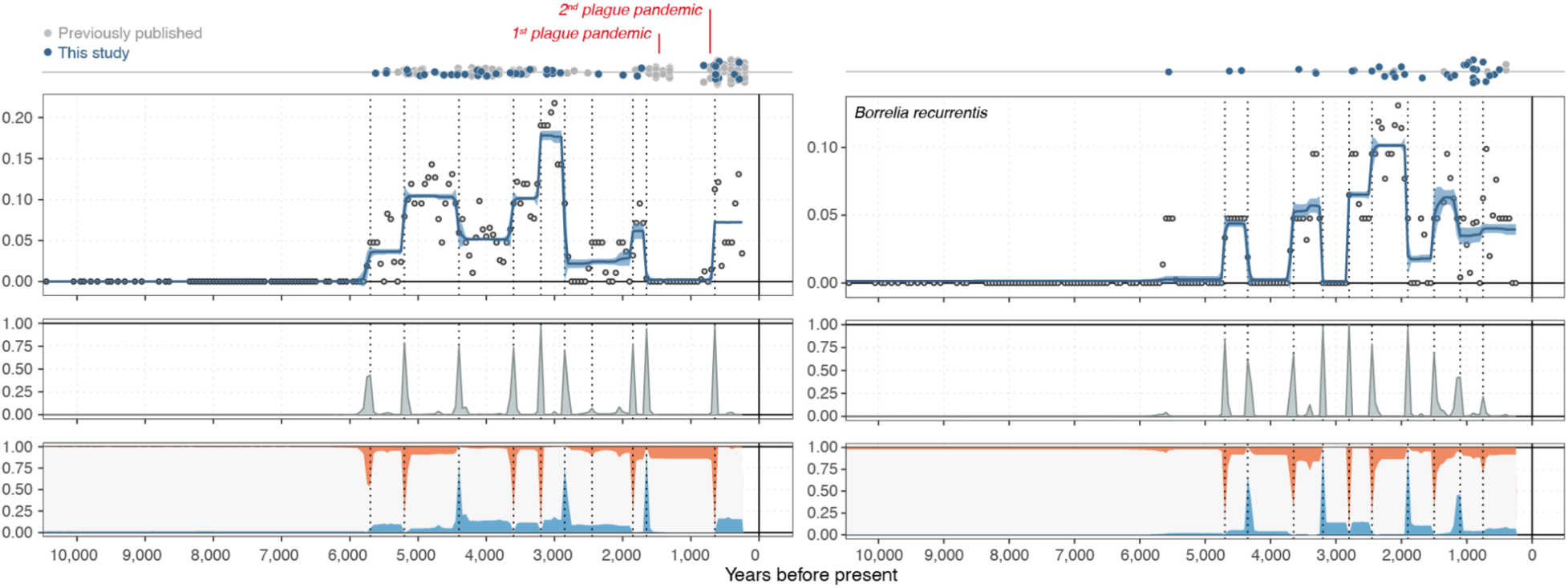
Bayesian time series decomposition of major epidemic pathogens. Panels show estimated trendlines and 95% credible interval for detection rates (top), probability distributions and locations (dotted lines) for change points (middle) and probability of trend slope (bottom) being positive (red), negative (blue) or zero (white), inferred using Bayesian change-point detection and time series decomposition. Top of panels show temporal distributions of newly reported pathogen hits (blue circles) as well as previously published ancient pathogens (grey circles) from the respective species.

The inferred temporal dynamics of LBRF show a first peak in detection around 5,500 BP, slightly more recent than for plague, but with more sporadic occurrences and sharper peaks during the first ∼2,000 years (Fig. 4). The geographic extent during the early period ranges from Scandinavia (NEO29, Denmark, 5,647-5,471 cal. BP) to the Altai mountains (RISE503, Russia, 3,677-3,461 cal. BP) (Fig. 3; Supplementary table S2). From ∼2,800 BP, LBRF was detected more consistently, peaking approximately 2,000 years ago, predominantly in the Eurasian Steppe region (Fig. 3). This change from epidemic outbreaks to endemicity overlaps in time with the estimated emergence of a distinct *Borrelia recurrentis* Iron Age clade^45^ (Supplementary information 5). The period of high LBRF detection coincided with a time without detectable plague activity (Fig. 4), reinforcing that the absence of plague is not due to sample size limitations or poor DNA preservation. This opposing pattern is unlikely to result from any cross-immunity between *Yersinia pestis* and *Borrelia recurrentis* but could plausibly, in part, be caused by population size decreases and behavioural and societal adjustments during plague epidemics. LBRF remained detectable until the end of the time series, particularly in Europe; the continued presence might have facilitated the emergence of a Medieval *B. recurrentis* clade ∼600 years BP^45^ (Supplementary information 5) (Fig. 3, 4).

A striking feature shared in the temporal dynamics of plague and LBRF was the absence of detectable cases before ∼6,000 BP, coinciding with a transition of individuals in predominantly hunter-gatherer contexts to those in farming or pastoralist cultural contexts. It has been hypothesized that this transition led to a higher risk of zoonotic disease transmission, and facilitated the spread of both old and new pathogens^3^. Our dataset allows us to test this hypothesis using molecular evidence for infectious disease burden. To increase power to detect changes in the load of different pathogen types, we focused on grouped ancient microbial hit categories (Supplementary table S4).

We found that species associated with the ancient oral microbiome showed the highest relative detection rate, accounting for up to 50% of ancient hits across various periods (Fig. 5a; Extended Data Fig. 8). Species in the “environmental” classes of likely exogenous origins were also detected at consistent rates throughout time. Species in the “infection” classes occurred at low detection rates throughout (mostly <10%). Strikingly, we found that species in the “zoonotic” reservoir classes were not detected until approximately 6,500 BP (Fig. 5a). Using Bayesian time series decomposition^62^, we inferred an overall increase in the detection rates of the “zoonotic” reservoir classes from ∼6,000 BP, thereafter remaining at elevated levels until the mediaeval period (Fig. 5b; Extended Data Fig. 8, 9a). While species in the “anthroponotic” reservoir classes also occur earlier (predominantly species with human-to-human transmission, Extended Data Fig. 9a), we observe increased detection rates from ∼2,500 BP onwards (Fig 5b, Extended Data Fig. 8). Our results provide the first direct evidence for an epidemiological transition of increased infectious disease burden after the onset of agriculture through to historical times.

**Fig. 5.**
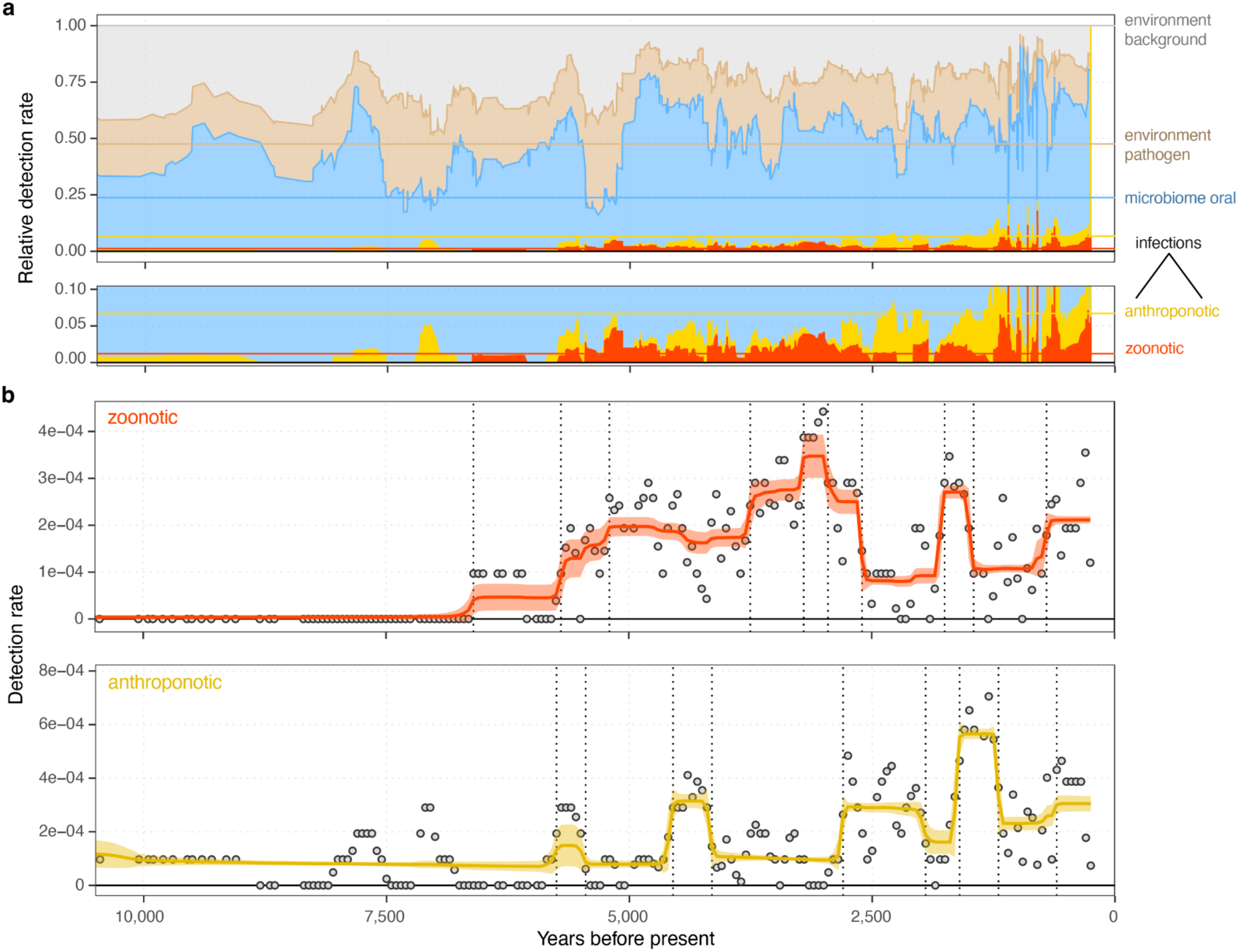
Time series of ancient microbes by microbial source. **a,** Timeline of relative detection rates in sliding windows of 21 temporally consecutive samples, for different ancient microbial species classes. Coloured horizontal lines indicate the expected rates if species in all classes would be detected at equal rates, based on the total number of distinct species in each class. **b,** Trendlines for detection rates inferred using Bayesian change-point detection and time series decomposition, for ancient microbial species in the “zoonotic” (top) and “anthroponotic” (bottom) reservoir class.

We used Bayesian spatiotemporal modelling^67^ to investigate possible drivers of the observed ancient microbial incidences. We modelled the presence/absence of either individual microbial species or combined species groups using sets of putative covariates, including spatiotemporal variables (longitude, latitude, and sample age), paleoclimatic variables (mean annual temperature and precipitation), human mobility and ancestry, sample material (tooth or other), and a proxy for “detectability” (number of human-classified reads). In the models for the “zoonotic” or “anthroponotic” infection species classes, sample age was an important predictor (Fig. 6; Extended Data Fig. 10; Supplementary table S6), consistently negatively associated with incidence, and high effect sizes in the individual species models for *Borrelia recurrentis* and *Leptospira interrogans* (Fig. 6, Extended Data Fig. 10). Longitude was another important factor in the “infection” classes; it was positively associated with incidence rates for the combined “anthroponotic” class, and in individual models for *Yersinia pestis* and *Borrelia recurrentis*. The positive effect of longitude suggests a higher incidence in the eastern part of our spatiotemporal range, where samples from the Eurasian Steppe predominate.

**Fig. 6.**
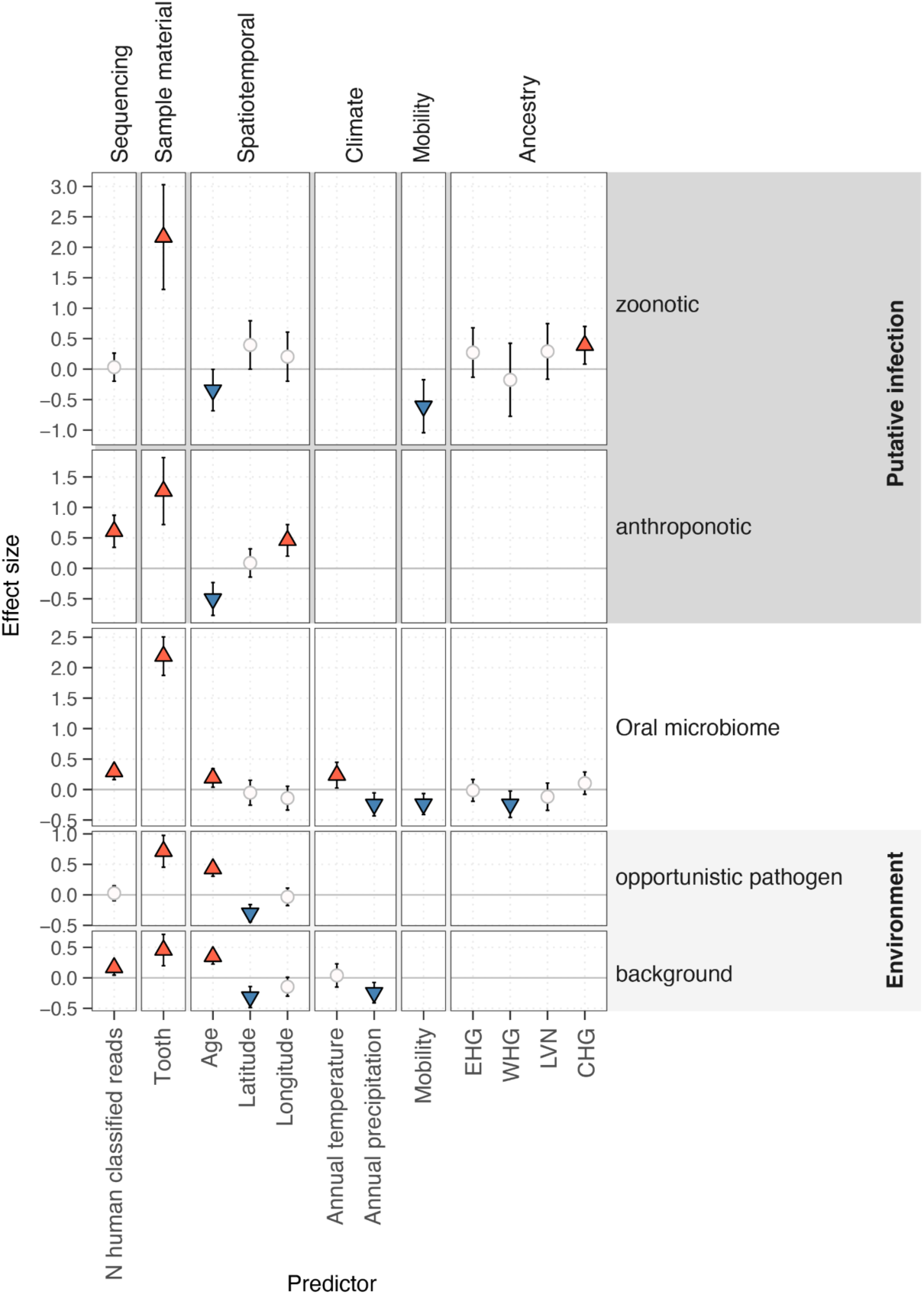
Predictors of ancient microbial species incidence. Matrix showing effect sizes and of 12 potential predictors (columns) for presence of selected combined ancient microbial species groups inferred from spatiotemporal modelling. For each class, the model with lowest Watanabe–Akaike information criterion is shown. Symbols indicate the predictors included in the respective model. Predictors with positive effect (2.5% and 97.5% posterior quantiles both positive) are shown as red triangles, whereas predictors with negative effect (2.5% and 97.5% posterior quantiles both negative) are shown as blue inverted triangles. Predictors included in the best-fitting model but without effect (posterior quantile range spanning zero) are indicated using white circles. Posterior standard error of effect sizes is indicated by error bars.

The increased infection incidence in Steppe populations could reflect an increased genetic susceptibility or a higher risk of acquiring diseases associated with the pastoralist lifestyle. The latter suggestion seems more plausible as continued exposure to selective pressures from certain infectious diseases likely would reduce susceptibility in these populations. Human ancestry showed small but consistent positive effects in some models, particularly the infection classes, for the Caucasus hunter-gatherers (CHG). Across all models, the incidence of ancient microbes was positively associated with teeth as sample material; the highest effect sizes were found in the "oral microbiome" and "infection" classes (Fig. 6, Extended Data Fig. 10). Teeth preserved ancient oral microbiome and pathogen DNA better than petrous bones (the source of 86% of our samples), likely due to oral cavity exposure and better access to microbial DNA in the bloodstream^68^. These results support the notion that species detected in those classes are predominantly of endogenous origin.

## Conclusions

During the Holocene, human lifestyles changed significantly as agriculture, animal husbandry, and pastoralism became key practices but the impact on infectious disease incidence is debated. Our study represents the first large-scale characterization of ancient pathogens across Eurasia, providing clear evidence that identifiable zoonotic pathogens emerged around 6,500 years ago and were consistently detected after 6,000 years ago. While zoonotic cases likely existed before 6,500 years ago, the risk and extent of zoonotic transmission probably increased with the widespread adoption of husbandry practices and pastoralism. Today, zoonoses account for over 60% of newly emerging infectious diseases^70^.

Strikingly, we observed some of the highest detection rates at ∼5,000 BP, a time of significant demographic changes in Europe due to the migration of Steppe pastoralists and the displacement of earlier populations^4,5^. Steppe pastoralists, through their long-term continuous exposure to animals, likely developed some immunity to certain zoonoses and their dispersals may have carried these diseases westward and eastwards. Consequently, the genetic upheaval in Europe could have been facilitated by epidemic waves of zoonotic diseases causing population declines, with depopulated areas subsequently being repopulated by opportunistic settlers who intermixed with the remaining original population. This scenario would mirror the population decline of Indigenous people in the Americas following their exposure to diseases introduced by European colonists^71,72^. Our findings support the interpretation of increased pathogen pressure as a likely driver of positive selection on immune genes associated with the risk of multiple sclerosis in Steppe populations ∼5,000 years ago^73^, and immune gene adaptations having occurred predominantly after the onset of the Bronze Age in Europe^10^.

Expanding our analyses to the broader pathogen landscape allowed us to infer and contrast incidence patterns between different species and types of pathogens to a greater extent than previously possible. If ancient pathogen DNA of a single species is not detected in a particular region or period, asserting whether this is due to low disease incidence or confounding factors such as differential DNA preservation between different periods and environments is challenging. Our analyses counter these limitations; we demonstrate that pathogens with known epidemic potential and high detection rates, such as *Yersinia pestis* (plague) and *Borrelia recurrentis* (LBRF), show striking differences in their detection rate over time, suggesting that low detection rate in these cases represent an actual reduction in incidence. During the early period (∼5,700-2,700 years ago), the continuous detection of *Yersinia pestis* is suggestive of endemic disease. The succeeding pattern of distinct waves and periods without detection indicate epidemic outbreaks; these detection peaks match the historically described plague pandemics. This shift from endemic to epidemic is concurrent with significant changes in the *Yersinia pesti*s genome, particularly increased flea-transmissibility and pathogenicity^16,18^. The pattern for *Borrelia recurrentis* is almost entirely the opposite, with narrow peaks and long periods without detection, suggesting local epidemics before ∼2,700 years ago and consistent detection afterwards.

This later endemicity of LBRF could be driven by changes in the bacterial genome and by human and environmental factors known to increase the risk of louse infestation^45,74,66^. Experimental studies have demonstrated that *Yersinia pestis,* like *Borrelia recurrentis,* can infect body lice in the midgut, and sometimes, also the Pawlowsky glands (PG), a putative salivary gland^66^. Body lice infected in the PG can transmit *Yersinia pestis* in concentrations sufficient to initiate disease in humans, possibly contributing to transmission during plague outbreaks. Infected body lice have higher mortality than uninfected lice, and it remains unknown whether co-infection of body lice with *Yersinia pestis* and *Borrelia recurrentis* is possible.

Our study has some important limitations. While ancient shotgun metagenomic data offers direct evidence of past infections, its usefulness depends on having a high pathogen load and the right tissue samples. Our ancient tooth and bone samples are well suited to detect high-load bloodstream infections like *Yersinia pestis* and *Borrelia recurrentis*, but pathogens with lower loads or different tissue preferences are underrepresented. Moreover, differentiating ancient infections from those arising from environmental sources, like the "necrobiome," is challenging. Finally, our dataset lacks information on RNA viruses, therefore underestimating the zoonotic disease burden. However, the timing is probably accurate as the conditions favouring zoonotic transmission of RNA viruses are similar to those of other zoonotic pathogens^69^.

Our findings represent the first example of how the nascent field of genomic paleoepidemiology can create a map of the spatial and temporal distribution of diverse human pathogens over millennia. This map will develop as more ancient specimens are investigated, as will our abilities to match their distribution with genetic, archaeological, and environmental data. Our current map shows clear evidence that lifestyle changes in the Holocene led to an epidemiological transition, resulting in a greater burden of zoonotic infectious diseases. This transition profoundly affected human health and history throughout the millennia and continues to do so today.

## Methods

### Dataset

We compiled a dataset of aDNA shotgun-sequencing data from 1,313 ancient individuals previously sequenced for studies of human population history (references for previous publications describing laboratory procedures and sample/site descriptions in Supplementary table S1). To facilitate ancient microbial DNA authentication, we excluded sequencing libraries subjected to UDG treatment which removes characteristic aDNA damage patterns from further analyses. Samples sequenced across multiple libraries were combined into single analysis units to maximise sensitivity for detection of ancient microbial DNA present in low abundance.

### Ancient microbial DNA screening

We carried out screening for ancient microbial DNA using a computational workflow combining *k*- mer-based taxonomic classification, read mapping and aDNA authentication. We first performed taxonomic classification of the sequencing reads (minimum read length 30 bp) using *KrakenUniq*^75^, against a comprehensive database of complete bacterial, archaeal, viral, protozoan genomes in the RefSeq database (built with default parameters of *k*-mer size 31 and low-complexity sequences masked). To increase sensitivity for ancient viral DNA, we re-ran the classification on a viral-specific database of complete viral genomes and neighbour assemblies from RefSeq (https://www.ncbi.nlm.nih.gov/genome/viruses/about/assemblies/), using all reads classified as non-human from the previous run.

Following this initial metagenomic classification, a subset of genera was further processed in the genus-level read mapping and authentication stages. For bacterial pathogens, we selected genera with two or more established species of human pathogens from a recent survey of human bacterial pathogens^2^ (n=125 genera). Genera with a single pathogenic species were not included in order to balance between including genera responsible for substantial human pathogenic burden and computational feasibility. We further included genera including human protozoan pathogens (n=11 genera), as well as all viral genera (n=1,356).

For each genus of interest showing ≥ 50 unique *k*-mers assigned, all sequencing reads classified were collected and aligned in parallel against a representative reference assembly for each individual species within the genus. We selected the assembly with the most unique *k*-mers assigned as the representative reference genome for each species in a particular sample. If no reads were assigned to any assembly of the species in *KrakenUniq*, we selected the first assembly for mapping. Read mapping against the selected assembly was carried out using *bowtie2*^76^, using the ‘very sensitive’ preset and allowing one mismatch in the seed (’-N 1’ option). Mapped BAM files were subjected to duplicate marking using ‘*samtools markdup’*^77^ , and filtered for mapping quality MAPQ≥20. aDNA damage rates were estimated using *metaDMG*^78^.

### Authentication of ancient microbial DNA

To authenticate ancient microbial DNA, we calculated sets of summary statistics quantifying expected molecular characteristics of true positive ancient microbial DNA hits^79^:

### Similarity to the reference assembly

Summary statistics in this category measure how similar sequencing reads are to a particular reference assembly, with true positive hits expected to show higher similarity than false positive hits. Summary statistics used include:

#### Average edit distance

The average number of mismatches in sequencing reads mapped to a particular reference (lower - more similar to reference).

#### Average nuclear identity (ANI)

The average number of bases in a mapped sequencing read matching the reference assembly, normalised by the read length (higher - more similar to reference).

#### Number of unique *k*-mers assigned

The number of unique *k-*mers assigned to a particular reference assembly from *KrakenUniq* classification (higher - more similar to reference).

### Ancient DNA characteristics

Summary statistics in this category measure the evidence for sequencing reads deriving from an aDNA source. Summary statistics used include:

#### Average read length

The average length in base pairs of sequencing reads mapped to a particular reference (shorter - more likely ancient).

#### Terminal aDNA substitution rates

The frequency of C>T (G>A) substitutions observed at the 5’ (3’) terminal base across all sequencing reads mapped to a particular reference (higher - more likely ancient).

#### Bayesian D_max_

Bayesian estimator of aDNA damage rate from *metaDMG* (higher - more likely ancient).

#### Bayesian Z_x_

Bayesian estimator of significance of evidence for aDNA damage rate from *metaDMG* (higher - more likely ancient).

### Evenness of genomic coverage

Summary statistics in this category measure how evenly mapped sequencing reads are distributed across a reference assembly. Summary statistics used include:

#### Average read depth

The average number of reads covering a base in the reference assembly.

#### Breadth of coverage

The fraction of the reference assembly that is covered by one or more sequencing reads.

#### Expected breadth of coverage

Breadth of coverage expected for a particular average read depth, calculated^80^ as 1 - e-^(average read depth)^

#### Ratio of observed over expected breadth of coverage

The ratio of breadth of coverage observed in mapping over breadth of coverage expected given observed average read depth (higher - more even coverage).

#### Relative entropy of read start positions

A measure for the information content of the genomic positions of mapped reads. To obtain this statistic, we calculate the frequency of read alignments with their start positions falling within windows along the reference assembly, using two different window sizes (100bp and 1000bp). The obtained frequency vector is converted into Shannon information entropy, and normalised using the maximum entropy attainable if the same total number of reads were evenly distributed across the windows (higher - more even coverage).

### Filtering of putative ancient microbial hits

From this initial screening, we then selected a subset of putative microbial “hits” (sample/species combinations) for further downstream analysis based on a set of aDNA authentication summary statistics:

- Number of mapped reads ≥ 20

- 5’ C>T deamination rate ≥ 0.01

- 3’ G>A deamination rate ≥ 0.01

- Ratio of observed/expected breadth of coverage ≥ 0.8

- Relative entropy of read start positions ≥ 0.9

- ANI > 0.965

- Rank of number of unique *k*-mers assigned ≤ 2

For this initial filtered list of putative microbial hits, we ran *metaDMG* using the full Bayesian inference method to obtain Z-scores measuring the strength of evidence for observing aDNA damage (Supplementary Data 2).

The final list of putative individual ancient microbial hits was then obtained using the filtering cutoffs

- *metaDMG* Bayesian D_max_ ≥ 0.05

- *metaDMG* Bayesian Z ≥ 1.5

- Rank of number of unique *k*-mers assigned == 1

For authentication of viral species, we used the same filtering cutoffs described above, except for a lower ANI cutoff (> 0.95), as well as a lower cutoff for relative entropy of read start positions (>0.7) for short viral genomes (< 10kb).

The result of this filtering is a single best-matching species hit for each sample and genus of interest Supplementary table S2. We note that this approach will miss potential cases where aDNA from multiple species of the same genus are present in the sample. However, due to the considerable challenges involved in distinguishing this scenario from false positives due to cross-mapping of ancient reads from a single source of DNA to reference assemblies of a closely related species (e.g., *Yersinia pestis / Yersinia pseudotuberculosis*), we opted for the conservative option of retaining only the best hit for each genus.

To further authenticate putative hits with low read counts (N ≤ 100 final reads), we carried out a BLASTn analysis. We extracted the reads for a species hit from the final filtered BAM files, and queried them against the ‘nt’ database (downloaded 20240828) using ‘blastn -task blastn -max_hsps 1’. For the reads of each putative ancient microbial hit, we then tabulated the number of times and proportion of the highest scoring BLAST hits matched either the genus or species inferred from our workflow Supplementary table S3).

### Simulations of ancient microbial DNA

We simulated aDNA fragments from microbial reference genomes *in silico* using *gargammel*^81^. We chose nine species representing pathogens of interest, and for each selected an assembly not present in the pathogen screening workflow database:

- *Brucella melitensis* (GCF_027625455.1)

- *Helicobacter pylori* (NZ_CP134396.1)

- *Mycobacterium tuberculosis* (NZ_CP097110.1)

- *Salmonella enterica* (NZ_CP103966.1)

- *Yersinia pestis* (NZ_CP064125.2)

- *Yersinia pseudotuberculosis* (NZ_CP130901.1)

- *Plasmodium vivax* (GCA_900093555.2)

- *Variola virus* (GCA_037113635.1)

- *Human betaherpesvirus 5* (GCA_027927465.1)

For each reference genome, we simulated 5 million single-end sequencing reads (100 bp read length) with adapter sequences, with read length distribution and damage patterns from a *mapDamage2* results of a previously published ancient pathogen genome (RISE509, *Yersinia pestis*^16^). The full-length simulated reads were then adapter-trimmed using *AdapterRemoval*^82^. To investigate the ability of the workflow to detect low abundance ancient microbes, we randomly down-sampled the full read set for each reference genome using *seqtk* (https://github.com/lh3/seqtk) (50, 100, 200, 500 reads; 10 replicates each).

### Topic model analysis

We carried out topic model analysis on taxonomic classification profiles for each sample using the R package *fastTopics*^83^ (https://github.com/stephenslab/fastTopics). We used the number of unique *k*- mers assigned to non-human genera from *KrakenUniq* as the observed count data for each sample, excluding genera with less than 50 unique *k*-mers assigned. The analysis was carried out for L=2 and L=3 topics, to capture broad structure in the classification profiles.

### Ancient microbial groups

For combined analyses, we grouped the ancient microbial hits into three categories, based on the likely source of the microbial DNA (Supplementary table S4):

1. Environmental, to capture all hits derived from environmental sources including the necrobiome (labelled environment_background, environment_pathogen, to distinguish potential pathogenic species from non-pathogenic ones);
2. Oral microbiome, including both commensal and pathogenic species (microbiome_oral)
3. likely pathogenic infections, further distinguished into different modes of transmission (infection_anthroponotic; infection_vector_borne; infection_zoonotic).

We define zoonotic pathogens here as those transmitted from animals to humans or which made such a host jump in our sampling time frame^40^.

### Time series

To infer temporal dynamics of ancient microbial species, we calculated detection rates in a sliding window of k=21 temporally consecutive samples across the entire timeline of the 1,266 samples with dating information. For individual species, the detection rate for each window corresponds to the proportion of the 21 samples in each window that were positive for the species of interest. For analyses of species combined in classes, we calculated the detection rate as the ratio of the total number of hits within a class in the window over the total number of possible hits across all species in a window (21 samples x 258 species across all classes). For individual species with n ≥ 20 hits or combined species classes, we further performed Bayesian change-point detection and time series decomposition (BEAST) ^62^ implemented in the R package *Rbeast* (https://github.com/zhaokg/Rbeast), using the detection frequencies described above as response variables.

### Spatiotemporal models of species incidence

To identify possible drivers of the observed spatiotemporal ancient microbial incidence, we combined the individual microbial species and the combined species groups with palaeoclimatic variables, human mobility estimates and kriged estimates of ancestry composition for Holocene West Eurasia. Palaeoclimatic reconstructions were accessed using the CHELSA-Trace21k data, which provides global monthly climatologies for temperature and precipitation at 30 arcsec spatial resolution in 100- year time steps for the last 21,000 years^84^. To pair the microbial species/groups to the palaeoclimatic reconstructions, we took the average climatic value across all the time steps that fall within the microbial species/group age ± sd at each of the sampling locations. Palaeoclimatic variables considered were annual mean temperature (BIO01) and annual precipitation (BIO12). Human mobility values were accessed from Schmid & Schiffels^85^ and approximately represent the distance in kilometres between the burial location of the ancient human individual and its putative ancestral origin, based on patterns of genetic similarity derived from a MDS analysis. Microbial species/groups were paired to the mobility estimate of the ancient human individual that occurs closest in space and time. Kriged ancestry estimates were extracted from Allentoft *et al.*^86^, using the spatiotemporal ancestry kriging method from Racimo et al.^87^, and paired to the closest spatiotemporal location of the ancient human remain where the corresponding microbial species/groups were sampled.

To determine the influence of the covariates on the microbial incidence, we used a hierarchical Bayesian model implemented in the *inlabru* R package^67,88^, where ancient microbial presence/absence follows a binomial distribution and the spatiotemporal variables (latitude, longitude and sample age), number of human-classified reads, sample material, palaeoclimatic variables, human mobility and human ancestry constitute the linear predictors. The sample material is a categorical variable indicating whether the material used for sequencing was a tooth or not (bone), which *inlabru* treats as a random effect variable. We followed the default *inlabru* priors, where distributions are distributed as a Gaussian variable with mean µ and precision τ. The prior on the precision τ is a Gamma with parameters 1 and 0.00005. The mean is a linear combination of the covariates. By default, the prior on the intercept of the linear combination is a uniform distribution, while the priors on the coefficients are Gaussian with zero mean and precision 0.001. All covariates were normalised before the analyses. For each microbial species and group, we tested multiple models with different sets of covariates: 1) palaeoclimate + mobility + ancestry, 2) palaeoclimate + mobility, 3) palaeoclimate + ancestry, 4) only climate, 5) mobility + ancestry, 6) only mobility, 7) only ancestry, 8) no climate, nor mobility, nor ancestry. Spatiotemporal variables, number of human-classified reads, and sample material were included in all models. Because covariates were normalised, results indicate deviations from the mean. The effect size is interpreted in units of standard deviation. We used the deviance information criterion (DIC) to assess the model fit to each set of covariates, and prevent overfitting. The results shown in the main text are for the best-performing models (i.e., models with the lowest DIC score for each microbial species or combined species group). DIC scores as well as Watanabe–Akaike information criterion (WAIC) for each model, and results for all the other models we tested can be found in the Supplementary table S6.

## Data availability

All sequencing data used in this study is available as trimmed read files (FASTQ) at the European Nucleotide Archive under accession PRJEB65256. Processed analysis files including *KrakenUniq* database file and metagenomic profiling results, microbial species read alignments (BAM format) as well as per-sample summary tables and plots from screening pipeline are available at Zenodo under accession XX.

## Code availability

A *Snakemake* workflow implementing the computational screening pipeline is available at https://github.com/martinsikora/pathopipe.

## Acknowledgements

The Lundbeck Foundation GeoGenetics Centre is supported by the the Lundbeck Foundation (R302- 2018-2155, R155-2013-16338), the Novo Nordisk Foundation (NNF18SA0035006), the Wellcome Trust (UNS69906), Carlsberg Foundation (CF18-0024), the Danish National Research Foundation (DNRF94, DNRF174), the University of Copenhagen (KU2016 programme) and Ferring Pharmaceuticals A/S to E.W. Additional support was provided by Germany’s Excellence Strategy (EXC- 2077), project 390741603 “The Ocean Floor – Earth’s Uncharted Interface”. We thank A. Razeto, P. Selmer Olsen, for administrative and technical assistance. We thankfully acknowledge Illumina Inc. for collaboration. EW thanks St. John’s College, Cambridge, for providing a stimulating environment of discussion and learning. This work was further supported by the Swedish Foundation for Humanities and Social Sciences grant (Riksbankens Jubileumsfond M16-0455:1) to KK. M.E.A. was supported by Marie Skłodowska-Curie Actions of the EU (grant no. 300554), The Villum Foundation (grant no. 10120) and Independent Research Fund Denmark (grant no. 7027-00147B). G.S. is funded by Marie Skłodowska-Curie Individual Fellowship ‘PALAEO-ENEO’ (grant agreement number 751349). H.S. was supported by a Carlsberg Semper Ardens Grant (grant no. CF19-0601) and an ERC Consolidator Grant (grant no. 101045643). F. R. is supported by a Villum Young Investigator Grant (project no. 00025300), a Novo Nordisk Fonden Data Science Ascending Investigator Award (NNF22OC0076816) and by the European Research Council (ERC) under the European Union’s Horizon Europe programme (grant agreements No. 101077592 and 951385). N.O., R.Å., L.H. and B.N. are financially supported by Knut and Alice Wallenberg Foundation as part of the National Bioinformatics Infrastructure Sweden at SciLifeLab. A.K.N.I. and L. F. thank the OAK Foundation.

## Author contributions

M.S. and E.W. conceptualized the study. M.S., E. C., A. F. G., S. H. N., A.K.N.I, and F. V. S analysed data. M.S., E.C. A. F. G., N. O., R. A., L. H., E. K. I.-P., B. M., S. H. N. and H.S. were involved in method development and implementation. G. S., M. E. A., F. V. S., H.S., C. G. J. S. and L. V. were involved in data generation. M. S., M. E. A., K. G., and K. K. curated bioarchaeological data. M.S. T. C. J., B. N., J. P., L. F., F. R. and E.W. supervised the research. M.S., A. K. N. I. and E.W. wrote the first draft of the paper. M.S, A. K. N. I., L. F. J. P. and E.W. were involved in reviewing drafts and editing. All co-authors read, commented on and agreed on the submitted manuscript.

---

**Extended Data Fig. 1.**
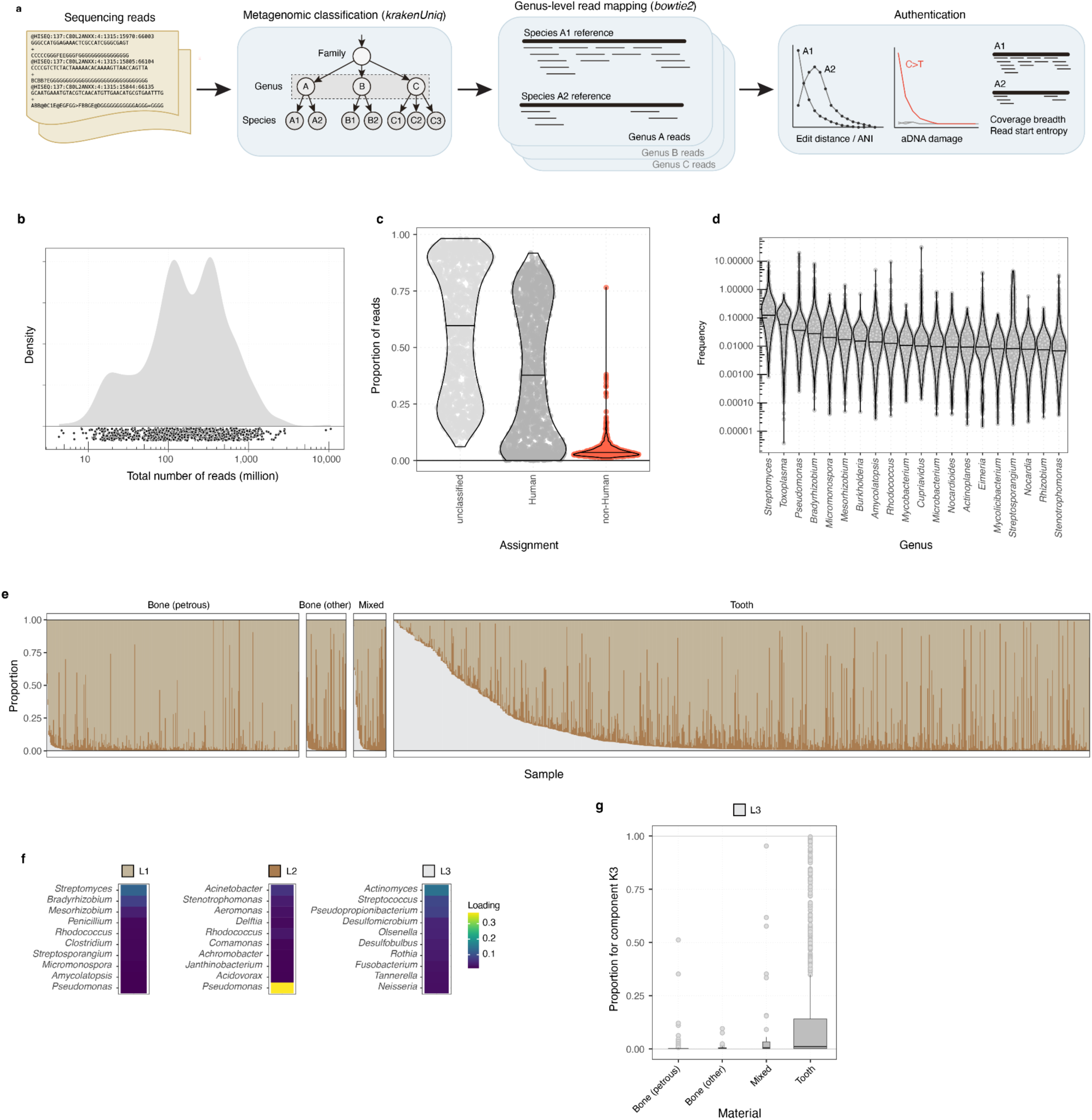
Workflow overview and metagenome composition. **b,** Distribution of total number of sequencing reads screened across the 1,313 study samples. **c,** Violin plots showing distributions of proportions of reads classified as human, non-human or not classified for the study samples. Median values for each genus are indicated by horizontal lines. **d,** Violin plots showing fraction of reads classified on the taxonomic level of genus, for the top 20 most abundant genera. **e,** Barplots showing inferred proportions for L=3 topics (indicated by fill colour) from topic model analysis for 1,272 study samples with sample material information. **f,** Factor loadings for the 10 highest loading genera for each of the L=3 topics from the topic model analysis. **g,** Boxplots showing distributions of proportions for topic K3 (associated with oral microbiome taxa) in different sample materials.

**Extended Data Fig. 2.**
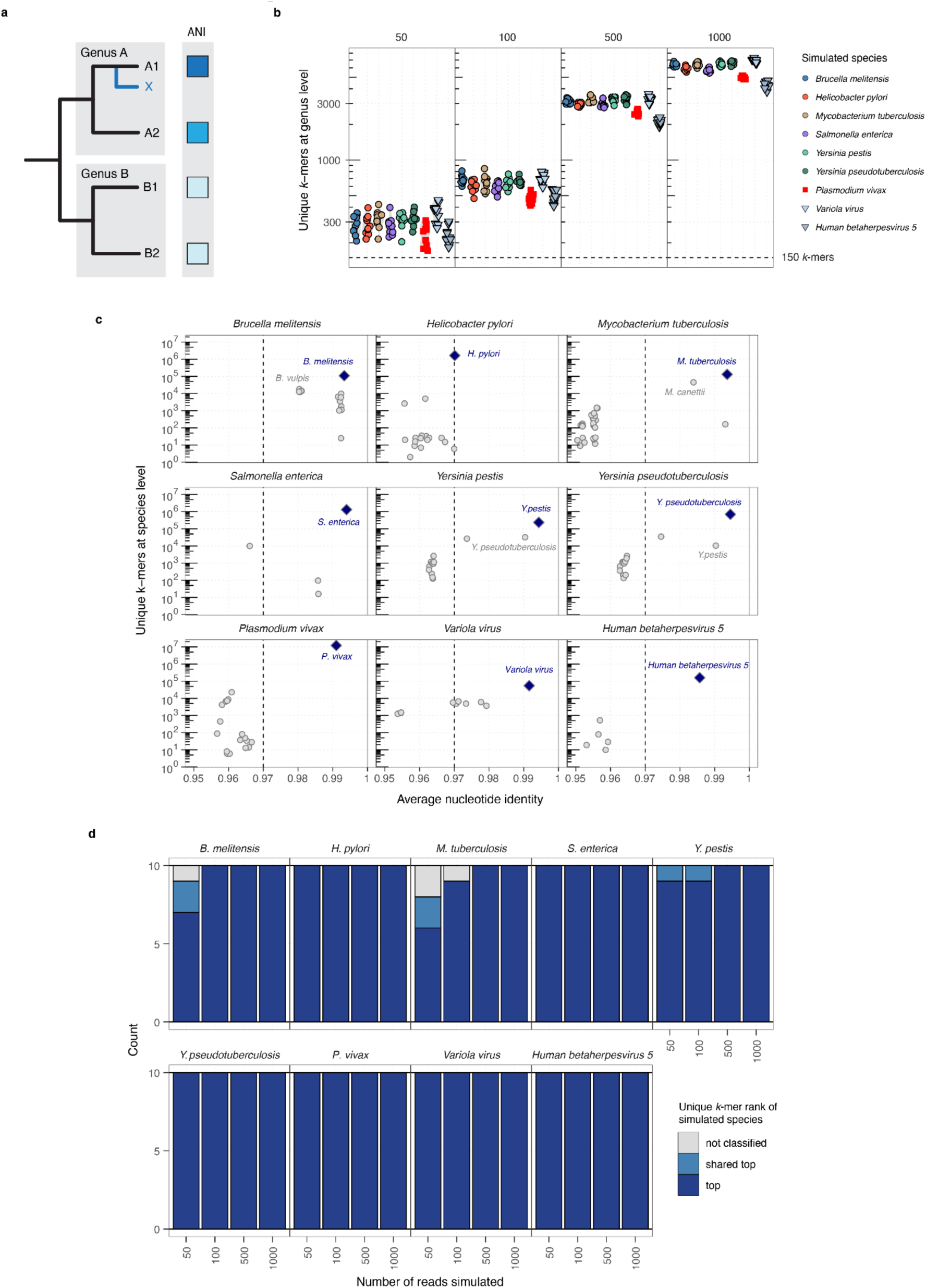
Reference genome similarity in simulated ancient microbial data. **a,** Illustration showing phylogenetic context and expected average nucleotide identity (ANI) for a hypothetical sampled microbial species X and four genomes (A1, A2; B1, B2) of two genera (A, B) present in the reference database. **b,** Number of unique *k*-mers classified at the level of genus using *KrakenUniq* for replicates of different read numbers across all simulated species. Dashed line indicates cutoff used in analysis of real data (150 unique *k*-mers). **c**, Number of unique *k*-mers classified at the level of species as a function of average nucleotide identity for mappings against all individual species reference genomes in the genus of reads simulated for a particular species. Blue diamonds indicate results for the mapping against a reference genome from the same species as the simulated read data, whereas grey circles indicate reference genomes of other species. Selected individual species results are highlighted by species name. Dashed line indicates ANI ≥ 0.97 cutoff value. **d**, Barplots showing number of replicates where the true positive species reference genome was highest ranking in numbers of unique *k*-mers classified at level of species.

**Extended Data Fig. 3.**
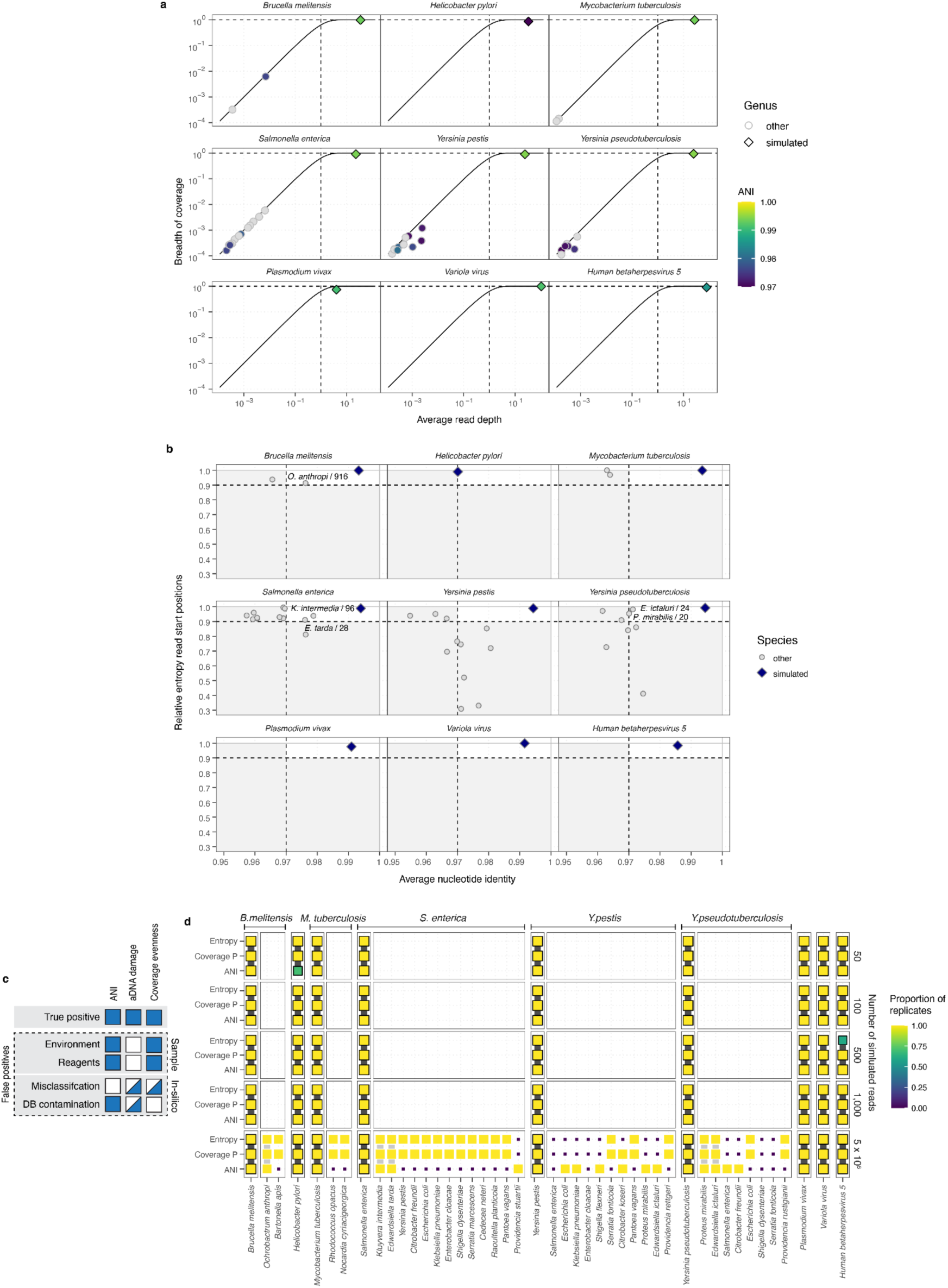
Read mappings across genera in simulated ancient microbial data. **a,** Observed breadth of genomic coverage as a function of average read depth for distinct species hits (i.e., mappings with highest number of unique *k*-mers at species level for a genus; n ≥ 20 reads mapped). Each panel shows results for reads simulated from species indicated. Results for mappings against the simulated species are indicated by diamond shape, whereas mappings against species from other genera are indicated with circles. Symbol fill colour indicates average nucleotide identity for mapped reads (grey symbols ANI < 0.97). Solid black line shows theoretical expected breadth of coverage for a given average read depth^80^. Vertical dashed line indicates 1X average read depth. **b,** Relative entropy statistic (1000 bp window size) as a function of average nucleotide identity. Blue diamonds indicate results for the mapping against reference genome from the same species as the simulated read data, whereas grey circles indicate reference genomes for species hits in other genera. Dashed lines indicate cutoffs used in analyses of real data (ANI ≥ 0.97, entropy ≥ 0.9). False positive hits of reads mapped to a reference genome from a different genome passing cutoffs and their final number of mapped reads (out of 5 million total simulated reads) are labelled. **c**, Illustration showing potential sources of false positive hits and expected results for authentication summary statistics. **d**, Matrix plot showing all microbial hits with n ≥ 20 reads mapped and their authentication statistics, for all simulated species and read numbers. Symbol colour and size indicates the number of replicates passing the cutoff for each of three summary statistics shown (ANI ≥ 0.97, ratio of observed / expected coverage breadth ≥ 0.8, entropy ≥ 0.9). Hits passing cutoffs for all three statistics are indicated with coloured outline and background lines (black - true positives; grey - cross-genus false positive mappings).

**Extended Data Fig. 4.**
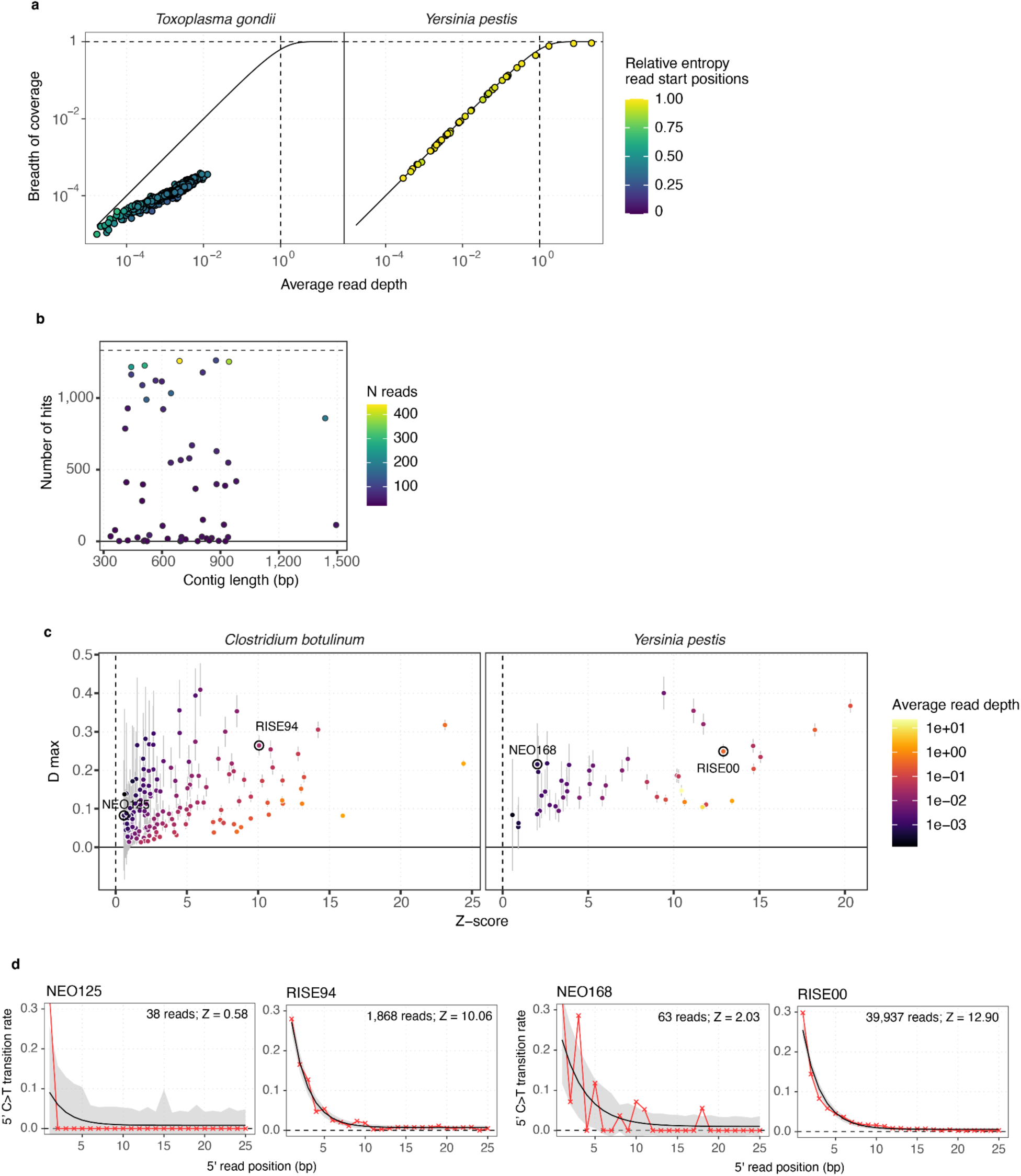
Examples of authentication for microbial hits. **a**, Observed breadth of genomic coverage as a function of average read depth. Coloured symbols indicate hits in species *Toxoplasma gondii* (left panel) and *Yersinia pestis* (right panel), with symbol colour indicating relative entropy of read start positions. Solid black line shows theoretical expected breadth of coverage for a given average read depth^80^. **b**, Lengths of contigs in the reference genome of *Toxoplasma gondii* and number of samples showing n ≥ 20 reads mapped. Symbol colour indicates the average number of reads mapped to a specific contig across samples. **c,** Bayesian estimator of aDNA damage (D max) and significance (Z-score) obtained from *metaDMG,* for hits in species *Clostridium botulinum* (left) and *Yersinia pestis* (right). Error bars indicate ± 1 standard deviation, and symbol fill colour indicates average read depth for mapped reads. Samples used as examples in aDNA damage curves (d) are labelled and indicated with black circles. **d**, aDNA damage patterns for four example hits in species *Clostridium botulinum* and *Yersinia pestis.* Plots show observed nucleotide misincorporation frequencies (red symbols and line) and *metaDMG* fit (black line) and 68% credible intervals (shaded region) for C>T transitions as a function of distance from the 5’ read end.

**Extended Data Fig. 5.**
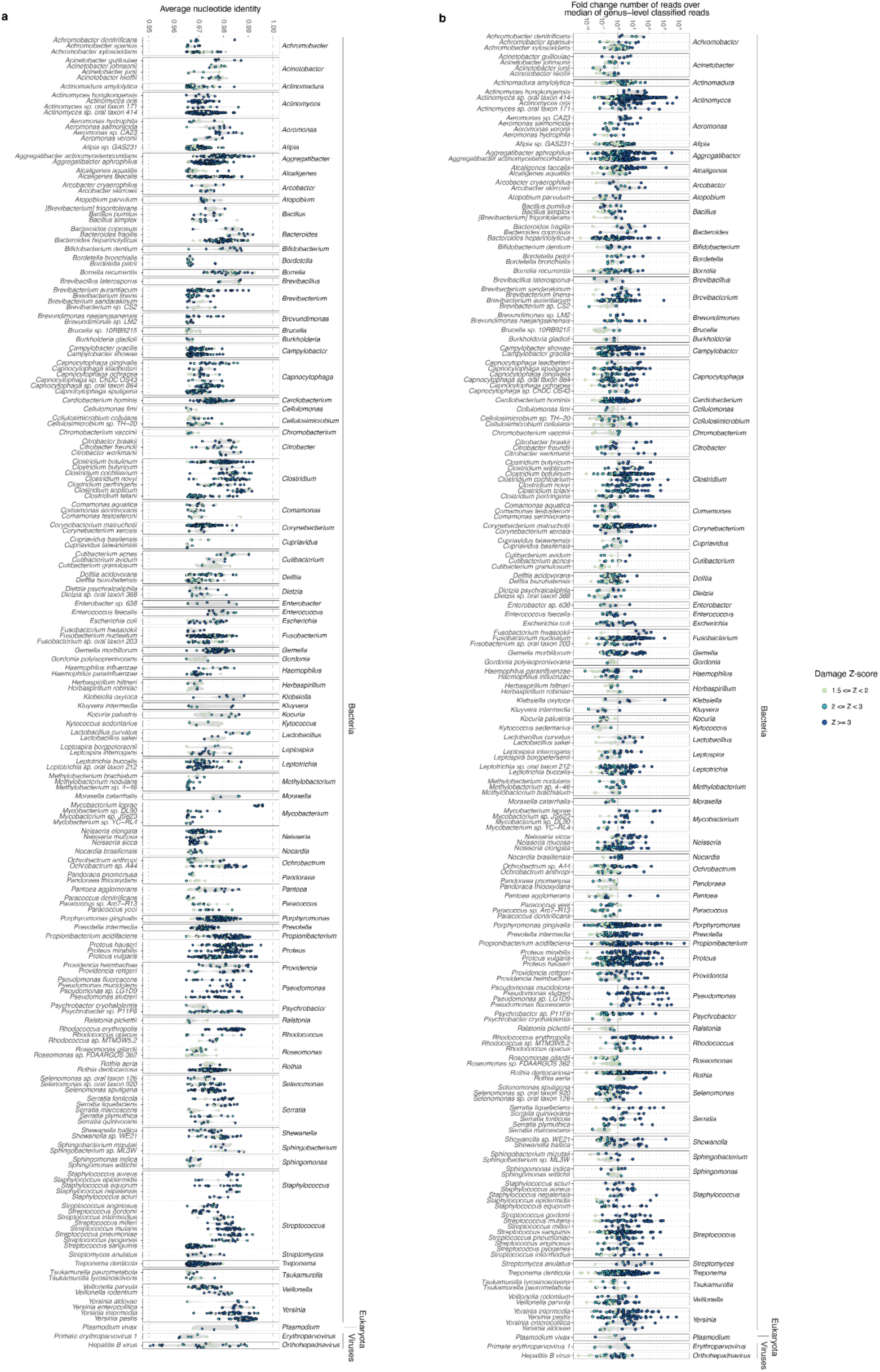
Ancient microbial hit ANI and read recruitment. **a, b,** Distributions of ANI (**a**) and log10-fold change of mapped reads over median of reads classified at taxonomic rank of genus per sample (**b**) for individual species hits detected in n ≥ 5 samples. Symbol colour indicates species hit category.

**Extended Data Fig. 6.**
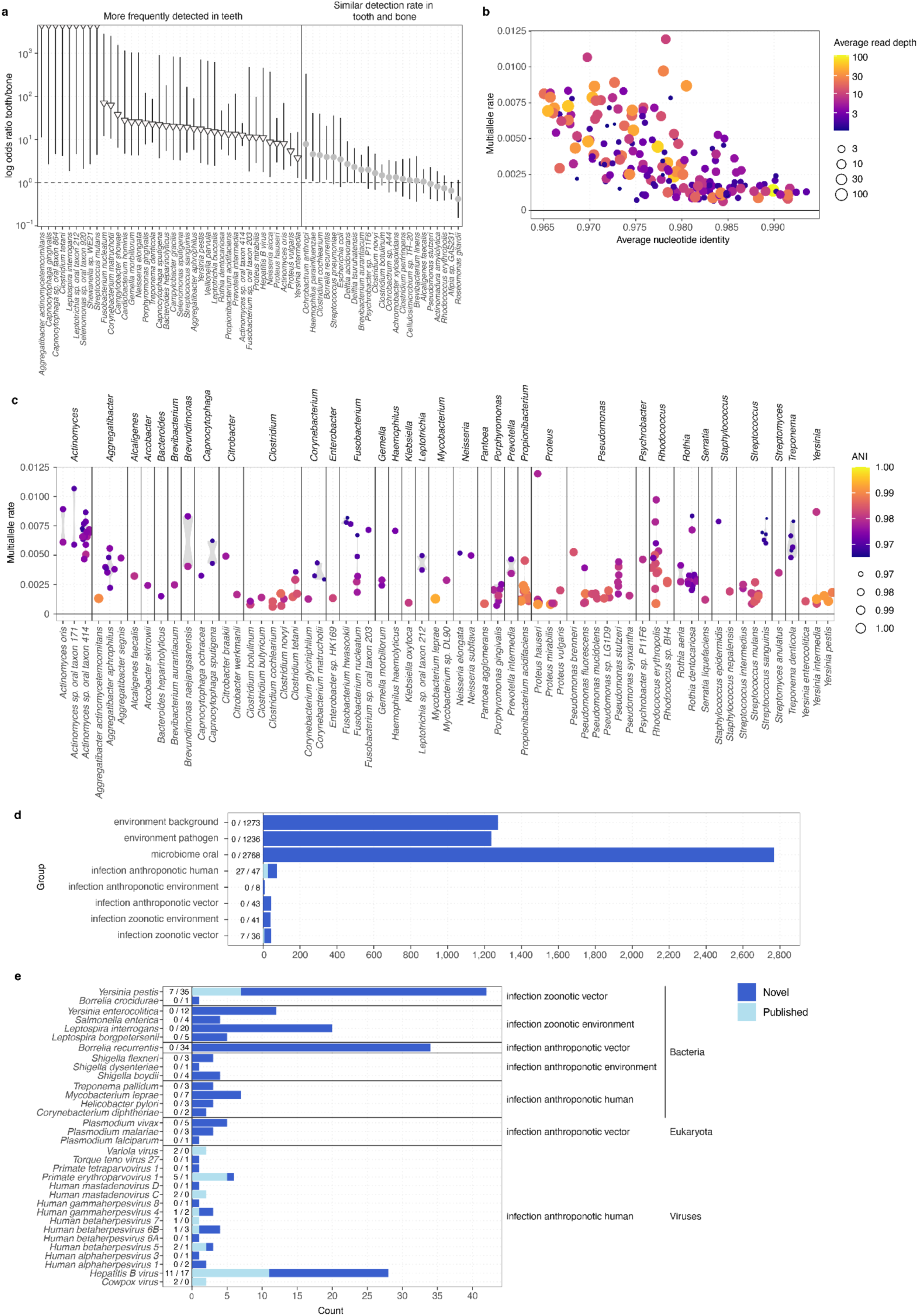
Ancient microbial hit characteristics. **a,** Odds ratios for association of ancient hits with sample material (tooth or bone) across 61 species with ≥ 20 ancient hits. Symbols indicate significance of association (p ≤ 0.01, Fisher’s exact test; white triangles - more frequently identified in tooth; grey circles - no significant association). Error bars indicate 95% confidence interval of odds ratio **b**, **c,** Rates of observing multiple alleles in 2 randomly sampled sequencing reads at genomic sites in 190 ancient hits (average read depth ≥ 1X) across 120 samples**. b,** Multi-allele rate as a function of ANI. Symbol colour indicates average read depth. **c,** Distribution of multi-allele rate across species hits. Symbol colour indicates ANI. **d, e** Barplots showing number of hits identified in each microbial species group (**d**) or each species within groups of likely infections (**e**). Novel and previously reported ancient pathogen hits are distinguished by bar colour, with total number in each category labelled.

**Extended Data Fig. 7.**
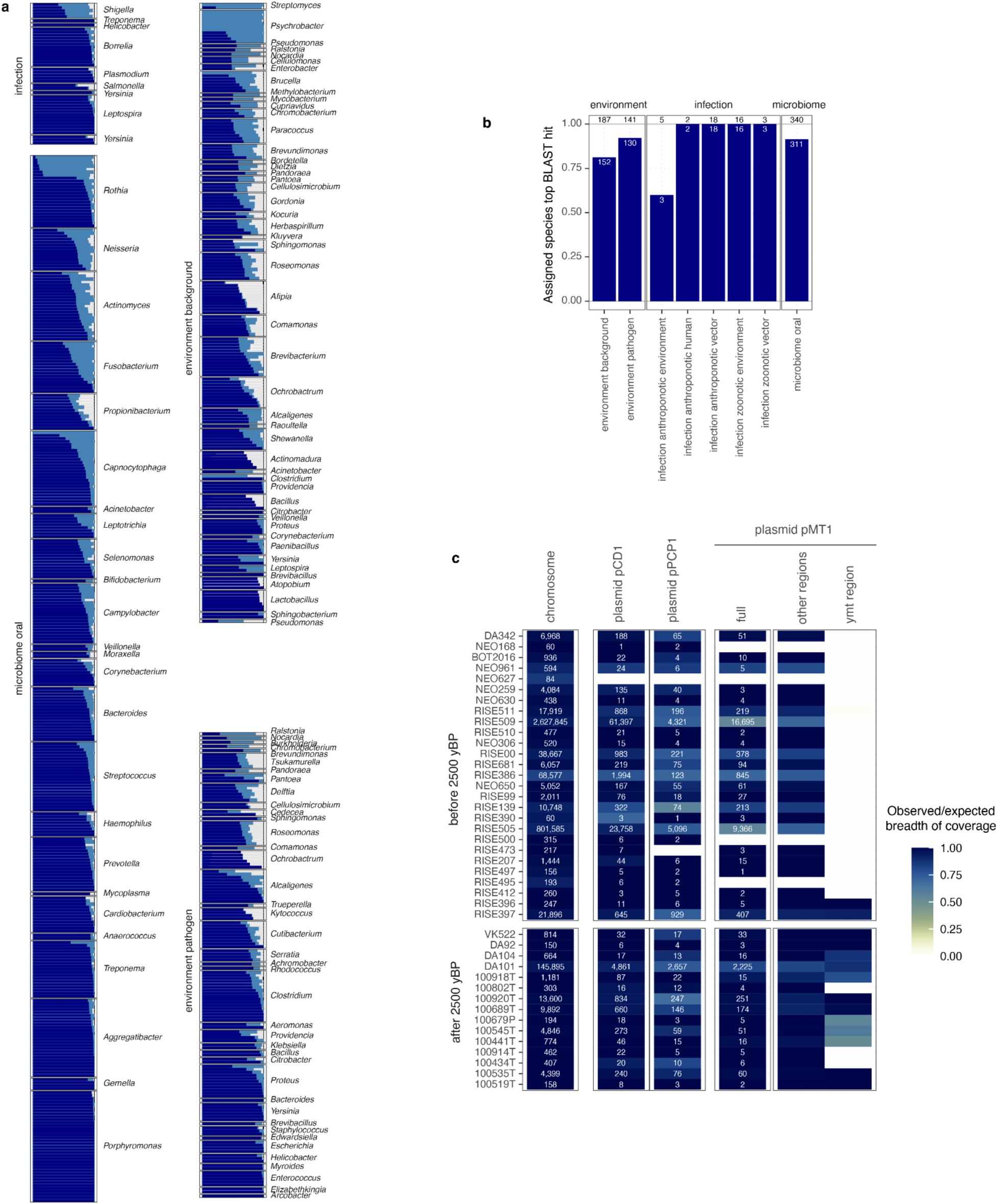
Additional ancient microbial hit authentication. **a,** Bar plots showing proportion of reads assigned to same species (dark blue) or genus (light blue) using BLASTn for all hits with N ≤ 100 final reads (N=712), stratified by genus and microbial source groups **b,** Bar plots showing the proportion of ancient microbial hits with N ≤ 100 final reads matching the species with most reads assigned using BLASTn, stratified by microbial source group. **c,** Heatmap showing number of reads mapped to *Yersinia pestis* CO92 chromosome and plasmids, for N=42 *Yersinia pestis* hits. Cell color indicates ratio of observed over expected breadth of coverage. Results for plasmid pMT1 are shown for full plasmid, as well as separately for the 19 kb region contaiinng the *ymt* gene absent in the LNBA- strains. Samples are ordered by decreasing age from top to bottom.

**Extended Data Fig. 8.**
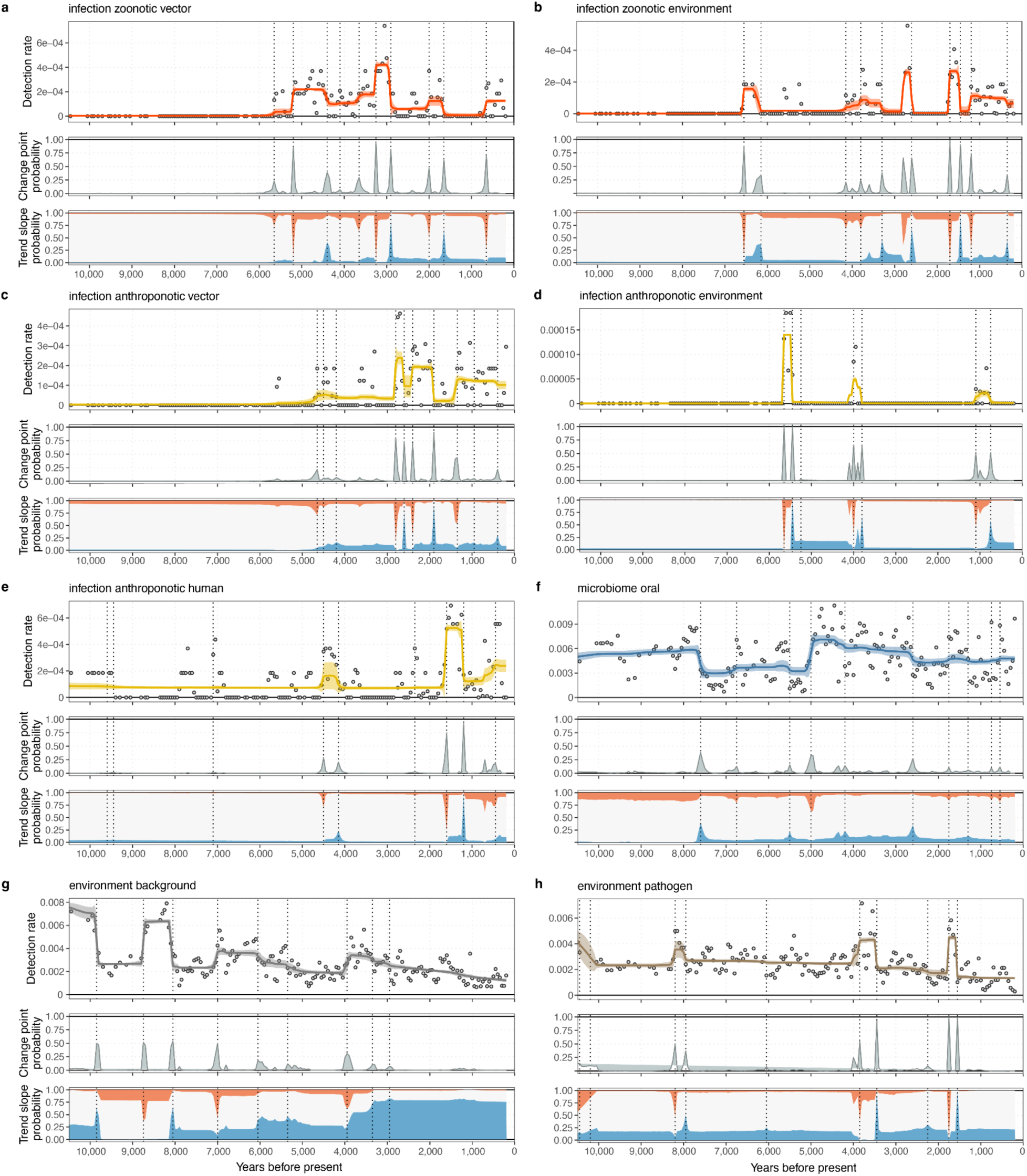
Time series of detection rates for ancient microbial groups. **a-h,** Panels show estimated trendlines and 95% credible interval for detection rates (top), probability distributions and locations (dotted lines) for change points (middle) and probability of trend slope (bottom) being positive (red), negative (blue) or zero (white), inferred using Bayesian change-point detection and time series decomposition.

**Extended Data Fig. 9.**
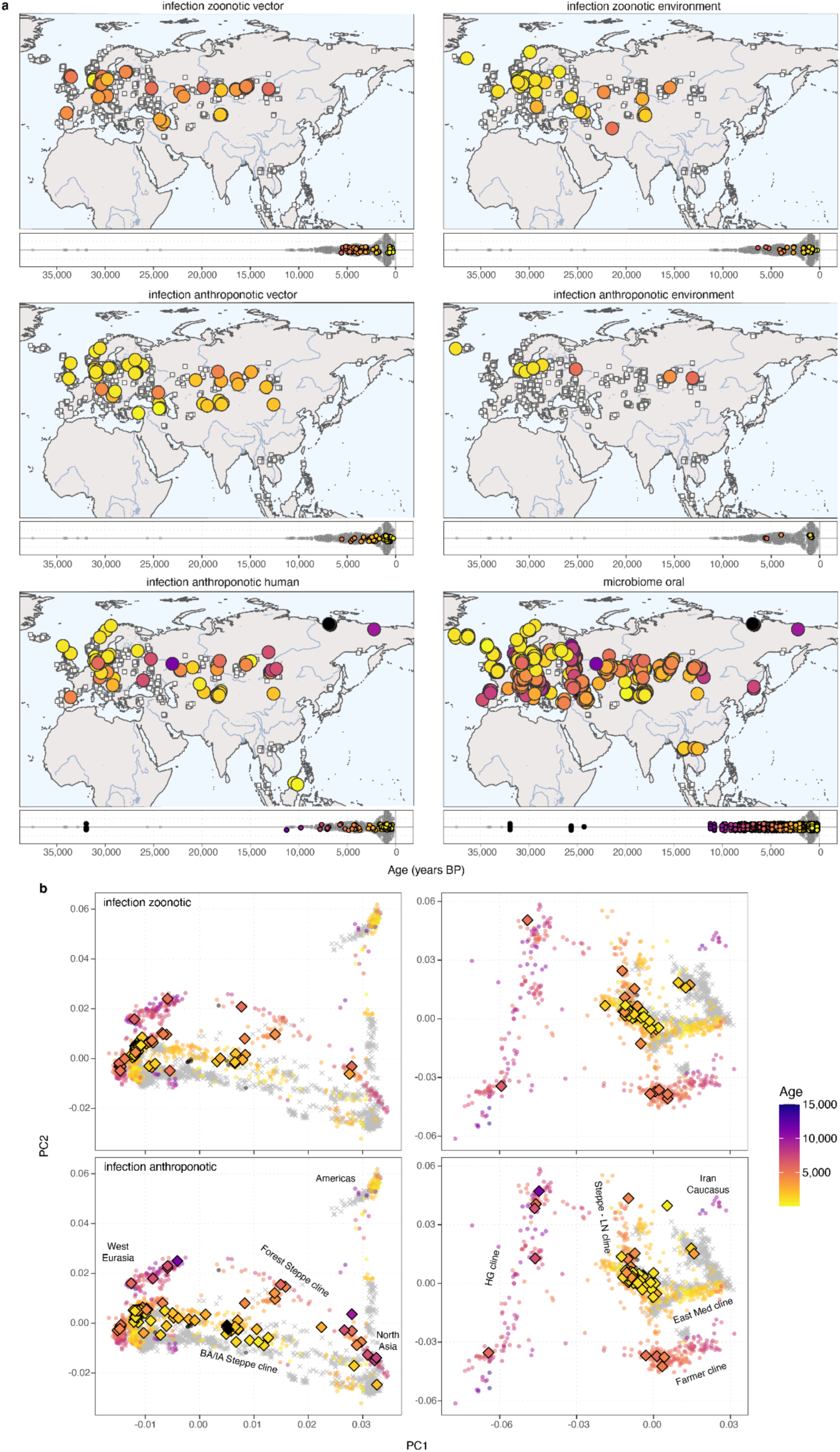
Spatiotemporal distribution and host genetic structure for ancient microbial groups. **a,** Panels showing geographic distributions (top) and timelines (bottom) for identified cases of ancient microbial hits in the oral microbiome and infection groups classes (indicated by coloured circle). Geographic locations and age distributions of all 1,313 study samples are shown in each panel using white squares. **b,** Principal component analyses showing ancient and modern human genetic population structure in non-African (left panels) and west Eurasian (right panels) individuals. Grey crosses indicate present-day individuals, whereas coloured symbols indicate ancient individuals (coloured by sample age). Diamonds with black outlines indicate position in PCA space for samples with hits in combined infection groups. Major clines of known ancient and modern human ancestry groups are indicated with labels.

**Extended Data Fig. 10.**
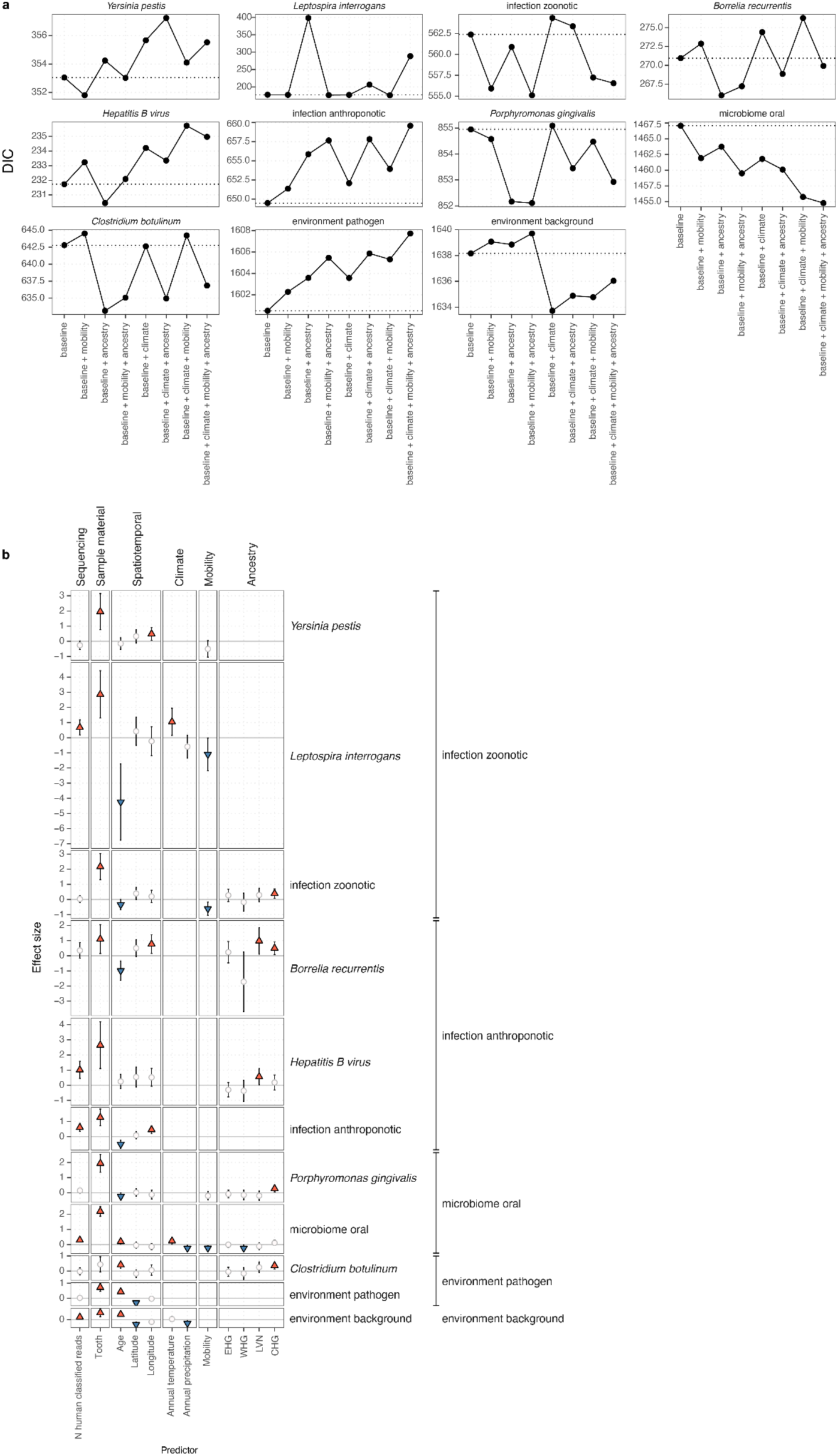
Predictors of ancient microbial species incidence. **a,** Watanabe–Akaike information criterion values for each model and response variable. **b,** Matrix showing effect sizes and of 12 potential predictors (columns) for presence of selected combined ancient microbial species and combined groups inferred from spatiotemporal modelling. For each class, the model with lowest Watanabe–Akaike information criterion is shown. Symbols indicate the predictors included in the respective model. Predictors with positive effect (2.5% and 97.5% posterior quantiles both positive) are shown as red triangles, whereas predictors with negative effect (2.5% and 97.5% posterior quantiles both negative) are shown as blue inverted triangles. Predictors included in the best-fitting model but without effect (posterior quantile range spanning zero) are indicated using white circles. Posterior standard error of effect sizes is indicated by error bars.

