## Supplementary information for "The spatiotemporal distribution of human pathogens in ancient Eurasia and the emergence of zoonotic diseases"

Supplementary information 1: Expanded workflow description 1
Supplementary information 2: Additional site descriptions 3
Supplementary information 3: The necrobiome 9
Supplementary information 4: Authentication details for selected ancient microbial species in the “infection” group. 11
Supplementary information 5: Borrelia recurrentis and louse-borne relapsing fever 36
References 40

#### Supplementary information 1: Expanded workflow description

To understand the distribution of ancient pathogenic challenges, we developed a workflow to identify ancient microbial DNA in shotgun-sequenced aDNA data consisting of three main components. First, in the metagenomic classification stage, sequencing reads are subjected to an efficient *k*-mer-based metagenomic classification using *KrakenUniq*[1] against a comprehensive database of complete bacterial, archaeal, viral, and protozoan genomes from the RefSeq database[2] and the human reference genome. This allows rapid grouping of the complete set of sequencing reads according to taxonomic rank, with no requirement for computationally more costly sequence alignment. We use the taxonomic rank of genus as the unit of grouping. Second, in the read mapping stage, the set of reads for a particular genus are aligned with *bowtie2*[3] against each best-matching reference assembly for all species within the respective genus, selected based on the maximum number of unique *k*-mers assigned to the assembly by *KrakenUniq*. Finally, in the authentication stage, a broad set of summary statistics are calculated to facilitate authentication of sequences based on the following key characteristics expected to be observed for true positives (Extended Data Fig. 1a)[4,5]:

1) Highest similarity to the reference genome representing the true species, quantified using statistics such as read edit distance and average nucleotide identity (ANI)
2) Characteristic degradation patterns of aDNA, in particular aDNA damage through deamination of cytosine residues, manifesting as increased C>T (and complementary G>A) substitution rates at the read ends.
3) Even distribution of the mapped sequences across the reference genome[6], quantified through statistics measuring evenness of genomic coverage such as expected breadth of coverage or read start position entropy (Methods).

To determine parameter values for the workflow facilitating the recovery of true positive ancient species, we applied it to simulated sequencing reads with characteristics of ancient DNA from assemblies not present in our database for a set of nine known pathogen species (bacteria n=6; eukaryota n=1; viruses n=2) . We found that the true positive species within a genus was reliably identified from as low as 50 simulated reads when selecting the species with the highest number of unique *k*-mers classified after an initial filter of $n \geq 150$ unique *k*-mers classified at the genus level (Extended Data Fig. 2). False positives from initial misclassification of reads to a different genus on the other hand were characterised by either low ANI and/or uneven coverage after read mapping (Extended Data Fig. 3). We observed a few cases of false-positive species hits (Extended Data Fig. 3b,d), but these were generally of lower ANI (ANI < 0.98), and only occurred in the simulations with highly abundant true positive species ($5 \times 10^6$ sequencing reads). As analysis of real data additionally includes ancient DNA damage patterns for authentication, such false positive results are thus only expected when a closely related ancient microbial species is present in the sample at high abundance. We nevertheless suggest caution in interpreting individual ancient species hits with low ANI and read numbers, particularly in scenarios of species where all hits show these characteristics.

Based on the results of the simulations, we selected the species with the highest number of unique *k*-mers assigned by *KrakenUniq* among the set of species with n ≥ 20 reads mapped and passing all authentication criteria (Methods) as the putative ancient microbial species "hit", for each sample and target genus. A particular sample can thus have multiple species hits across different genera, but for each individual genus at most a single species hit is reported. A limitation of this approach is therefore that other potential true positive ancient microbial hits deriving from a different species within the same genus will be missed. However, all species irrespective of their pathogen status are included in the read mapping and can result in a hit for a genus, thus avoiding potential bias that could occur if only specific pathogen species were targeted.

### Supplementary information 2: Additional site descriptions

#### Nerkin Naver, Armenia[10]

Kurgan cemetery, about 30 km to the west of Yerevan, excavated from 2002 onwards. A series of tombs (N1 to N7) contained a large variety of archaeological artefacts as well as animal offerings, evidencing the high status of the persons that were buried in the tombs. Most of the tombs contained single burials, with the exception of tomb 5, which contained three burials.

RISE399, tomb 4. Tooth from a mature male, 50-55 years old. Directly dated to the Bronze Age.

#### Cachovice, Czech Republic[11]

Large cemetery, excavated due to coal mining 1980-1982, with 53 Corded Ware and 21 Bell Beaker graves. The site was only briefly published (Schwarz 2008).

RISE571, grave 39. Tooth.

RISE572, grave 44. Tooth.

#### Bodal mose (Bodal tørveskær), Zealand, Denmark

FoF no 030309-258

Museum no: NM1 333/45; AS 46/45, 33:14

Beheaded cranium and three neck vertebrae of an adult male found during peat digging in 1945. No traces of the rest of the skeleton. Older healed trauma on the cranium. Found with Iron Age beads and bone comb. 14C-dated to the Iron Age.

Sample: NEO881, CGG_2_021417. Left lower pre-molar of adult, probably male.

#### Bodal mose?, Zealand, Denmark

Museum no: NNU 4800, AS 13/99

Fof no 030309-180A

Bog find from 1940/1945 of uncertain provenience, possibly Bodal mose. 14C-dated to the Iron Age.

Sample: NEO562, CGG_2_018466. Left upper canine tooth.

#### Egedal mose, Zealand, Denmark

Also called Ølstykke. Bog find from 1944 during peat digging. Skeleton of adult male, 35-45 years old. A bronze sword was found nearby. Directly dated to the Early Iron Age.

Fof no 010607-64

Museum no: NM B 14367

Sample: RISE274. Upper right canine tooth.

#### Nedergård, Jutland, Denmark

VHM 065/1943, 1945/0020, 100109-13

Single mandible found in a bog during peat digging in 1943. 14C-dated to the Late Bronze Age/Early Iron Age.

Sample: NEO953, CGG_2_023281. Tooth and mandible.

##### Lauda-Königshofen, Baden-Württemberg, Germany[12,13]

Lauda-Königshofen "Wöllerspfad" is located in the valley of the Tauber in northeastern Baden-Württemberg. The Lauda-Königshofen site was excavated between 1998 and 2000 by C. Oeftiger. The cemetery covered an area of at least 150 x 120m, but was not completely excavated. In addition to the Corded Ware burials, some metal age settlement remains were also found. The geology of the Tauber valley is characterised by loess deposits, and the Lauda-Königshofen cemetery is located on loess soil, on the lower terrace of the river.

A series of CW cemeteries have been excavated in the Tauber valley. There are three large cemeteries known and some 30 smaller sites. The largest is Lauda-Königshofen with 91 individuals. The cemeteries are dispersed rather regularly along the Tauber valley, on both sides of the river, suggesting a quite densely settled landscape.

The Lauda-Königshofen graves consisted mostly of single inhumations in contracted position, usually oriented E-W or NE-SW (Fig 3). A total of 91 individuals were buried in 69 graves. At least 9 double graves and three graves with 3–4 individuals were present. In contrast to the common CW pattern, sexes were not distinguished by body position, only by grave goods. This trait is common in the Tauber valley and suggests a local burial tradition in this area. Stone axes were restricted to males, pottery to females, while other artefacts were common to both sexes. About a third of the graves were surrounded by ring ditches, suggesting palisade enclosures and possibly over-plowed barrows. The 14C dates place the cemetery in the middle part of the German CW.

Sample: RISE336

##### Regensburg-Dechbetten, Bavaria, Germany

Skull, long bones and bronze sword and dagger found during clay extraction.

Sample: RISE473, M1 tooth from adult male

##### Schöngeising, Bavaria, Germany[14]

Small group of barrows, excavated in 1892 by J Naue. Only one of them, barrow 1, was well documented. An Iron Age cremation was found in the upper part, and deeper down a stone frame with the skeleton of an adult male stretched out on his back. Trauma was seen on the cranium. A bronze needle (Lochhalsnadel) was found on the chest, and a pottery vessel ca 2 m from the skeleton.

RISE478, tooth 28, adult male

##### Wolkshausen, Bavaria, Germany[13,15]

This site is located near Würzburg in NW Bavaria. An Iron Age hillfort was excavated here 1983–1985. During the course of the investigations, nine CW burials were found. One individual from this site was sampled, grave 9/RISE440. This grave contained an adult male skeleton in contracted position on his right side, head towards the west. The grave goods consisted of a stone axe, a flint flake, a flint blade, a hammer stone, three bone implements, a few pottery shards, and a few animal bones.

Sample: RISE440, grave 9

#### Jegvermi-Kert (Bekes 103), Hungary[16]

Large MBA cemetery by the Körös river, excavated by Paul Duffy 2011-2015. The minimum number of individuals was estimated to 68, of which only 5 were inhumations and the rest were cremations. The site has only been briefly published.

Sample: RISE320, grave 3. Human bone, directly dated to the Middle Bronze Age.

#### Veszto-Magor, Hungary[17]

Tell site, excavated on several occasions from 1968 to 1986. It covers 3.9 ha and is up to 9 m high. After a first occupation in the Middle and Late Neolithic, it was reoccupied during the Chalcolithic as well as the Middle Bronze Age, and a monastery was built on top of it in the Medieval period. In addition to settlement material, some 33 Neolithic and 19 Early Copper Age burials were also found. The date information suggests two periods of burial, with 7 graves dated to the Late Neolithic and 3 graves to the early Chalcolithic. Two of these are included here, both dated to the Late Neolithic.

Samples:

RISE377, grave 4. Tooth 15.

RISE379, grave 16. Canine tooth of a mature male.

#### Arano, Lombardy, Italy[18]

Arano is a large Early Bronze Age cemetery in the commune of Verona in Lombardy. It was excavated in 2007 by Salzani. 68 burials were documented, containing a minimum number of 73 individuals. All but one were flat inhumation graves. They were buried in a contracted/hypercontracted posture, with flexed legs and arms, lying either on their left side with the head to the south, or on their right side with the head to the north. The data would seem to indicate that these two distinct types of positioning might be sex-dependent (left-crouched males and right-crouched females, with the head in opposite directions). Grave goods were sparse and only found in 13 graves. A series of 17 14C datings confirms a rather short period of use, probably less than 200 years, and a general dating to the Early Bronze Age.

Samples:

RISE533, grave 40, right M2 tooth.

RISE540, grave 32, left M3 tooth of an adult female.

#### Olmo di Nogara, Veneto, Italy[19]

This cemetery, located in the Po river basin, was excavated by Salzani in 1998-2009. It is one of the largest Bronze Age cemeteries in Italy dating to the Middle and Late Bronze Age. The graves are predominantly inhumations, and ca 10% cremations. The deceased are usually placed in supine position with arms straight at the sides or bent over the basin. There is also evidence of on the side or prone placements. The burials are often accompanied by sets of artefacts: generally weapons for males and an ornamental parure, bronze pendants and combs made of bone/horn for females.

Sample: RISE470, grave 33. Right lower M1 tooth from an adult male, 30-40 years old. He was buried in an extended position on his back, with a bronze sword and a bronze dagger.

#### Valdaro, Lombardy, Italy[20]

Excavations in 2007 in San Giorgio in the outskirts of Mantua revealed 20 single and one double inhumation burial, dating from the Chalcolithic to the Bronze Age. The skeletons were placed in contracted positions on the side. The double burial (T2) contained a male and a female (later reclassified as two males), and became famous as the “lovers of Valdaro”. Five graves have been dated within the RISE project, giving consistent dates to the Chalcolithic, ca 3300-3100 cal BC. This should also apply to the individual included here, RISE521.

Sample: RISE521, grave 8. Lower right M2, juvenile-young adult. Dating failed.

#### Oblaczkowo 7, Great Poland, Poland[21]

The Corded Ware grave in Obłaczkowo was unearthed in the course of rescue excavations carried out between 2006 and 2008. The grave was located on a top of a small hill and no traces of a burial mound were found. It was rectangular in shape (2,5x2,15 m) and oriented N-S. It contained skeletal remains of two adult individuals and a child, all buried in contracted position on a side. The grave offerings included a battle-axe and decorated antler plates (56). DNA results from the child and an adult were published by Malmström et al (2019).

Sample: RISE1. Dm1 tooth from child ca 6 years. Directly dated to the early Corded Ware period.

#### Torre Velha 3, Baixo Alentejo Portugal[22]

Large cemetery with chalcolithic pit graves, 25 BA hypogeums, IA and Roman flat graves. Excavated 2008-2009 by the company Palimpsesto Lda. Various mortuary treatments were recorded in the hypogeums, including 23 skeletons, one ossuary, five bone heaps, two cases of isolated bones and 48 funerary deposits, consisting of pottery, metal and bone artefacts, and animal bones.

Sample: RISE543, tooth from skeleton 1382 in hypogeum 1489-1490. Male, adult, >30 years. Direct dating failed, but a cattle bone associated with skeleton 1382 was previously dated to the Bronze Age, 3300+-45 uncal BP, SAC-2489.

#### Arban 2, Khakkassia, Russia

Cemetery, Late Bronze Age Karasuk culture. Excavated by Savinov 1986-1987.

Sample: RISE498, 7332-189. Tooth of adult male, 35-50 years old. Not directly dated.

#### Kam-Tyttugem, Altai, Russia

Mummy found by schoolboys in a wooden coffin under a rock shelter. Context dated to the Iron Age, 300-500 AD.

Sample: RISE603, tooth.

#### Lebedy-I, Krasnodarsk, Russia

Cemetery, context dated to the Catacomb culture, ca 3000-2000 BC.

Sample: RISE592, Lb-I-5-14. Lower M3 tooth from a male.

#### Oleniy-I, Krasnodarsk, Russia

Cemetery of the Bronze age Srubnaya culture.

Sample: RISE589, Ol-I-2-23. Lower PM1 from a male. Directly dated to the Neolithic.

##### Orak Ulus, Khakkassia, Russia[23]

Cemetery consists of oval kurgans with rectangular stone enclosures. Classed as belonging to the Bronze Age Federovo variant of the Andronovo culture. Excavated by Sosnovski in 1926-1928. Details of burials are not well recorded. 11 DNA samples were analysed by Narasimhan et al (2019).

Sample: RISE501, 3390-2. Tooth of female. Not directly dated.

##### Plastynovsy-I, Krasnodarsk, Russia

Cemetery with graves spanning ca 4000-2000 BC.

Samples:

RISE587, Pl-I-1-15. Upper M2 tooth from a male?. Context dated to the Catacomb culture, 3000-2000 BC

RISE588, Pl-I-1-20. Lower M2 tooth from a male. Context dated to the North Caucasian, 3000-2000 BC

RISE590, Pl-I-1-34. Lower M1 tooth from a male. Context dated to the Maykop, 4000-2900 BC

RISE591, Pl-I-1-30. Lower M1 tooth from a female?.  Context dated to the Novosvobodnensk, 4000-3500 BC

##### Sukhaya Termista-I, Rostov, Russia

Eneolithic-Yamnaya kurgan, excavated by Shishlina in 2009. The grave included here was a shallow oval-shaped pit oriented in east-west direction, with a slight deviation to the north. Its dimensions are 1.9 by 1.07 m. Within the pit lay the skeleton of a female individual aged 18-25 years, positioned contracted on her right side, with the skull oriented west. The cranial bones were coated with ochre. A small bronze ring and a sheep bone were found in the grave.

Sample: RISE239, kurgan I grave 14. Lower left M1 tooth from adult female, 18-25 years old. Directly dated to the Eneolithic.

##### Zhana-Aul, Altai, Russia

Kurgan, partly damaged by construction work. Excavated by Kocheev in 1991. Context dated to AD 600-700.

Museum id: 1-1

Sample: RISE599, tooth from adult female.

##### Snorthög, Scania, Sweden[24,25]

This barrow and flat grave cemetery was excavated by Vifot in 1938. It consisted of a barrow, ca 2 m high and 21 m in diameter, with 11 BA cremations. Below the mound, a cemetery with 10 flat inhumation graves was found. Grave 7 was directly dated to the Early Neolithic, while the other inhumations were dated to the Late Neolithic. Nine of the graves were sampled for the RISE project, of which one (grave 5) is included here. This grave was an oval pit without stone construction, containing a male skeleton in crouched position on his left side, head to the SE. A leaf-shaped, bifacially flaked flint arrowhead was found in the chest area. Grave 5 has been dated twice, giving consistent dates to the Late Neolithic I.

Sample: RISE189, grave 5. Lower left PM1 tooth from an adult male?, 16-22 years.

Smr no: Lilla Isie 23:1

Museum no: LUHM 28424

##### Åkeshög, Scania, Sweden[26]

Bronze Age barrow, investigated by Montelius in 1896. The barrow contained a wooden coffin with a bronze dagger. Under the barrow was a small cairn with four skeletons, resin, quern stones, a flint scraper and a flint chisel. Samples come from this cairn, but details of their positions were not documented. Direct dating indicates that the skeletons belong to the MN A and MN B.

Sample: RISE270. Lower left M3 tooth of an adult, 25-35 years old. Directly dated to the early MN B

Museum no: SHM 10288

#### Supplementary information 3: The necrobiome

Following death, decomposition starts to take place. This process is studied in anthropological research centres using human donors and animals, e.g., pigs. Terrestrial mammalian decomposition is a dynamic process, and its progression is partly governed by environmental conditions such as temperature, rainfall, and soil composition. However, a universal microbial decomposer network appears to exist despite location, climate and seasonal effects[27]. Decomposition involves microbes, insects, and vertebrate scavengers, but in the following sections, we will only consider bacteria. The microbial community associated with decomposition is far from a static group of microbes. It undergoes dynamic changes over time, with bacteria and microbial eukaryotes engaging in a fierce competition for nutrients and even consuming each other[27,28]. This microbial succession is often likened to a clock, albeit one that requires calibration based on geography and season.

Initially, the bacterial flora is dominated by fast-growing, metabolically diverse, opportunistic bacteria such as *Pseudomonas* (our samples revealed several environmental *Pseudomonas* species, including *P. aeruginosa*, which also has been detected in saliva and dental plagues in patients[29]) and bacteria in the genus *Acinetobacter*. Bacteria such as *Wohlfahrtiimonas* (not detected in our samples) and *Ignatzschineria* (not detected in our samples) are carried to the decomposing body by blow flies soon after death. During advanced decay, bacteria in the genus *Acinetobacter* are again prominent, as these bacteria may be able to utilize diverse substrates. We observed several *Acinetobacter* species, e.g., *Acinetobacter baumannii* (oral microbiome, potential pathogen that can cause severe infections in many organs), *A. guillouiae* (environment, potential pathogen), *A. johnsonii* (oral microbiome), *A. junii* (oral microbiome, potential pathogen), *A. Iwoffii* (oropharynx (not strictly oral), skin, and perineum), *A. oleivorans* (environment), and *A. radioresistens* (skin), potential rare pathogen). Once a corpse has reached the dry remains stage, bacteria such as *Sporosarcina* (not found in our study) are detected.

Many of these decomposition-associated bacteria are ubiquitous[27,28]. They can be detected in low amounts both outside the human body and within it, for example, in soil and insects and the human gut, mouth and skin. *Acinetobacter* is commonly found in human skin and in low abundance in soil. Many other key bacteria associated with decomposition have been found to represent a unique phylogenetic diversity that was extremely rare or undetected in host-associated or soil microbial communities and rarely detected among our samples (*O. alkaliphile* (not found in our study), *Ignatzschineria* (not found in our study), *Wohlfahrtiimonas* (not found in our study), *Bacteroides* (previously believed to derive mostly from the human gut, several species detected), *Vagococcus lutrae* (not found in our study), *Savagea* (not found in our study), *Acinetobacter rudis* (not found in our study) and *Peptoniphilaceae* (found in one sample). Among the *Bacteroides* observed in this study, *B. heparinolyticus* predominated especially in dental samples (94/218 *Bacteroides hits*, 43%); *B. heparinolyticus* can be found in subgingival dental plaques[30]. The other *Bacteroides* species were environmental, some with pathogenic potential.

As a result, while some of the bacterial species identified in this study might be the result of environmental contamination or potentially part of the necrobiome, many were likely also present prior to death as part of the normal microbial flora or as an infection. This observation, coupled with the absence of most bacteria uniquely associated with the necrobiome, could suggest the necrobiome itself plays a minor role in our findings, which could be attributed to the age of the sampled remains. These findings underscore the need for further research to fully comprehend the microbial dynamics of decomposition over time and what traces of this process have been left after thousands of years.

#### Supplementary information 4: Authentication details for selected ancient microbial species in the “infection” group.

In this section we provide some additional authentication details for ancient microbial species hits in the “infection” group. Detailed summary statistics for each species hit are provided in Supplementary Table S5. Individual plots of read edit distance distributions and ancient DNA damage patterns for all hits are available as part of Supplementary dataset S1 (available for review at https://www.dropbox.com/scl/fi/60qnclu515egbkl8kbklr/fig_infections_nm_dmg.pdf?rlkey=9xyoq548l0capbwamapdlf3o5&dl=0 ).

##### *Borrelia recurrentis*

A total of 34 ancient microbial hits were assigned to *Borrelia recurrentis*, represented in the database by a single genome from strain A1 (assembly accession GCF_000019705.1). The strain A1 genome assembly consists of a linear chromosome with ~930 kb length, and seven linear plasmids ranging from 6.1 kb to 124 kb in length. Examination of read mapping distributions showed that each of the ancient *B. recurrentis* hits showed expected breadth of coverage across the large chromosome and multiple plasmids (Fig. S4.1). The highest rates of dropout at plasmid contig mappings were observed for hits with the lowest number of reads mapped, as expected (e.g. NEO29, 2/8 contigs covered).

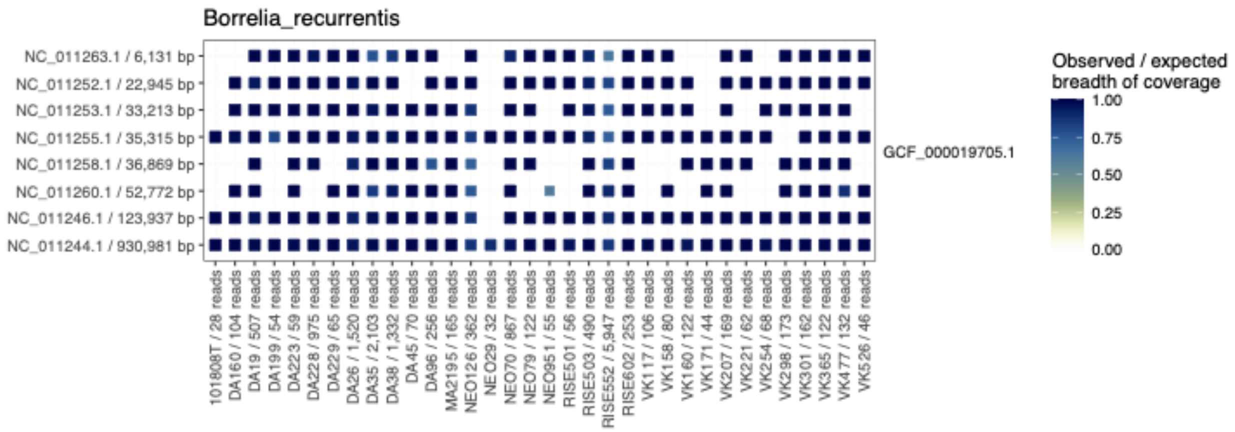


**Fig. S4.1 Genomic coverage for *B. recurrentis* hits.** Heatmap showing the number of reads mapped to the *B. recurrentis A1* reference assembly contigs for N=34 *B. recurrentis* hits. Symbol color and size indicate the ratio of observed over expected breadth of coverage.

The distribution of genomic similarity as measured by ANI across different species within the genus Borrelia followed patterns based on the phylogenetic relationships of the species[31] , with highest ANI observed for *B. recurrentis* and the two closely related species *B. duttoni* and *B. crocidurae* (Fig. S4.2).

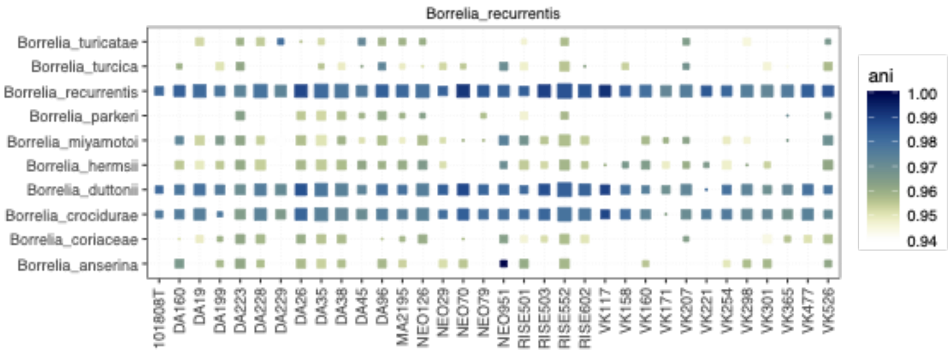


**Fig. S4.2 Genomic similarity for *B. recurrentis* hits.** Heatmap showing ANI of reads mapped to assemblies of different species within the genus *Borrelia*, for N=34 *B. recurrentis* hits. Symbol color and size indicate ANI and numbers of unique *k*-mers assigned to species in *krakenUniq*, respectively.

#### *Corynebacterium diphtheriae*

We identified two ancient hits assigned to *Corynebacterium diphtheriae,* the causative agent of diphtheria. Both were mapped to the assembly of strain NCTC3529 (accession GCF_900638705.1). Despite the limited numbers of reads mapped (Sidelkino, N=298 reads; VK408, N=338 reads), both hits showed clear evidence for authenticity based on ancient DNA damage as well as edit distance distributions (Fig. S4.3).

The assembly of strain NCTC3529 carries the gene for the diphtheria toxin responsible for diphtheria when present in *C. diphtheriae* and to a lesser extent in related *Corynebacterium* species (*tox*, NZ_LR134538.1:1,942,803-1,944,485), however due to the low read counts its presence in the two ancient hits could not be determined.

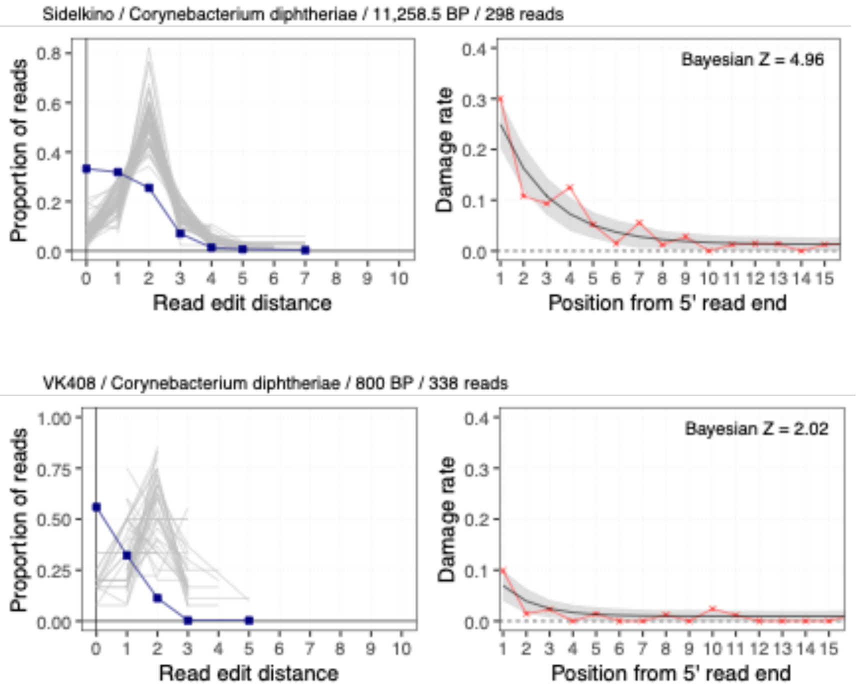


**Fig. S4.3 Read edit distance distributions and ancient DNA damage patterns for *C. diphtheriae* hits.** (left) Blue line and squares indicate edit distance for reads mapped to *C. diphtheriae,* whereas grey lines show mapping to other species within the *Corynebacterium* genus. (right) Observed nucleotide misincorporation frequencies (red symbols and line) and *metaDMG* fit (black line) and 68% credible intervals (shaded region) for C>T transitions as a function of distance from the 5' read end.

##### *Helicobacter pylori*

We identified three ancient hits assigned to *Helicobacter pylori,* each of which was mapped to a different reference assembly (Fig. S4.4). Despite the low overall number of mapped reads, the two hits with >100 reads mapped (RISE501, N=114 reads; Sidelkino, N=146 reads) showed expected breadth of coverage at the main chromosome as well as associated plasmids of the respective assemblies (Fig. S4.4)

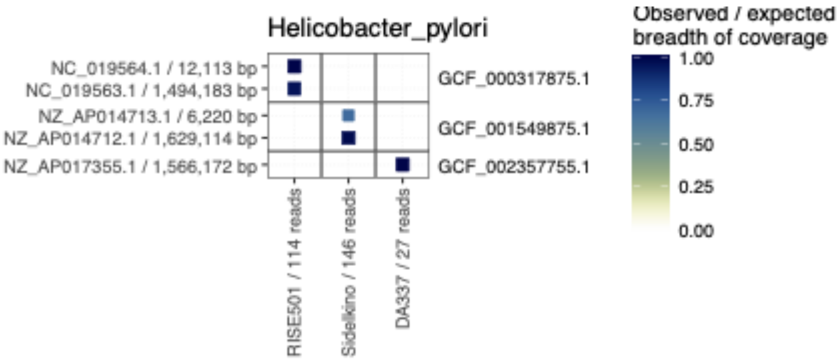


**Fig. S4.4 Genomic coverage for *H. pylori* hits.** Heatmap showing the number of reads mapped to the *H. pylori* reference assemblies, for N=3 *H. pylori* hits. Symbol color and size indicate the ratio of observed over expected breadth of coverage.

The distribution of genomic similarity across different species within the genus showed the highest ANI for *H. pylori*, consistent with authentication criteria (Fig. S4.5).

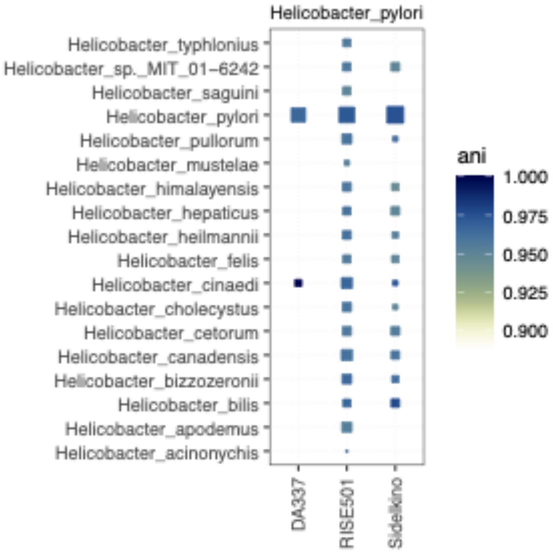


**Fig. S4.5 Genomic similarity for *H. pylori* hits.** Heatmap showing ANI of reads mapped to assemblies of different species within the genus Heliobacter, for N=3 *H. pylori* hits. Symbol color and size indicate ANI and numbers of unique *k*-mers assigned to species in *krakenUniq*, respectively.

##### *Leptospira spp.*

A total of 25 ancient microbial hits were assigned to *Leptospira spp.*, out of which five were *L. borgpetersenii* (two distinct assemblies), and 20 were *L. interrogans* (four distinct assemblies). All hits across both species showed expected breadth of coverage at the main chromosomes (Fig. S4.6, S4.7).

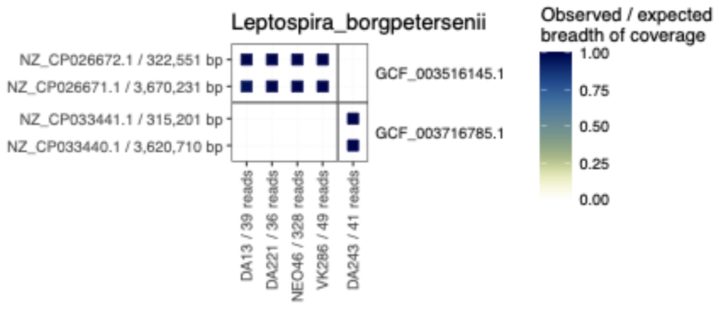


**Fig. S4.6 Genomic coverage for *L. borgpetersenii* hits.** Heatmap showing the number of reads mapped to the *L. borgpetersenii* reference assemblies, for N=5 *L. borgpetersenii* hits. Symbol color and size indicate the ratio of observed over expected breadth of coverage.

Most of the 20 hits within *L. interrogans* were mapped to two different strains, *L. interrogans* serovar Linhai str. 56609 (N=10; accession GCF_000941035.1) and *L. interrogans* strain FMAS_AW1 (N=8; accession GCF_005222625.1). For both reference assemblies, hits with higher read counts also showed coverage at the associated plasmids (e.g. VK467, VK279).

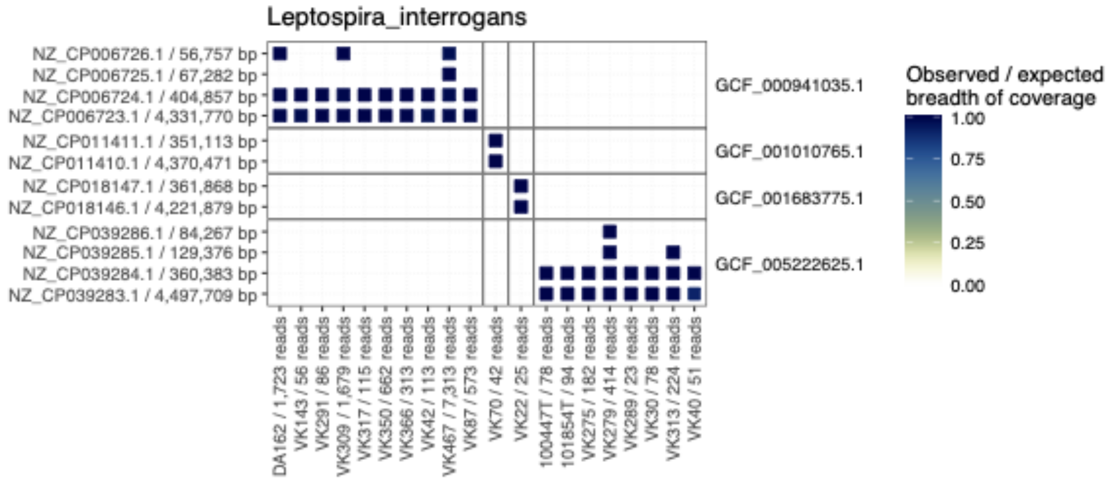


**Fig. S4.7 Genomic coverage for *L. interrogans* hits.** Heatmap showing the number of reads mapped to the *L. interrogans* reference assemblies, for N=20 *L. interrogans* hits. Symbol color and size indicate the ratio of observed over expected breadth of coverage.

The distribution of genomic similarity across different species within the genus showed the highest ANI for the respective *Leptospira* species, consistent with authentication criteria (Fig. S4.8).

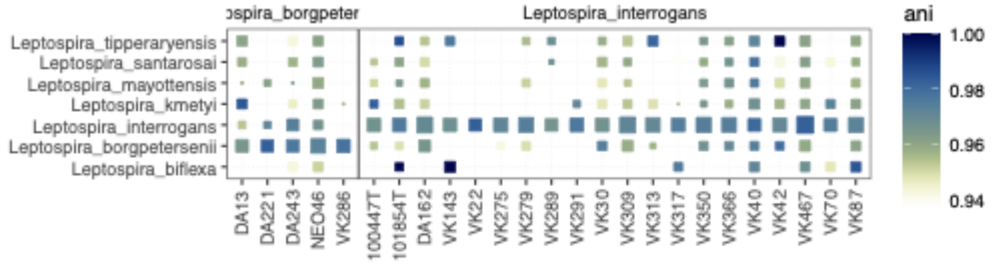

**Fig. S4.8 Genomic similarity for *Leptospira spp.* hits.** Heatmap showing ANI of reads mapped to assemblies of different species within the genus *Leptospira*, for N=25 *Leptospira spp.* hits. Symbol color and size indicate ANI and numbers of unique *k*-mers assigned to species in *krakenUniq*, respectively.

#### *Salmonella enterica*

We identified four ancient hits assigned to *Salmonella enterica,* each of which was mapped to a different reference assembly (Fig. S4.9). The highest coverage hit DA92 was mapped to *S. enterica* strain RKS4594, a serovar Paratyphi C (accession GCF_000018385.1). Correspondingly, DA92 also showed coverage for the associated virulence plasmid pSPCV (accession NC_012124.1; Fig. S4.9). Notably, previously reported ancient Eurasian *S. enterica* strains have been shown to belong to a clade including modern serovar Paratyphi C[32].

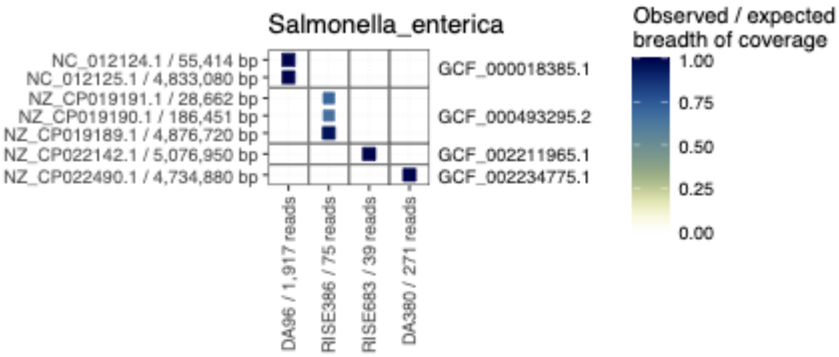


**Fig. S4.9 Genomic coverage for *S. enterica* hits.** Heatmap showing the number of reads mapped to the *S. enterica* reference assemblies, for N=4 *S. enterica* hits. Symbol color and size indicate the ratio of observed over expected breadth of coverage.

#### *Shigella spp.*

A total of 8 ancient microbial hits were assigned to *Shigella spp.*, out of which five were *S. boydii* (three distinct assemblies), three were *S. flexneri* (three distinct assemblies) and one was *S. dysenteriae* (Fig. S4.10-4.12).

The hits for *S. boydii* were mapped to reference assemblies of three different serotypes (serotype 9, strain ATCC 49812, accession GCF_002950135, serotype 8, strain CDC 3083-94, accesion GCF_000020185.1; serotype 11, strain 59-2708, GCF_002949495.1) and also showed coverage at plasmids associated with the respective strains for the three hits with >100 reads mapped (Fig. S4.10).

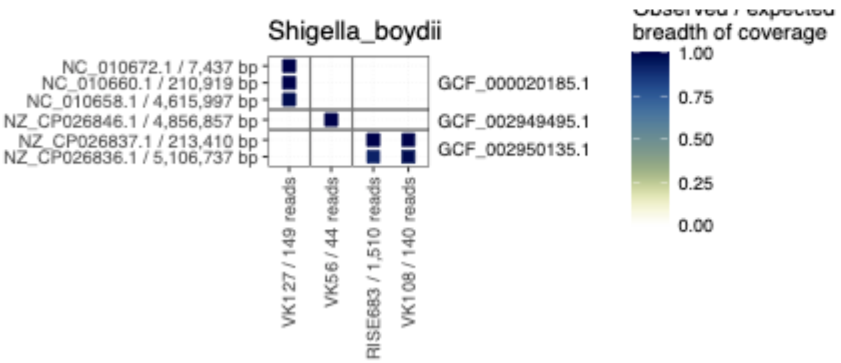


**Fig. S4.10 Genomic coverage for *S. boydii* hits.** Heatmap showing the number of reads mapped to the *S. boydii* reference assemblies, for N=4 *S. boydii* hits. Symbol color and size indicate the ratio of observed over expected breadth of coverage.

The hits for S. boydii and S. dysenteriae showed lower read numbers, but we did observe reads mapped to associated plasmids in two ouf the four cases (Figs. S4.11, S4.12)

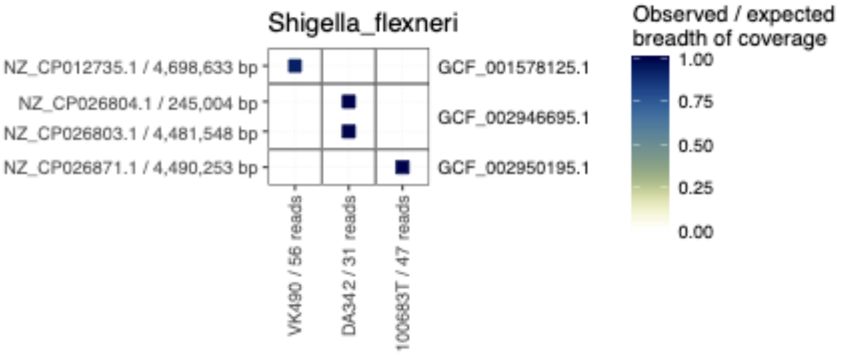


**Fig. S4.11 Genomic coverage for *S. flexneri* hits.** Heatmap showing the number of reads mapped to the *S. flexneri* reference assemblies, for N=4 *S. flexneri* hits. Symbol color and size indicate the ratio of observed over expected breadth of coverage.

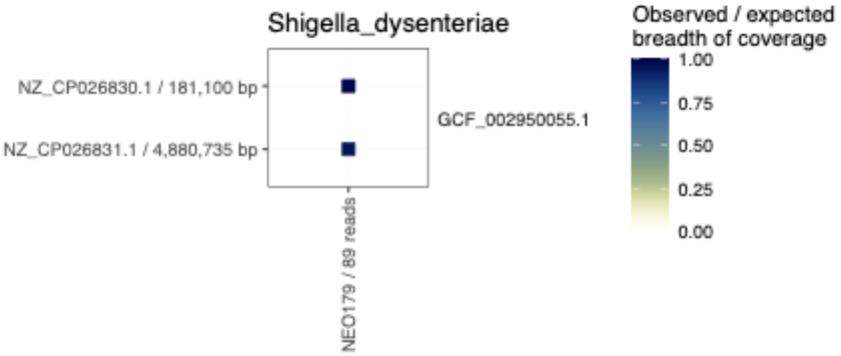


**Fig. S4.12 Genomic coverage for *S. dysenteriae* hits.** Heatmap showing the number of reads mapped to the *S. dysenteriae* reference assemblies, for N=4 *S. dysenteriae* hits. Symbol color and size indicate the ratio of observed over expected breadth of coverage.

##### *Treponema pallidum*

We identified three ancient hits assigned to *Treponema pallidum,* mapped to two different reference assemblies (Fig. S4.13). Due to the low read counts a confident assignment to different *T. pallidum* subspecies was not possible, although we note that the highest covered sample CGG_2_19555 (N=1,593 reads) showed highest unique *k*-mer count assigned to the syphilis causing *T. pallidum pallidum.*

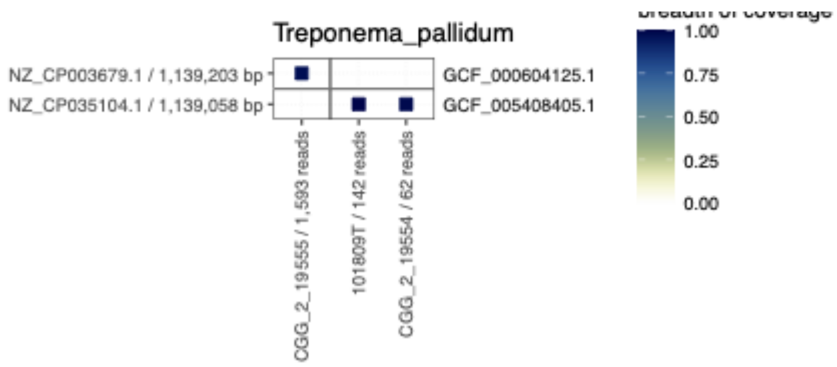


**Fig. S4.13 Genomic coverage for *T. pallidum* hits.** Heatmap showing the number of reads mapped to the *T. pallidum* reference assemblies, for N=3 *T. pallidum* hits. Symbol color and size indicate the ratio of observed over expected breadth of coverage

#### *Yersinia spp.*

A total of 54 ancient microbial hits were assigned to *Yersinia spp*, out of which 12 were *Y. enterocolitica* and 42 were *Y. pestis.* The distribution of genomic similarity across different species within the genus showed the highest ANI for the respective *Yersinia* species, consistent with our authentication criteria based on the highest number of *k*-mers assigned by krakenUniq (Methods; Fig. S4.14).

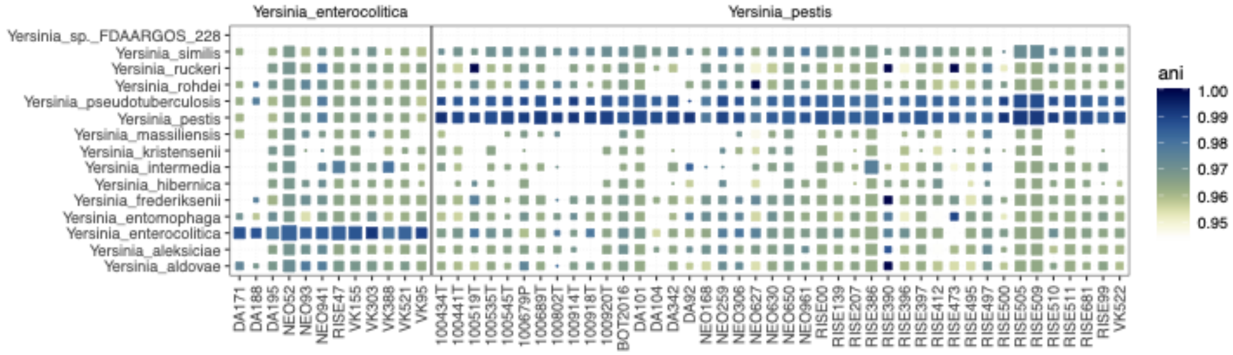


**Fig. S4.14 Genomic similarity for *Yersinia spp.* hits.** Heatmap showing ANI of reads mapped to assemblies of different species within the genus *Yersinia*, for N=54 *Yersinia spp.* hits. Symbol color and size indicate ANI and numbers of unique *k*-mers assigned to species in *krakenUniq*, respectively.

The ancient hits for *Y. enterocolitica* were all mapped to strain YE53/03 (accession GCF_000968115.1; Fig. S4.15), a biotype 1A strain isolated from a human fecal sample. *Y. enterocolitica* biotype 1A has historically been considered as non-pathogenic[33,34], although this view has been challenged by more recent evidence[33]. Strain YE53/03 does not carry the pYV virulence plasmid associated with pathogenic strains, which also appears absent in the highest covered ancient hit NEO52.

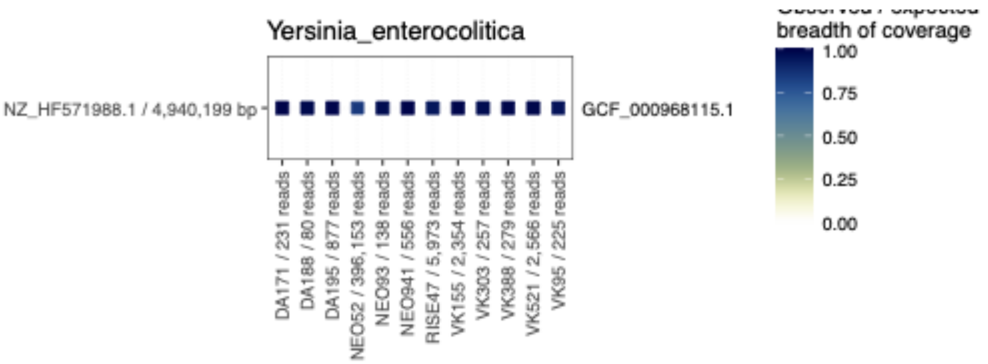


**Fig. S4.15 Genomic coverage for *Y. enterocolitica* hits.** Heatmap showing the number of reads mapped to the *Y. enterocolitica* reference assemblies, for N=12 *Y. enterocolitica* hits. Symbol color and size indicate the ratio of observed over expected breadth of coverage

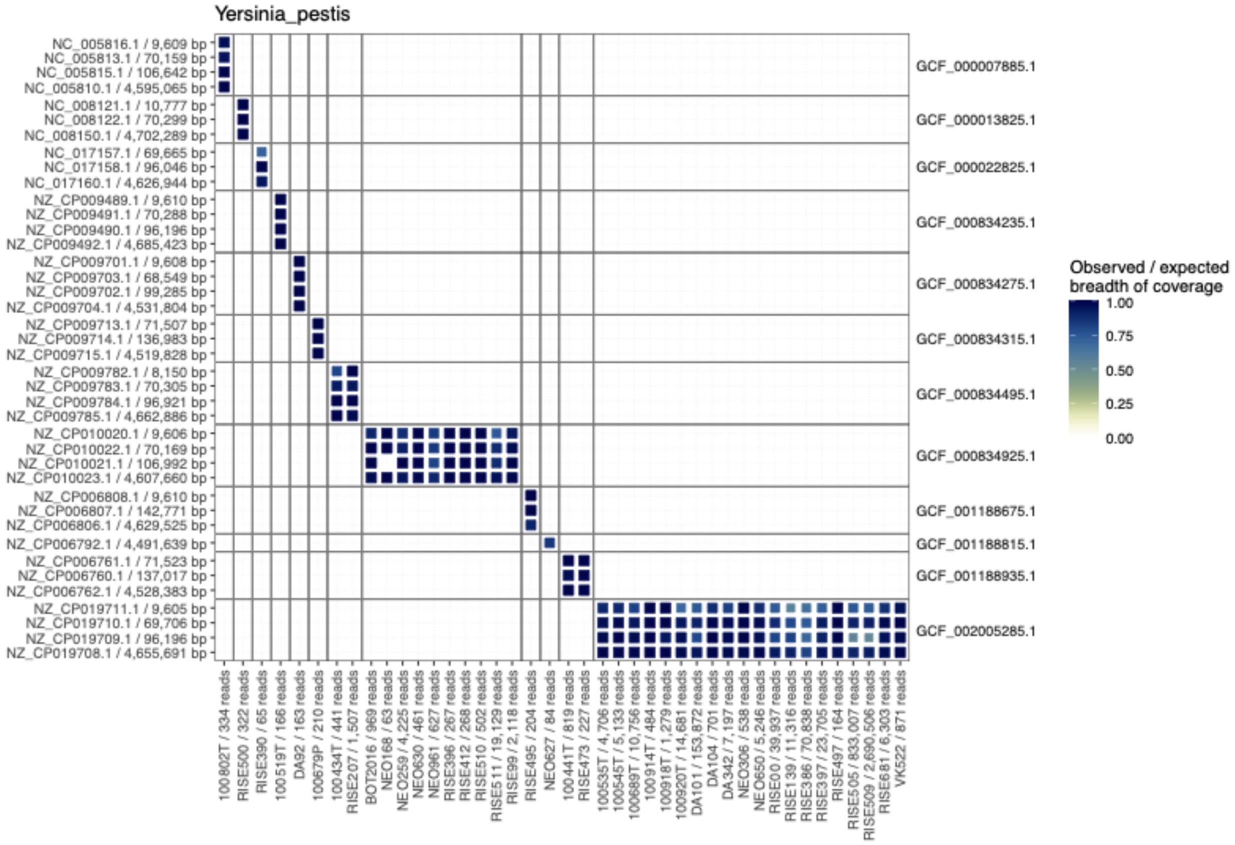


**Fig. S4.16 Genomic coverage for *Y. pestis* hits.** Heatmap showing the number of reads mapped to the *Y. pestis* reference assemblies, for N=42 *Y. pestis* hits. Symbol color and size indicate the ratio of observed over expected breadth of coverage

The ancient hits for *Y. pestis* were mapped to a variety of different reference assemblies, but generally showing coverage at the main chromosomes and  associated plasmids (Fig. S4.16). To further characterize the newly reported *Y. pestis* hits, we re-mapped the reads assigned to the genus of *Yersinia* in the classification step to the *Y. pestis* strain CO92 reference assembly (accession GCF_000009065.1). All hits (except sample NEO627 with a low read count) showed coverage patterns across the *Y. pestis* virulence plasmids pCD1 and pMT1 (Extended Data Fig. 7c).

To further investigate the evolutionary relationships of our ancient hits, we carried out phylogenetic placement of ancient hits with >0.01X read depth onto a phylogeny of known high-quality ancient and modern *Y. pestis* genomes, as described previously[35]. The obtained placements were consistent with sample age and context (Figs. S4.17-S4.32).

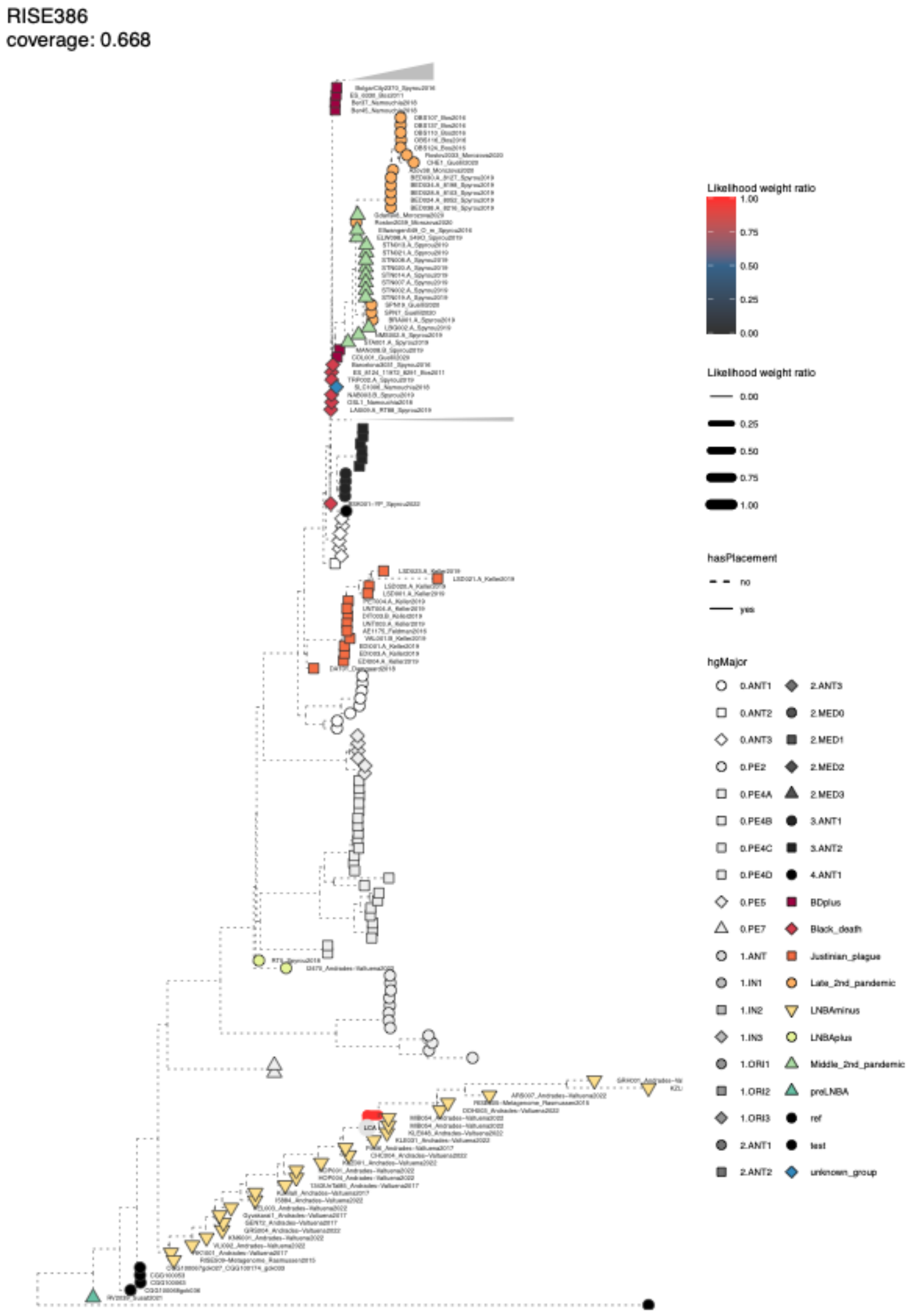


**Fig. S4.17 Phylogenetic placement for Y. pestis hit RISE386.** Phylogenetic tree showing the placement of a lower coverage sample using epa-ng. Branches with placement weights are indicated with a solid colored line, and the last common ancestor of all placement branches is indicated with LCA.

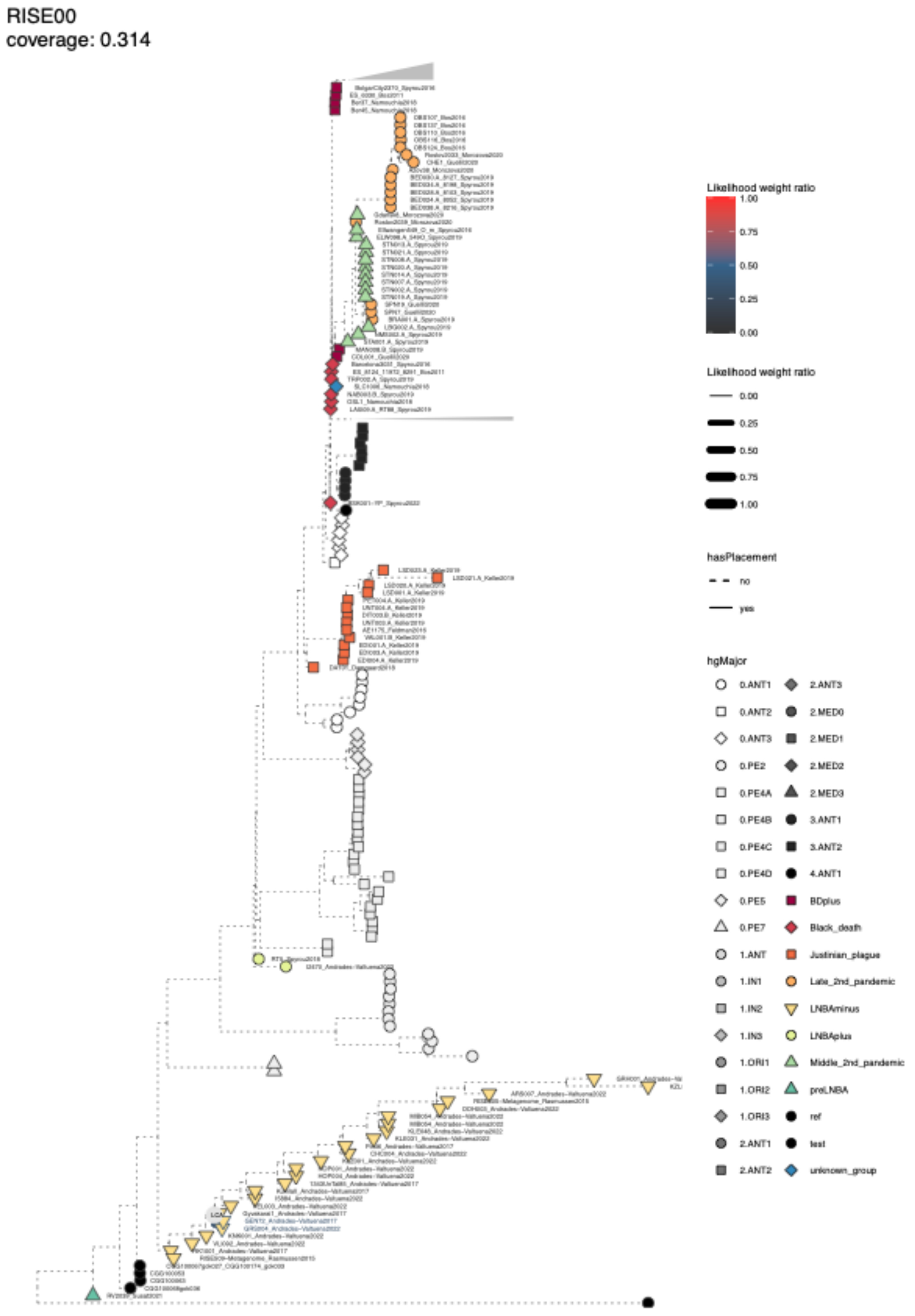


**Fig. S4.18 Phylogenetic placement for Y. pestis hit RISE00.** Phylogenetic tree showing the placement of a lower coverage sample using epa-ng. Branches with placement weights are indicated with solid colored line, and the last common ancestor of all placement branches is indicated with LCA.

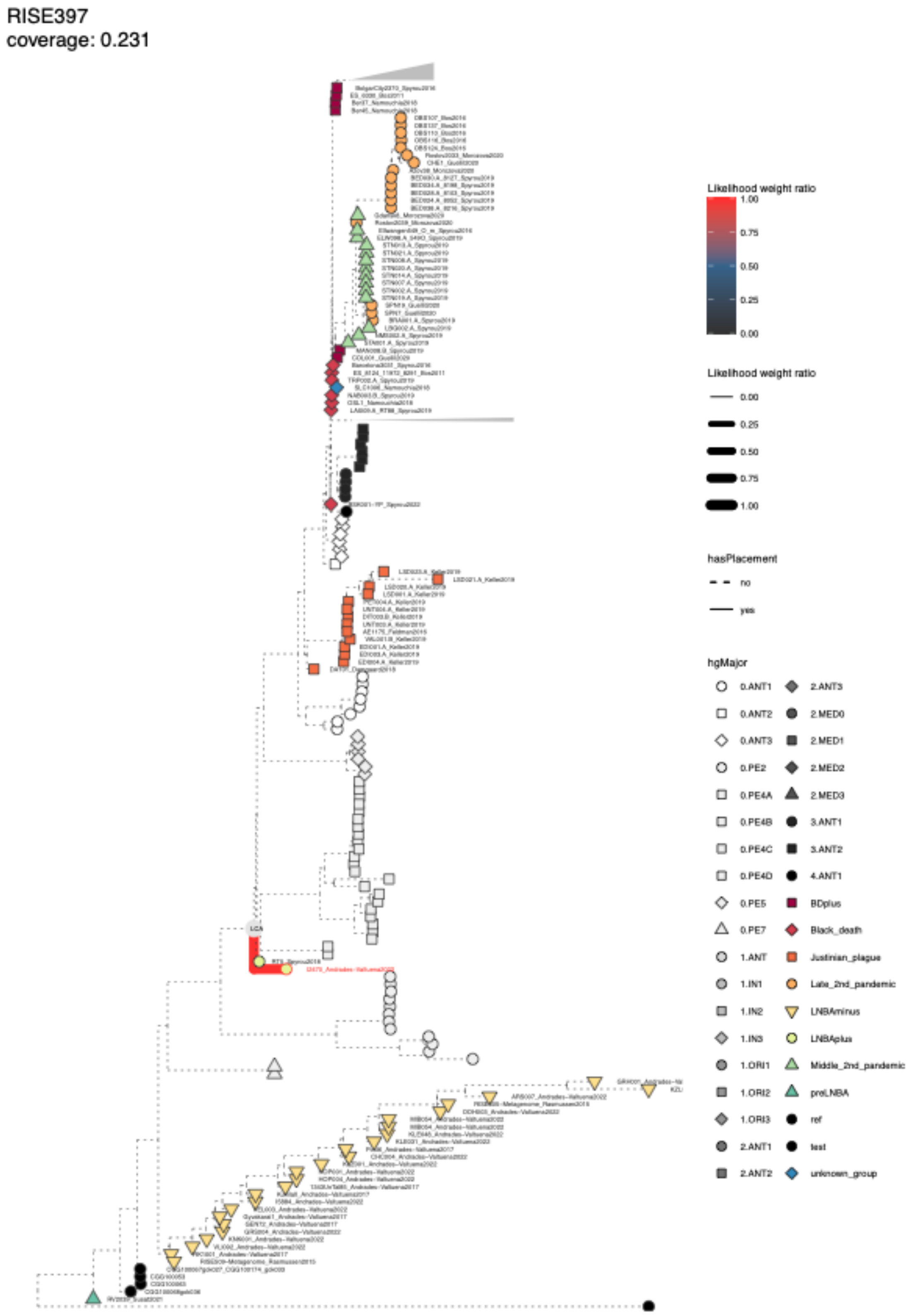


**Fig. S4.19 Phylogenetic placement for Y. pestis hit RISE397.** Phylogenetic tree showing the placement of a lower coverage sample using epa-ng. Branches with placement weights are indicated with solid colored lines, and the last common ancestor of all placement branches is indicated with LCA.

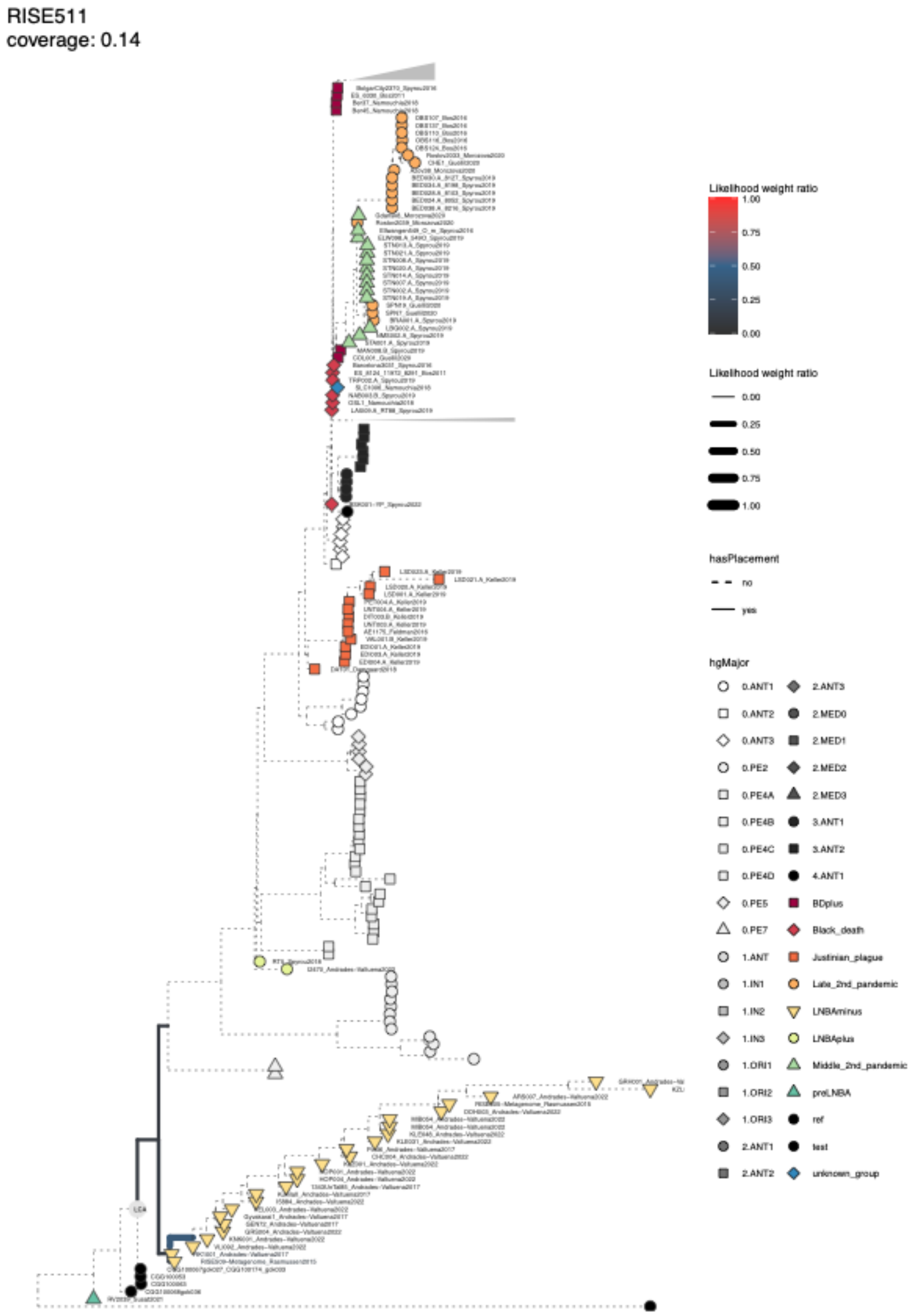


**Fig. S4.20 Phylogenetic placement for Y. pestis hit RISE511.** Phylogenetic tree showing the placement of a lower coverage sample using epa-ng. Branches with placement weights are indicated with solid colored lines, and the last common ancestor of all placement branches is indicated with LCA.

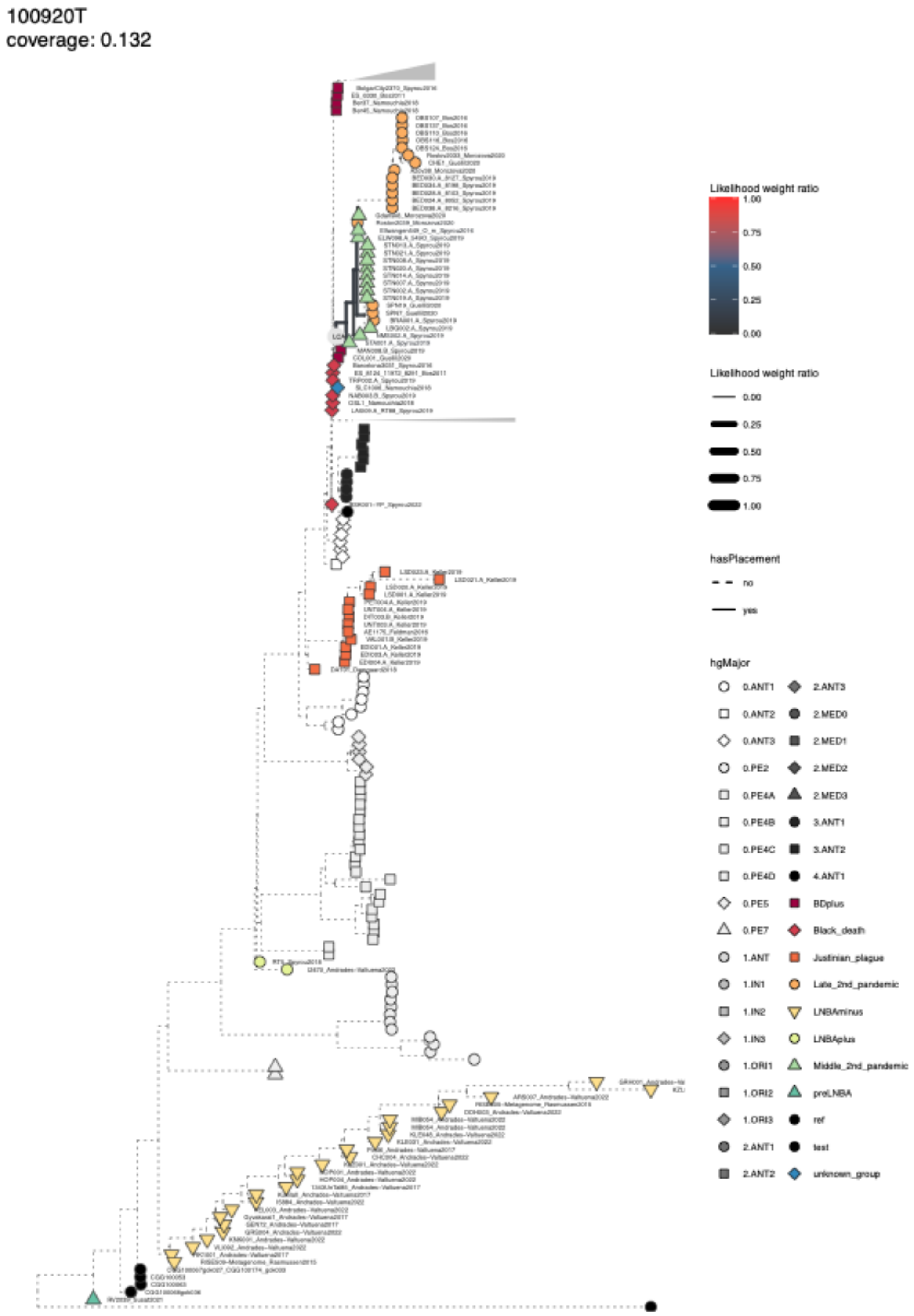


**Fig. S4.21 Phylogenetic placement for Y. pestis hit 100920T.** Phylogenetic tree showing the placement of a lower coverage sample using epa-ng. Branches with placement weights are indicated with solid colored lines, and the last common ancestor of all placement branches is indicated with LCA.

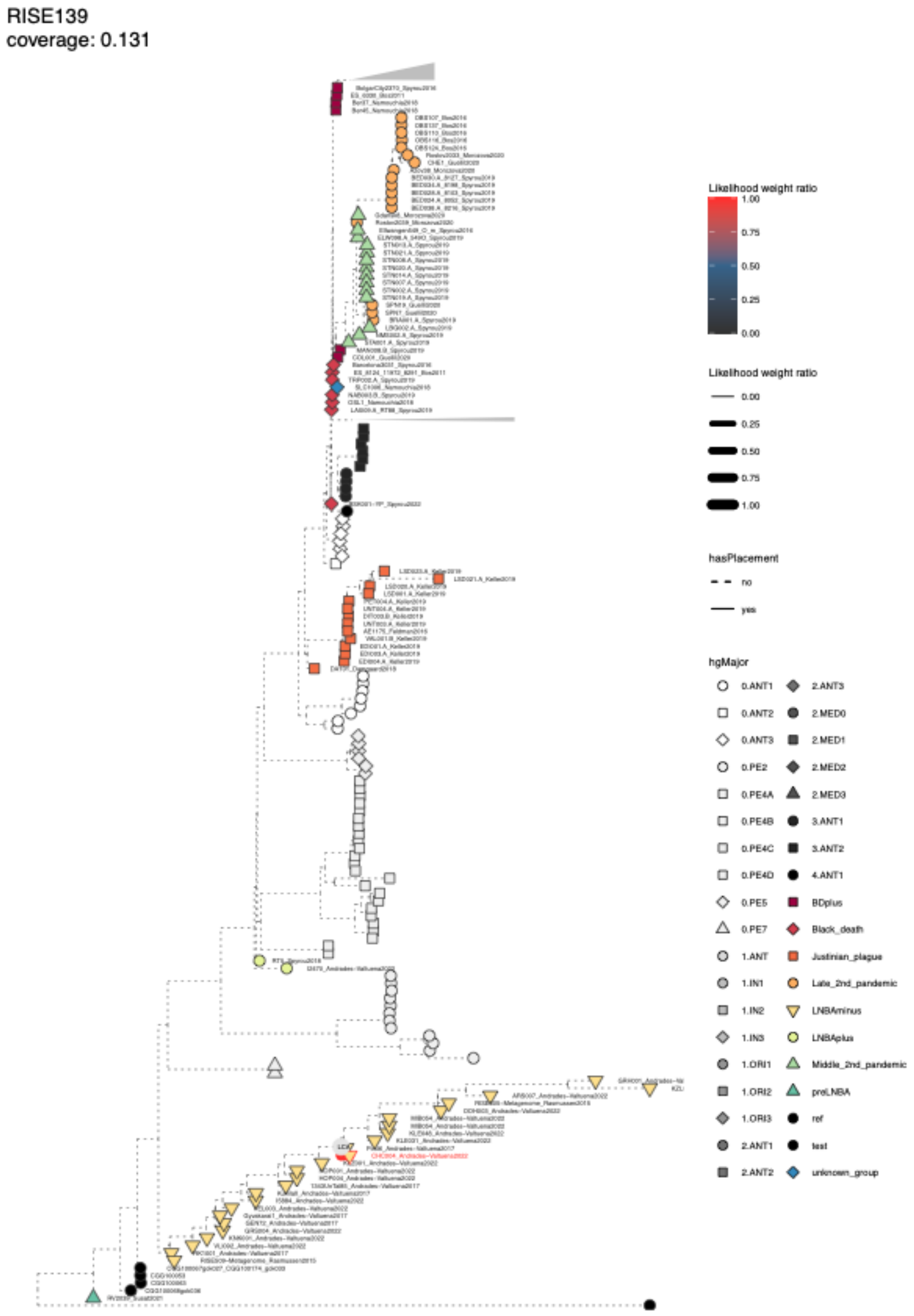


**Fig. S4.22 Phylogenetic placement for Y. pestis hit RISE139.** Phylogenetic tree showing the placement of a lower coverage sample using epa-ng. Branches with placement weights are indicated with a solid colored line, and the last common ancestor of all placement branches is indicated with LCA.

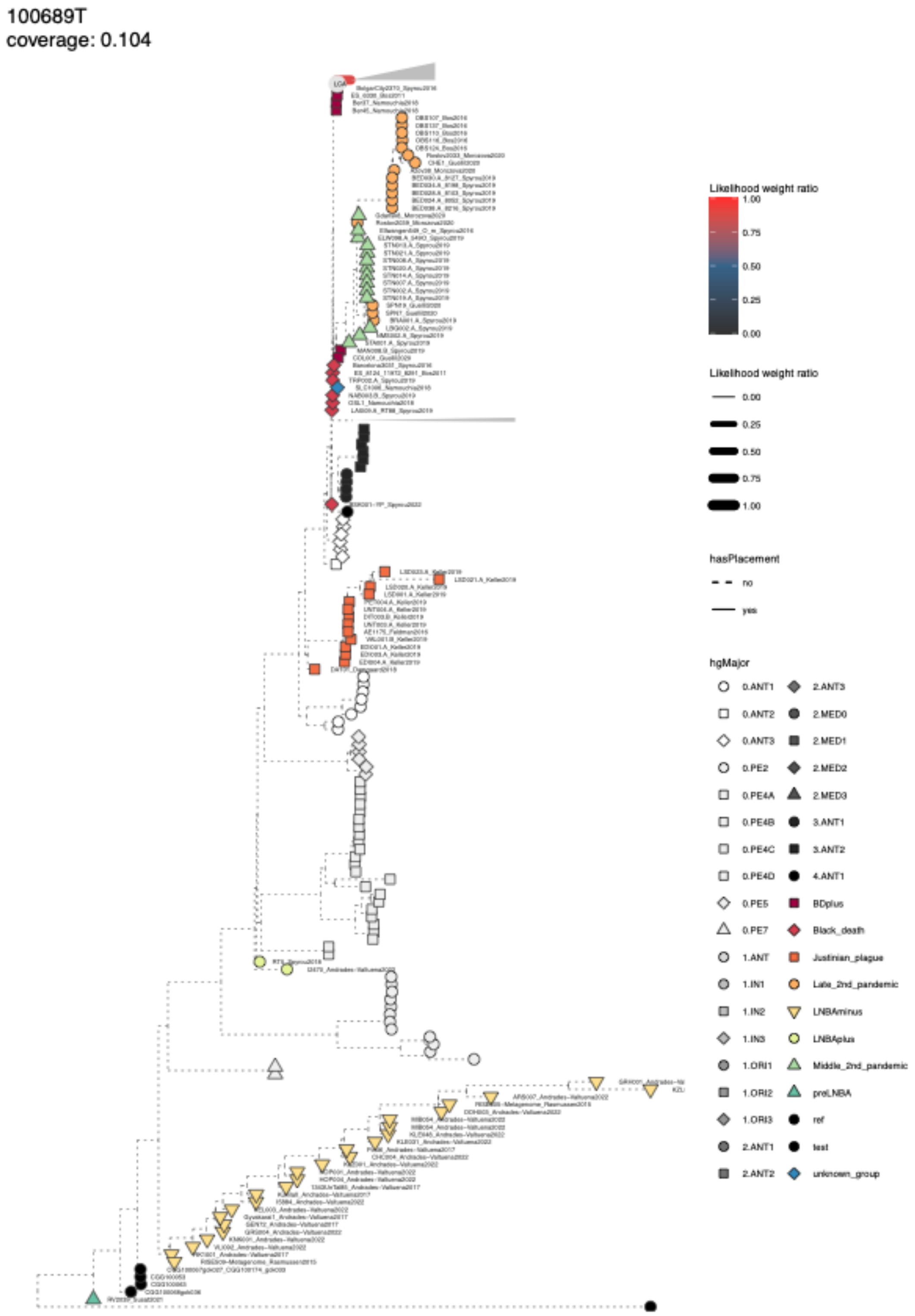


**Fig. S4.23 Phylogenetic placement for Y. pestis hit 100689T.** Phylogenetic tree showing the placement of a lower coverage sample using epa-ng. Branches with placement weights are indicated with a solid colored line, and the last common ancestor of all placement branches is indicated with LCA.

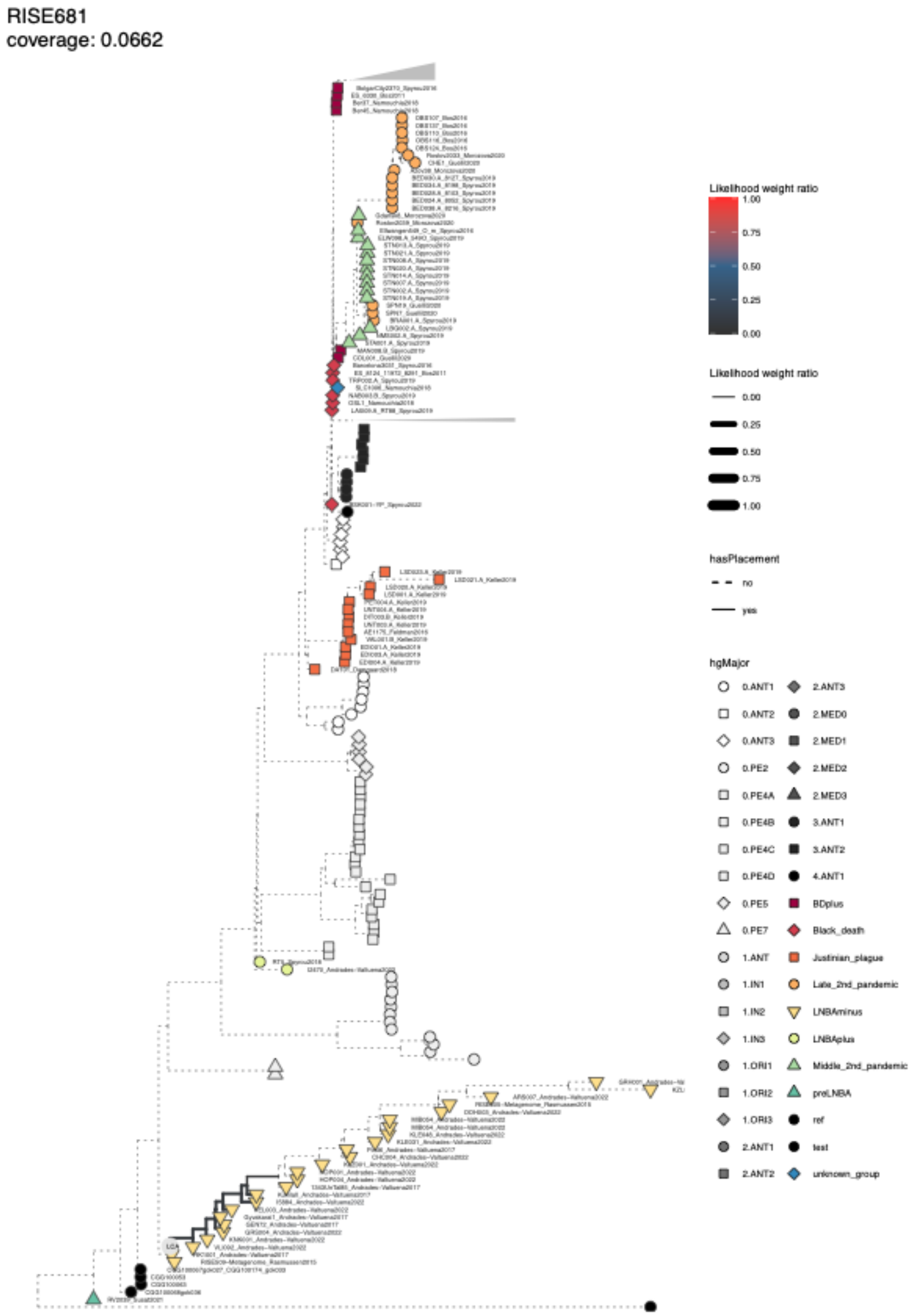


**Fig. S4.24 Phylogenetic placement for Y. pestis hit RISE681.** Phylogenetic tree showing the placement of a lower coverage sample using epa-ng. Branches with placement weights are indicated with solid colored lines, and the last common ancestor of all placement branches is indicated with LCA.

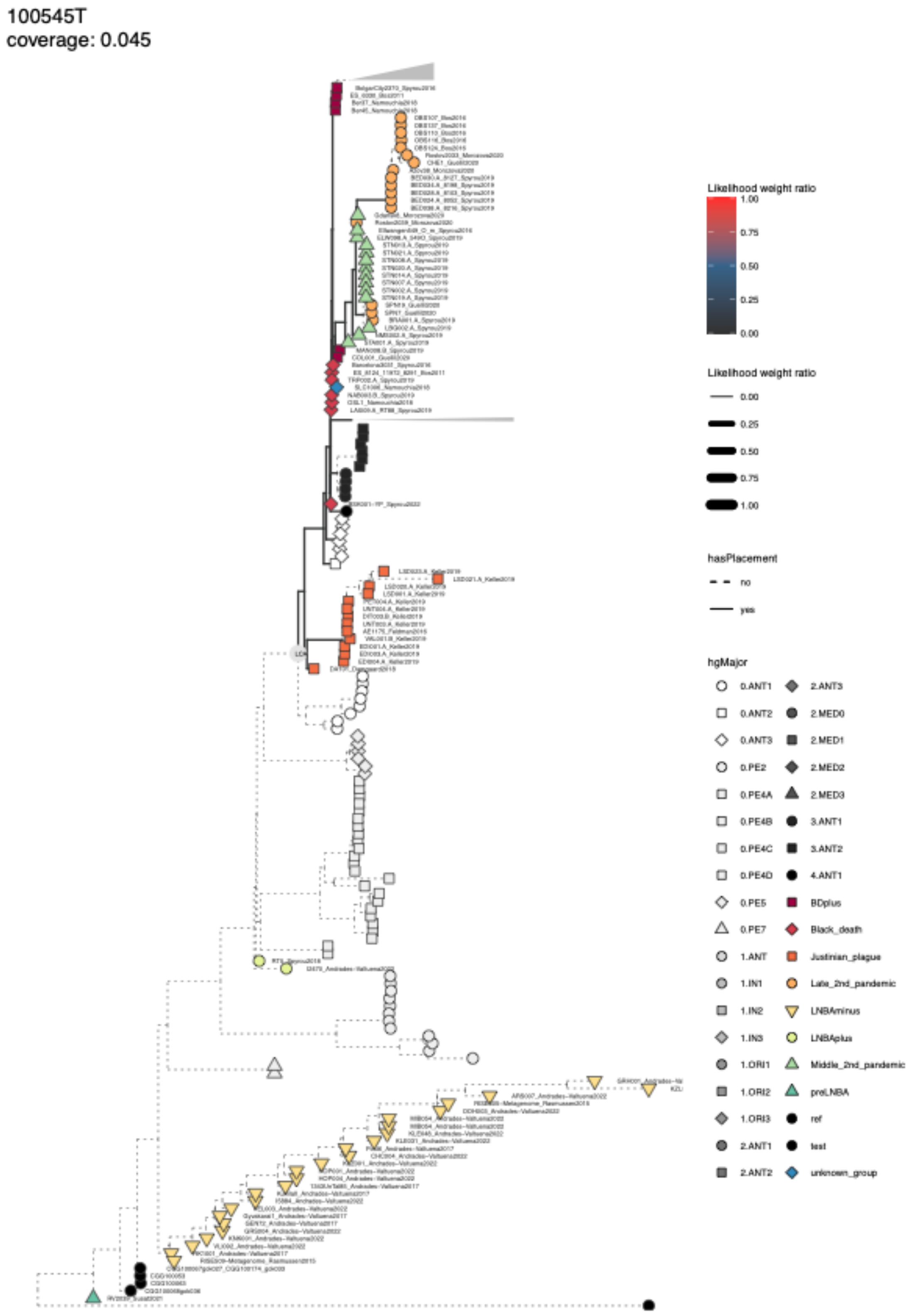


**Fig. S4.25 Phylogenetic placement for Y. pestis hit 100545T.** Phylogenetic tree showing the placement of a lower coverage sample using epa-ng. Branches with placement weights are indicated with solid colored lines, and the last common ancestor of all placement branches is indicated with LCA.

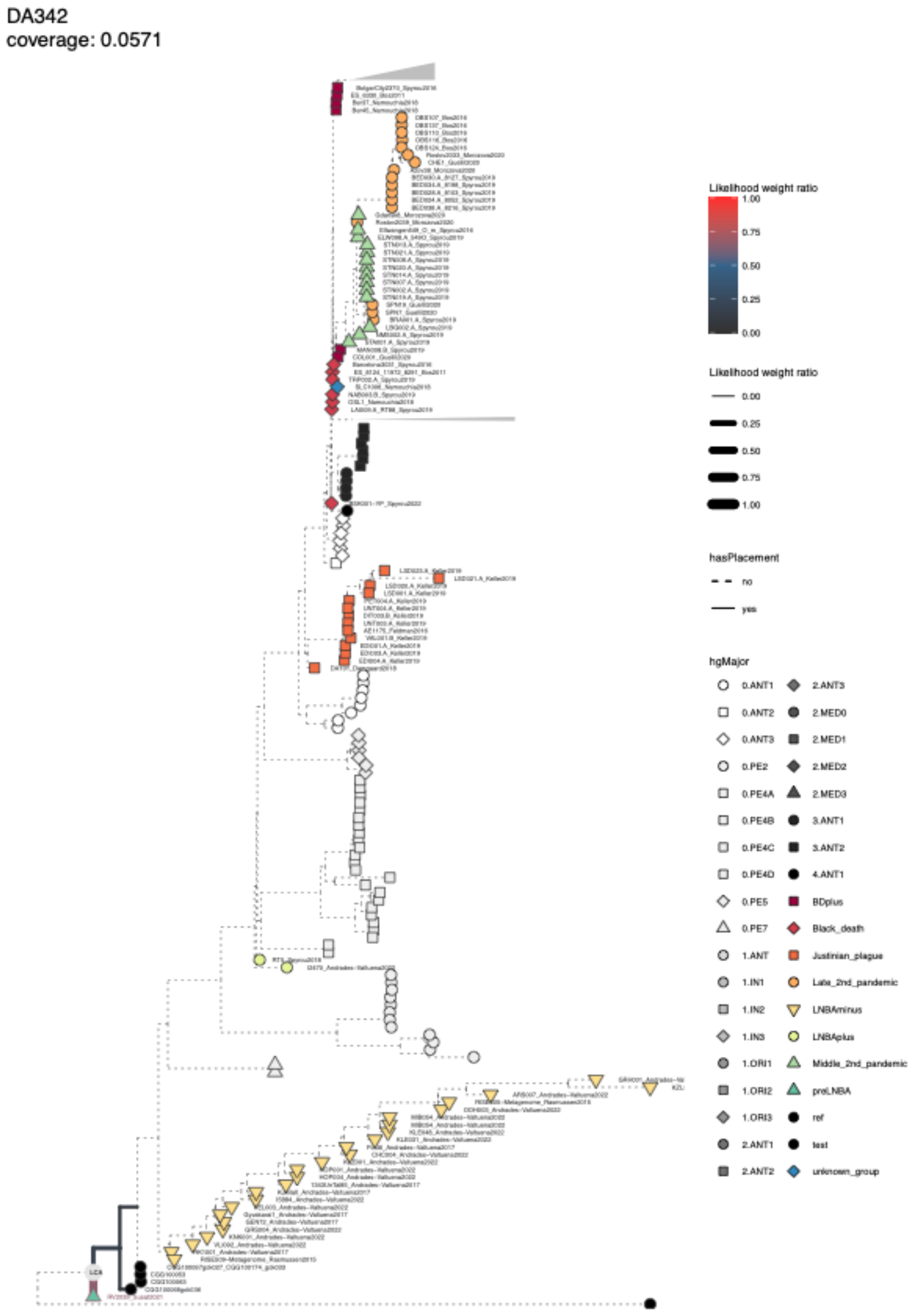


**Fig. S4.26 Phylogenetic placement for Y. pestis hit DA342.** Phylogenetic tree showing the placement of a lower coverage sample using epa-ng. Branches with placement weights are indicated with solid colored lines, and the last common ancestor of all placement branches is indicated with LCA.

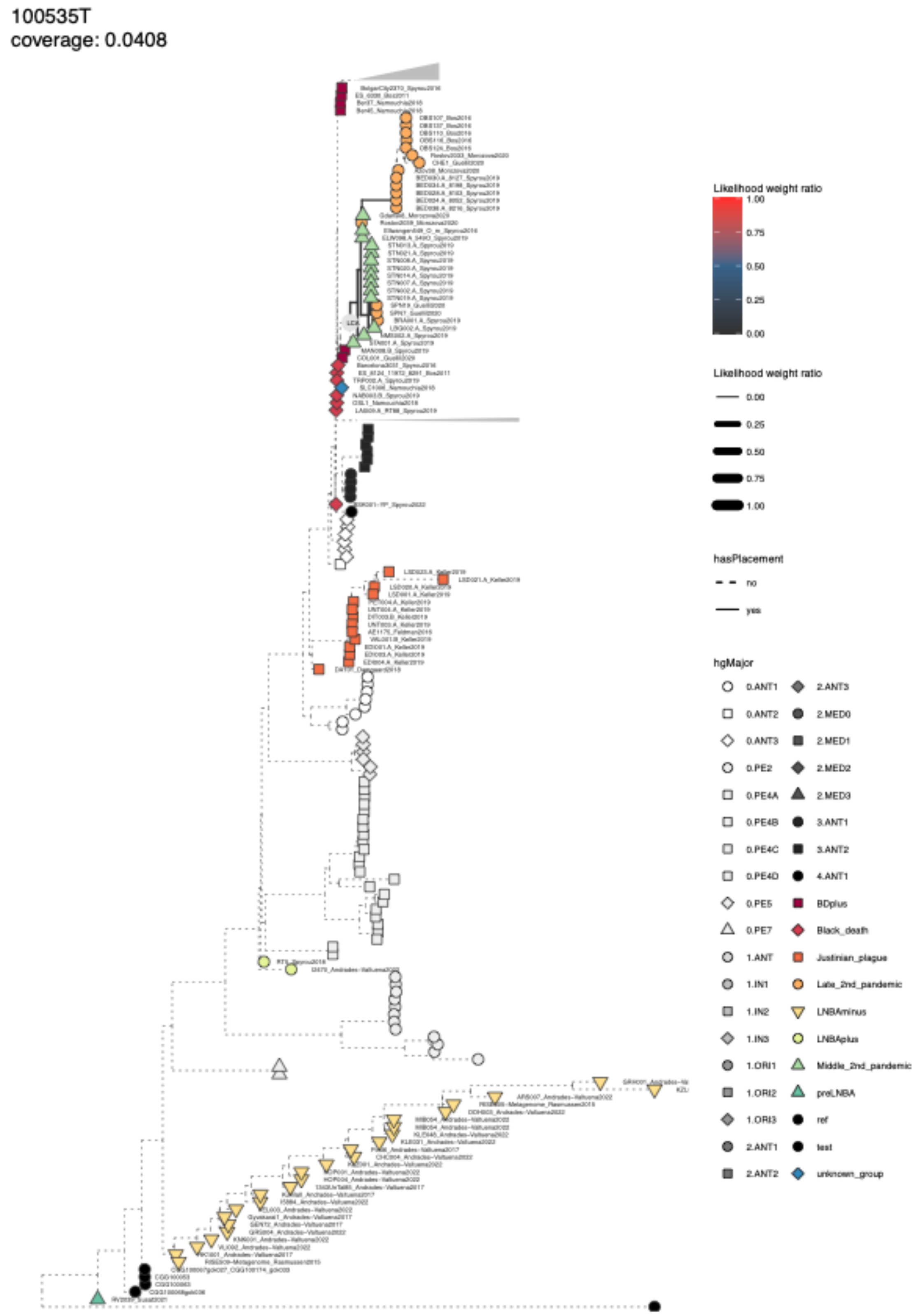


**Fig. S4.27 Phylogenetic placement for Y. pestis hit 100535T.** Phylogenetic tree showing the placement of a lower coverage sample using epa-ng. Branches with placement weights are indicated with solid colored lines, and the last common ancestor of all placement branches is indicated with LCA.

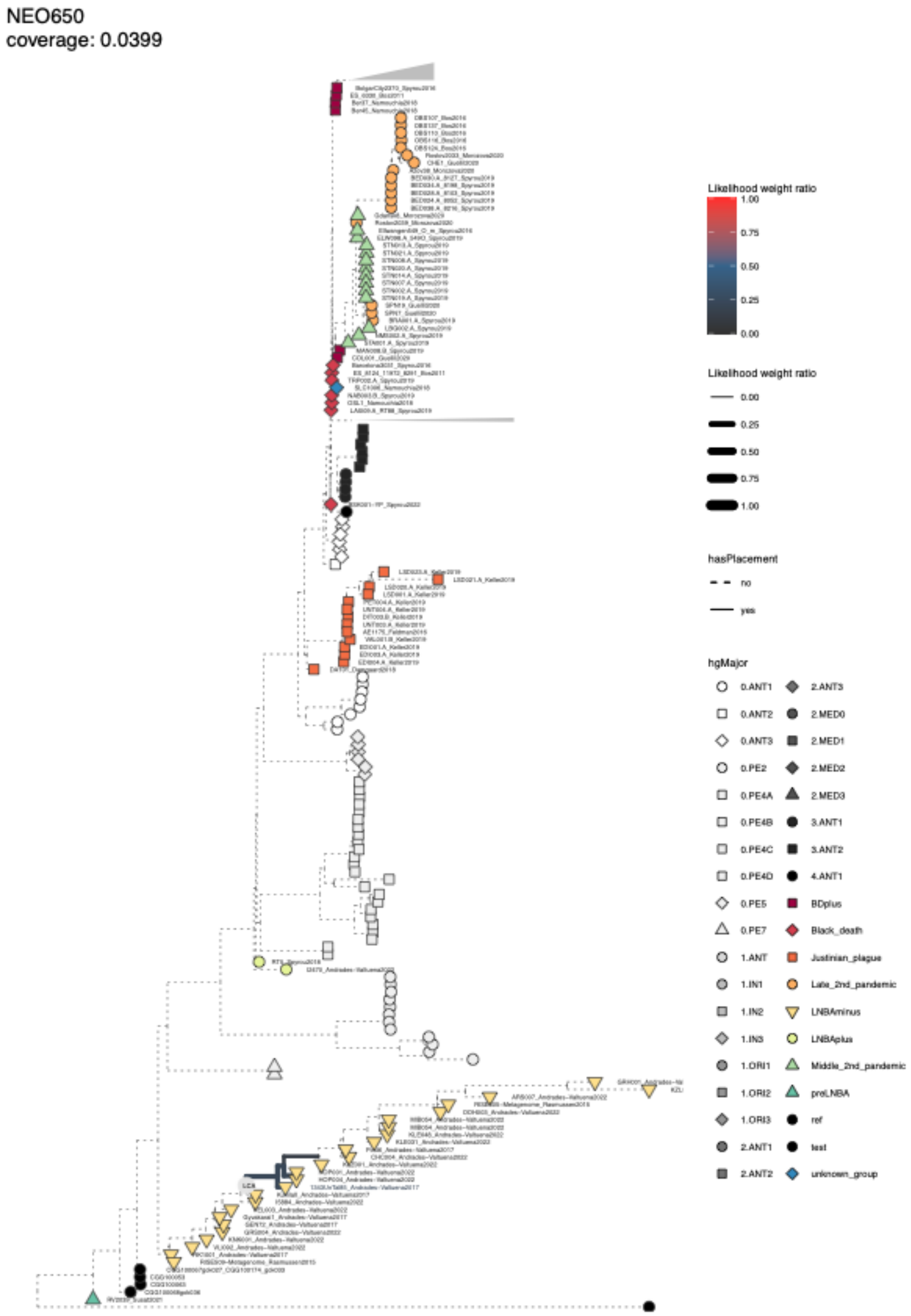


**Fig. S4.28 Phylogenetic placement for Y. pestis hit NEO650.** Phylogenetic tree showing the placement of a lower coverage sample using epa-ng. Branches with placement weights are indicated with solid colored lines, and the last common ancestor of all placement branches is indicated with LCA.

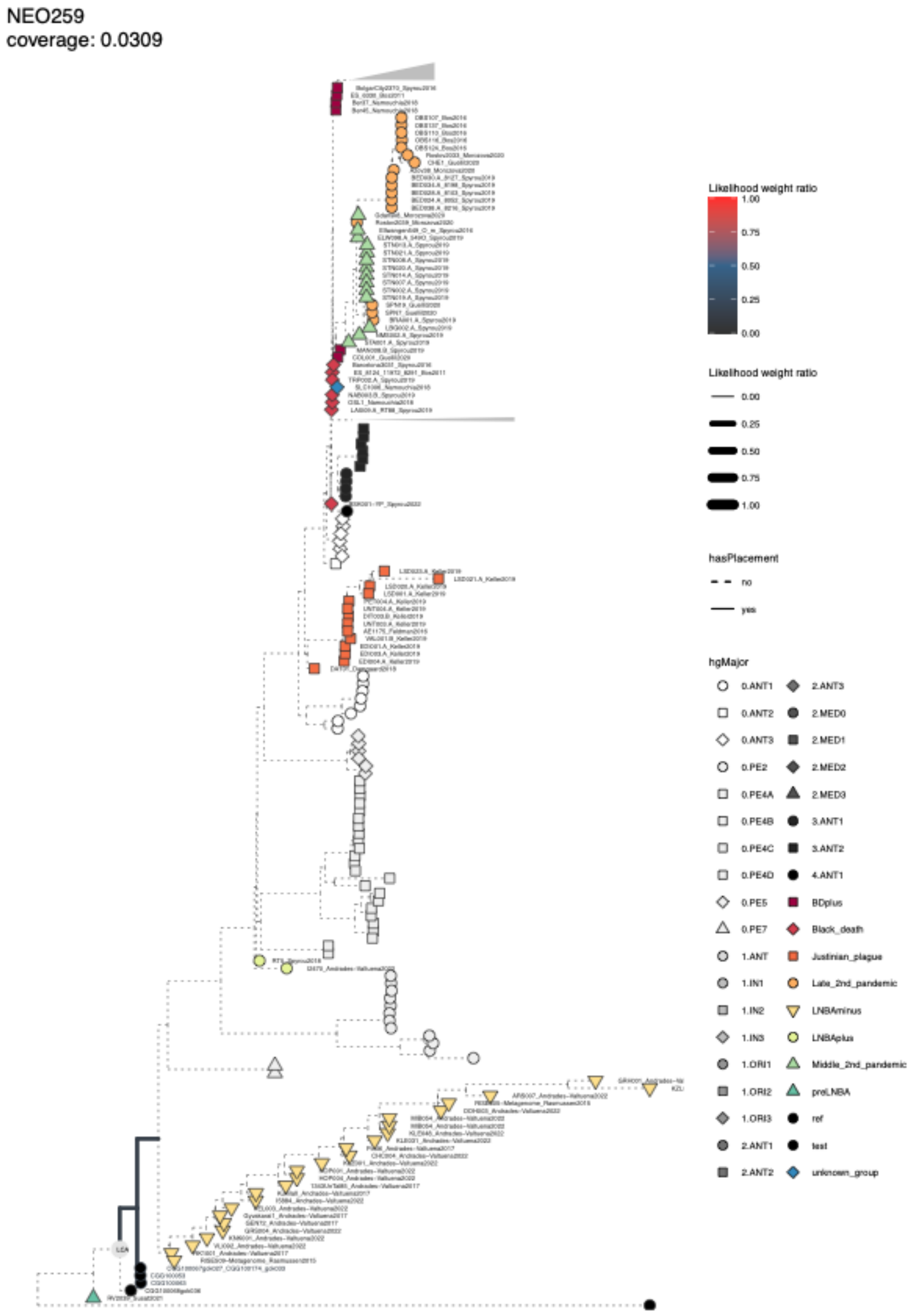


**Fig. S4.29 Phylogenetic placement for Y. pestis hit NEO259.** Phylogenetic tree showing the placement of a lower coverage sample using epa-ng. Branches with placement weights are indicated with solid colored lines, and the last common ancestor of all placement branches is indicated with LCA.

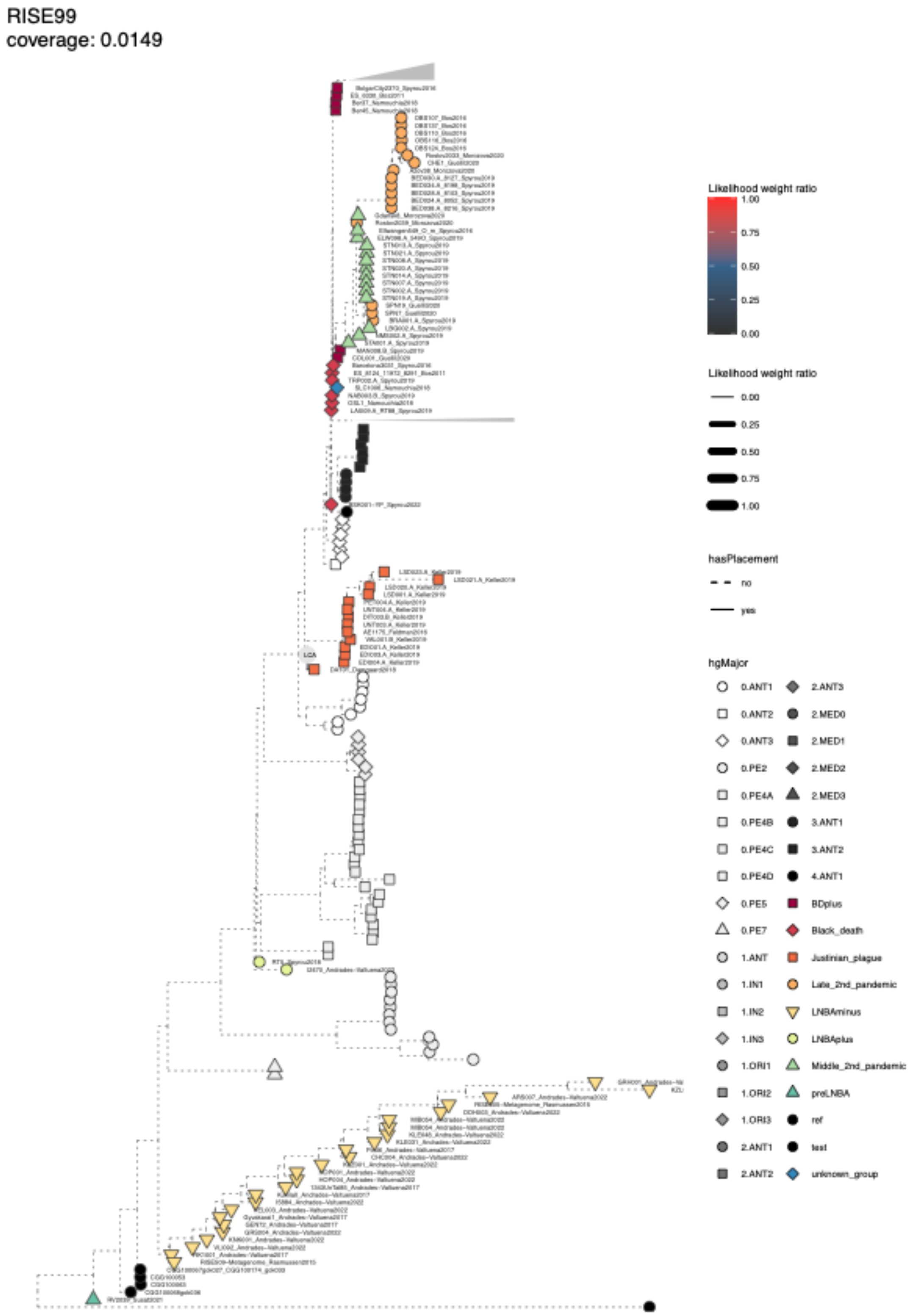


**Fig. S4.30 Phylogenetic placement for Y. pestis hit RISE99.** Phylogenetic tree showing the placement of a lower coverage sample using epa-ng. Branches with placement weights are indicated with solid colored line, and the last common ancestor of all placement branches is indicated with LCA.

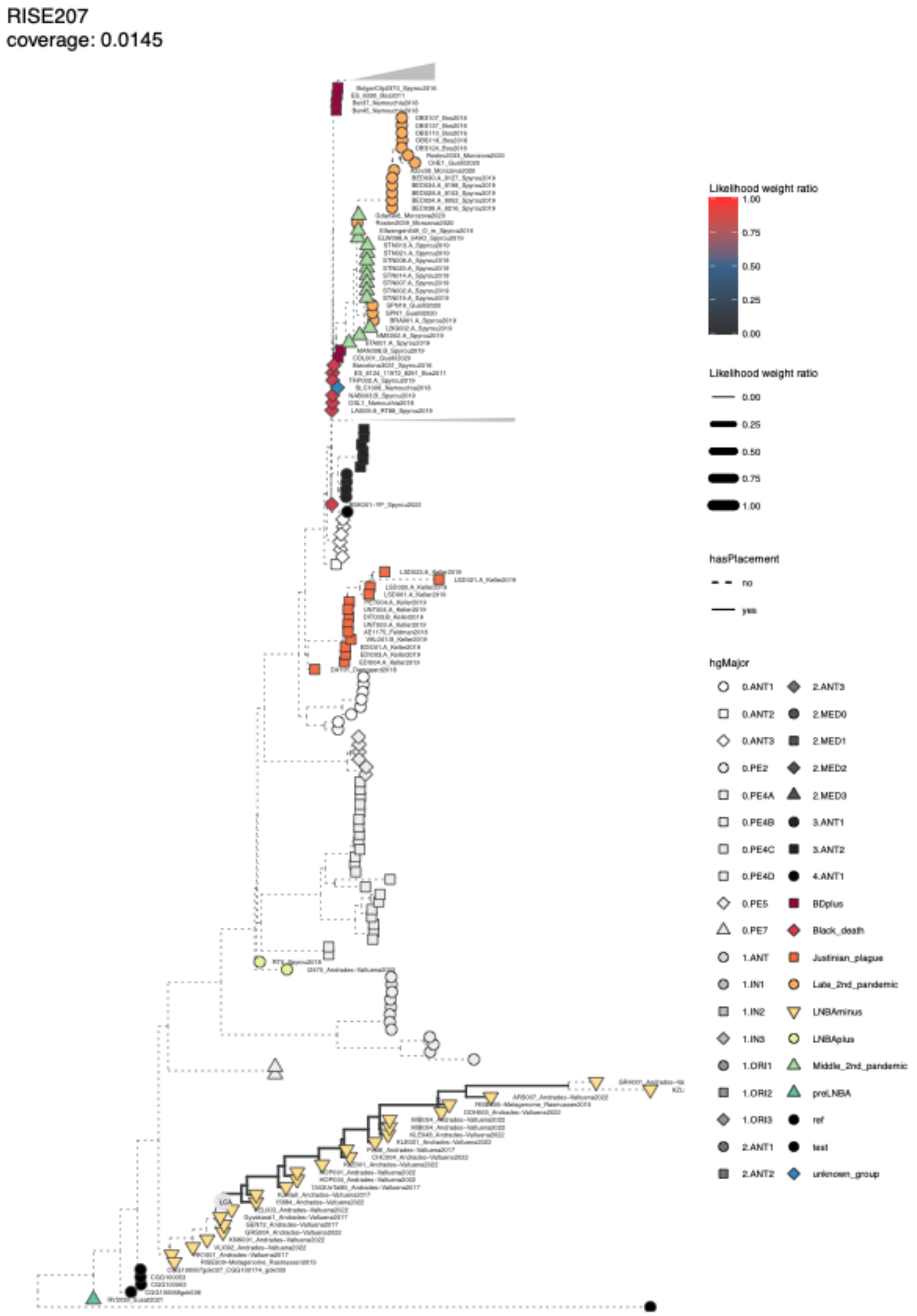


**Fig. S4.31 Phylogenetic placement for Y. pestis hit RISE207.** Phylogenetic tree showing the placement of a lower coverage sample using epa-ng. Branches with placement weights are indicated with solid colored lines, and the last common ancestor of all placement branches is indicated with LCA.

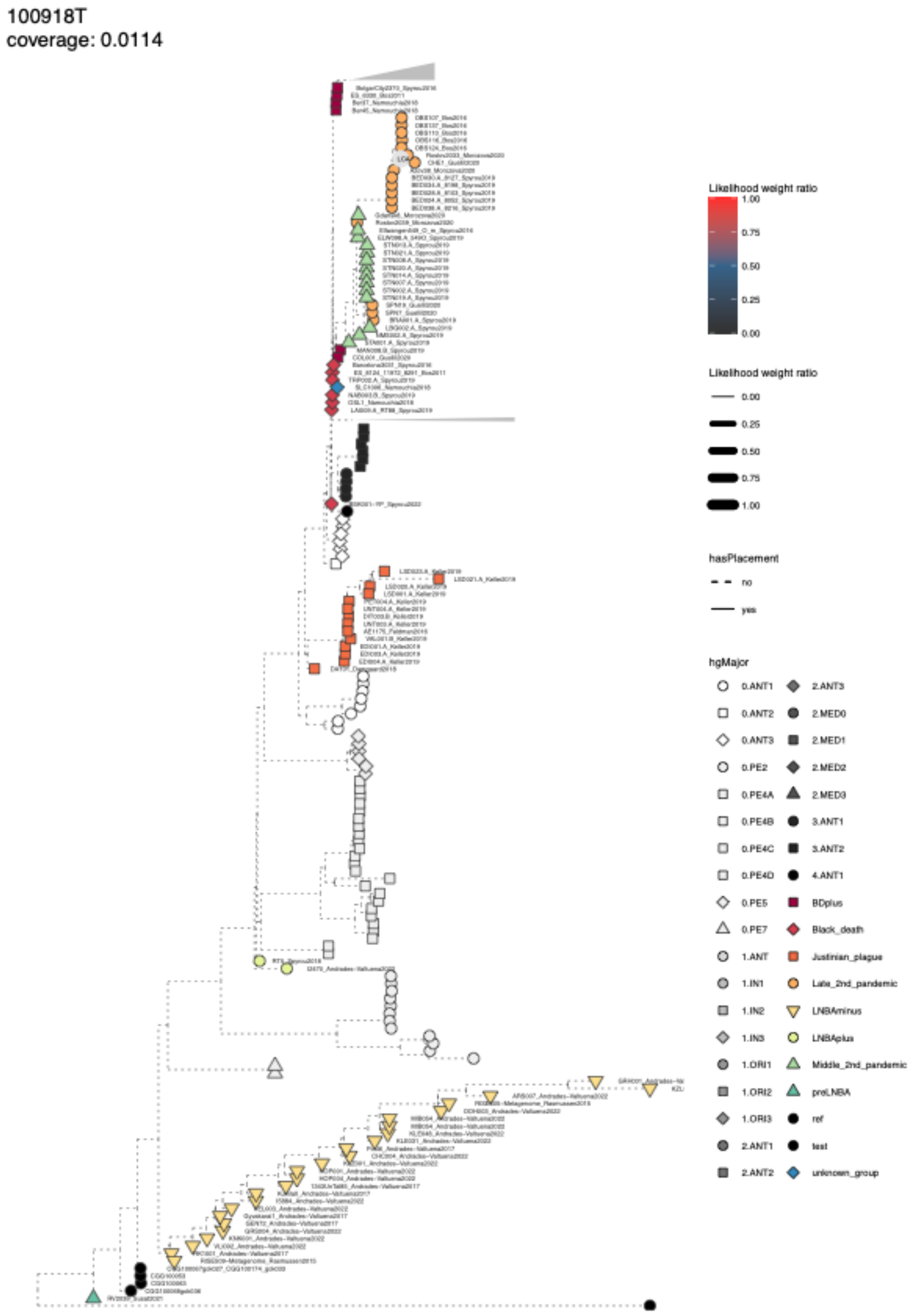


**Fig. S4.32 Phylogenetic placement for Y. pestis hit 100918T.** Phylogenetic tree showing placement of lower coverage sample using epa-ng. Branches with placement weights are indicated with a solid colored line, and the last common ancestor of all placement branches is indicated with LCA.

### Supplementary information 5: *Borrelia recurrentis* and louse-borne relapsing fever

#### Transmission of *Borrelia recurrentis*

Louse-borne relapsing fever (LBRF) is caused by the bacteria *Borrelia recurrentis*, a large, loosely coiled, motile spirochaete[36]. Human body lice (*Pediculus humanus humanus*) and perhaps also head lice (*P. humanus capitis*)[37] become infected in the midgut with *B. recurrentis* when feeding on infected people. Lice are obligate hematophagous human ectoparasites. They are intolerant of deviations in human body temperatures caused by fever, climatic exposure or death, and they can survive in clothing without feeding for approximately a month.

*B. recurrentis* spreads from an infected person (or via infected louse-infested clothes) to an uninfected person via infected human body lice and maybe head lice. Transmission from a louse to a person typically occurs when an infected louse is crushed by scratching, and coelomic fluid or louse feces is inoculated through broken skin or transferred by unclean hands to intact mucous membranes, e.g., conjunctiva, and possibly through the bite wound. No other animal hosts for *B. recurrentis* have been identified.

#### Disease

LBRF can be severe, with reported mortality rates today varying widely depending on social conditions and geography (untreated case mortality estimates range from 10-40% to 30-80%[38]). The 30-80% mortality estimates derive mainly from observational studies of specific outbreaks in Africa and are typically underpinned by little or poor diagnostic evidence. Moreover, other factors such as co-infections and/or malnutrition and starvation appear to play a key role in the outcome of LBRF in these studies[38]. Treatment reduces the case mortality rates to 2-5%. Past mortality rates are unknown and may have been variable due to genetic changes in *B. recurrentis* over time and/or in specific locations (as indicated by the mediaeval case in Norway described below), and, as today, would depend on social conditions, general health, and geography. Unidentified case numbers were, and are today, likely high and would reduce the estimated case mortality rates.

#### Evolution

The genus *Borrelia* comprises evolutionarily and genetically diverse bacterial species responsible for various diseases in humans and domestic animals[39]. These vector-borne spirochetes bifurcate into two principal evolutionary groups: the Lyme borreliosis clade and the relapsing fever clade, characterised by complex transmission cycles involving multiple host species and arthropod vectors.

When *Borrelia* colonises a novel vector species, it encounters unique selective pressures arising from distinct ecological interactions across disparate geographic ranges, leading to genetic divergence among populations vectored by different arthropod species[39]. As populations of the tick-borne relapsing fever species *B. duttonii* adapted to the human body louse they diverged from the ancestral *B. duttonii* populations in Africa, which are vectored by *O. moubata* (the African hut tampan or the eyeless tampan, a tick in the family Argasidae). Following this vector switch, the louse-borne populations experienced a massive geographic expansion due to their close association with human populations and hence human migrations. Genetic data have revealed sufficient evolutionary divergence and genome degradation to classify these human body louse-vectored bacterial populations as a distinct species, *B. recurrentis.*

A paper aiming to understand how a tick-borne pathogen adapts to the body louse sequenced and compared the genomes of the recurrent fever agents *Borrelia recurrentis* (LBRF) and *B. duttonii* (the cause of tick-borne recurrent fever, TBRF, prevalent in Sub-Saharan Africa) to their reconstructed ancestral genome[40]. Borrelia are unique among bacteria in that their genome consists of a linear chromosome and both linear and circular plasmids, and *B. recurrentis* and *B. duttonii* contain a unique 23-kb linear plasmid. This linear plasmid exhibits a large polyT track within the promoter region of an intact variable large protein gene and a telomere resolvase unique to Borrelia. The genome content is characterised by several repeat families, including antigenic lipoproteins (variable major proteins, vmp). Changes in these surface-exposed lipoproteins are likely implicated in immune escape. *B. recurrentis* exhibited a 20.4% genome size reduction, and it appeared to be a strain of *B. duttonii* with a decaying genome, possibly due to the accumulation of genomic errors induced by the loss of recA and mutS. Accompanying this were increases in impaired genes and a reduction in coding capacity, including surface-exposed lipoproteins and putative virulence factors. The results of comparative analyses of the reconstructed ancestral sequence and *B. duttonii* and *B. recurrentis,* respectively, was consistent with an accelerated evolution observed in *B. recurrentis*[40]*.* The correlation between gene loss and increased virulence of *B. recurrentis* parallels that of *Rickettsia prowazekii* (another bacteria transmitted through the human body louse and the cause of endemic typhus fever), with both species being genomic subsets of less-pathogenic strains[40].

A recent unpublished preprint presented phylogenetic analyses of four ancient British *B. recurrentis* genomes, which date back between 2,300 and 600 years ago, covering the Iron Age through to the later medieval period[41]. This study revealed the presence of at least four distinct clades over this time period. The first ancestral clade likely emerged after the split from *B. buttoni*, which was followed by the emergence of an Iron Age clade that appeared approximately 2,326 to 2,410 years ago. The timing of the Iron Age clade coincides with our observation of a transition from epidemic outbreaks to a state of endemicity of *B. recurrenti*s. This was followed by the emergence of a Medieval clade within the last 700 years and covers a period in time when we observed a continuous and relatively high steady level of *B. recurrentis* infections. Finally, a contemporary *B. recurrentis* clade emerged 46 to 69 years ago, which appears to be responsible for all recent LBRF infections. While ancient *B. recurrentis* diversity might be underestimated due to the scarcity of samples, contemporary diversity might also be underestimated because all modern *B. recurrentis* genomes are exclusively of African origin. This study also reconstructed a timeline of gene losses and additions using the pan-genome of related species; their analysis demonstrated that most of the reductive evolution observed in the *B. recurrentis* genome had already occurred around 2000 years ago, which might reflect rapid adaptation to the human louse vector and possibly also the human host. However, the acquisition and loss of plasmids continued at least until the Medieval time, e.g. pl26, pl27, and pl28 were lost sometime between the Iron Age and Medieval samples, while pl53 was only partially covered in the Iron Age samples but present in Medieval and present day samples. Similarly the presence of two ortholog genes implicated in plasmid segregation differed over time; *Soy-1* was found in *B. dottoni*, and in the Iron Age samples, but lost in medieval and contemporaneous samples, while *ParA-1* was only found in Medieval and present day samples. The latter pattern was also true for, e.g., the gene *oopA-1*. Likewise, the genes encoding the variable major proteins also appeared to change over time. The 5' end of many of these genes on pl33, pl37 and pl53 appeared to be present in some later Iron Age samples, but not the basal sample, while the 3' ends were absent in many Medieval samples, suggesting variability within *B. recurrentis* lineages. However, 3 of 4 Medieval vmps were similar to present day vmps, including two pseudogenes. The pathogenic implications of this variability require functional studies for clarification.

#### History and modern perspective

LBRF is an ancient epidemic disease often associated with war, famine, refugees, poverty, crowding and poor personal hygiene, but, as suggested by our findings, may well have been a more constant companion in the lifeways of past populations for thousands of years[36]. Today, outbreaks in endemic regions in Africa are more frequent during wet and cold seasons irrespective of other circumstances. These weather conditions promote prolonged stays indoors, possibly increase the likelihood of crowding, and increase the need for warm clothing that might be difficult to clean regularly, which increases the risk of body lice infestation and, thus, bacterial diseases transmitted by the louse vector. This pattern suggests that the Eurasian temperate climate with prolonged cold and often wet seasons, especially in the Northern parts, in the past could have contributed to the spread of LBRF even in peaceful and prosperous times.

LBRF can be identified in historical descriptions of disease epidemics by the repeated recurrences of fever between asymptomatic periods of 4–7 days and by two typical symptoms, jaundice and bleeding. The earliest convincing description of this disease was given by Hippocrates 2500 BP in Thassos, Greece: “The great majority (of sufferers) had a crisis on the sixth day, with an intermission of six days followed by a crisis on the fifth day after the relapse”[42]. Other typical features of LBRF were severe rigour, jaundice, profuse epistaxis (nosebleeds) and a tendency to precipitate abortion[36]. Arguments have been made that the ‘Yellow Plague’ that engulfed Europe in 550 AD (~1500 BP), following the Justinian plague, was caused by LBRF[36,43,44], a possibility which is supported by our data. An ancient genome of *B. recurrentis* was recovered from the skeleton of a young woman in Norway (AD 1430–1465, ~500 BP))[43,45]. This mediaeval genome displayed an ancestral oppA-1 gene, and gene loss in antigenic variation sites (variable short and long membrane protein genes) that translated into a genome reduction of 1.2% of the pan-genome, and 5.1–21% of the affected plasmids, perhaps associated with increased pathogenicity but a reduced number of relapses. The famine fevers of the 17th and 18th centuries in Ireland and elsewhere were likely predominantly LBRF epidemics[36]. LBRF was a common infection in Europe and North America until the end of the 19th century, after which it became a somewhat forgotten disease in these regions despite still causing severe epidemics elsewhere. In the 20th century, from 1903 to 1936, LBRF swept across North Africa, the Middle East and Africa, causing an estimated 50 million cases with 10% mortality[36]. A second epidemic in 1943–46 resulted in an estimated 10 million cases[36].

Today, LBRF mainly causes sporadic illness and outbreaks in regions of Africa, especially Ethiopia, Eritrea, and Somalia[36]. In the Ethiopian highlands, there are annual epidemics of thousands of cases coinciding with the rains, as the wet and cold weather promotes crowding. Moreover, it is the most frequently reported infection in Eritrean immigrants[46]. Since July 2015, LBRF has been diagnosed in almost 100 mainly young male refugees who arrived in several European countries, most in Italy and Germany, seeking asylum after travelling from Ethiopia, Eritrea, Somalia and other African countries, usually through Libya, where their disease progression profiles suggested many of these infections took place.

Notably, *B. recurrentis* retains the potential to cause future epidemics when conditions become conducive, and there are unmet needs for greater awareness, understanding and diagnostic ability of *B. recurrentis* and LBRF.
